# Mechanism and cellular actions of the potent AMPK inhibitor BAY-3827

**DOI:** 10.1101/2025.02.28.640688

**Authors:** Conchita Fraguas Bringas, Mohd Syed Ahangar, Joyceline Cuenco, Hongling Liu, Alex B. Addinsall, Maria Lindahl, Marc Foretz, Olga Göransson, John W. Scott, Elton Zeqiraj, Kei Sakamoto

**Affiliations:** Novo Nordisk Foundation Center for Basic Metabolic Research, Faculty of Health and Medical Sciences, University of Copenhagen, Copenhagen, Denmark; Astbury Centre for Structural Molecular Biology, School of Molecular and Cellular Biology, Faculty of Biological Sciences, University of Leeds, Leeds, United Kingdom; Department of Experimental Medical Science, Lund University, Lund, Sweden; Université Paris Cité, _CNRS_, _Inserm_, _Institut Cochin_, Paris, 75014, France; Drug Discovery Biology, Monash Institute of Pharmaceutical Sciences, Parkville, Melbourne, 3052, Australia; The Florey Institute of Neuroscience and Mental Health, Parkville, Melbourne, VIC 3052, Australia; St Vincent’s Institute of Medical Research, Fitzroy, Melbourne, VIC 3065, Australia

**Author notes:** These authors contributed equally to this work (Conchita Fraguas Bringas, Mohd Syed Ahangar).

## Abstract

Inhibition of AMP-activated protein kinase (AMPK) is under increasing investigation for its therapeutic potential in many diseases, including certain cancers. However, existing AMPK- inhibitors available as tool compounds are largely limited to compound C/dorsomorphin and SBI-0206965, both of which suffer from poor selectivity and off-target effects. Here we describe the structure-based molecular insights and cellular actions of a recently identified potent AMPK inhibitor, BAY-3827. Kinase selectivity profiling and sequence analyses of kinases that are highly or weakly inhibited by BAY-3827 uncovered key conserved residues involved in its inhibitory mechanism. A 2.5 Å co-crystal structure of the AMPK kinase domain (KD)-BAY-3827 complex and comparison with known KD-inhibitor structures, revealed an overlapping site in the ATP-binding pocket and an αC helix-out conformation. A distinct feature of the BAY-3827-bound state is the formation of a disulfide bridge between the αD helix Cys^106^ and the activation loop residue Cys^174^. This bridge appears to stabilize the activation loop such that Asn^162^ repositions the DFG motif Phe^158^ toward the C-terminal kinase lobe, displacing His^137^ and disrupting the regulatory spine, thereby promoting an inactive state. In hepatocytes, 2.5-5 μM BAY-3827, but not the structurally resembling inactive BAY-974, fully blocked AMPK activator (MK-8722)-mediated phosphorylation of ACC1 and corresponding inhibition of lipogenesis. Unbiased transcriptome analysis in MK- 8722-treated wild-type and AMPK-null hepatocytes revealed that 5 μM BAY-3827 downregulated >30% of MK-8722-stimulated AMPK-dependent genes. Based on its greater selectivity and potency substantiated by comprehensive structural and cellular investigations, BAY-3827 is a powerful tool to delineate AMPK functions.

**One-sentence summary:** We provide the mechanism of action of the potent and selective AMPK inhibitor BAY-3827, which blocks AMPK-dependent cellular functions.

## INTRODUCTION

AMP-activated protein kinase (AMPK) governs cellular energy metabolism and coordinates a myriad of cellular functions to maintain organismal homeostasis (*1–3*). AMPK is ubiquitously expressed in eukaryotic cells as heterotrimeric complexes comprising catalytic α and regulatory β and γ subunits. In mammals, the three subunits are encoded by multiple genes, giving rise to seven subunit isoforms (α1, α2; β1, β2; and γ1, γ2, and γ3) that can generate up to twelve distinct heterotrimeric complexes (*4,5*). AMPK is activated through phosphorylation of Thr^172^ within the activation segment of the α subunit kinase domain. The upstream kinases phosphorylating Thr^172^ are liver kinase B1 (LKB1) and Ca^2+^-calmodulin-dependent protein kinase kinase-2 (CaMKK2) (*1,2*). Thr^172^ is located in the activation loop of the catalytic α subunit, and the negative charge added upon residue phosphorylation leads to a conformational change that prepares AMPK for substrate phosphorylation acquiring an active kinase state. The Thr^172^-containing-activation loop is stabilized via charge-interactions with positively charged αC helix Lys^60^ and Arg^138^ and Asn^162^, swinging outwards to direct the γ- phosphate of ATP towards the peptide substrate. The conserved Asp-Phe-Gly (DFG) kinase motif will point away from the ATP-binding pocket, interacting with the conserved Lys^45^ residue and with ATP-Mg^2+^ interacting with Asp^157^ (*6,7*), allowing for phosphoryl transfer to take place. The γ-subunits contain four tandem cystathionine β-synthase (CBS) repeats that assemble into two Bateman domains, creating three functional nucleotide-binding sites that provide AMPK with its energy-sensing capabilities (*5*). The binding of ADP and/or AMP to CBS motifs causes conformational changes that increase net Thr^172^ phosphorylation. In addition, the binding of AMP, but not ADP, further increases AMPK activity through allosteric stimulation. AMPK activity is therefore controlled by multiple regulatory mechanisms, often involving a combination of post-translational modifications and allosteric regulation.

AMPK has long held promise as a therapeutic target for metabolic syndrome, as its physiological and pharmacological activation in multiple metabolic tissues has resulted in the amelioration of insulin resistance and reversal of hyperglycemia in preclinical studies (*8*). For example, active AMPK inhibits fatty acid and cholesterol biosynthesis in the liver through phosphorylation and inactivation of acetyl-CoA carboxylase-1 (ACC1) and HMG-CoA reductase (*9,10*). The activation of AMPK leads to increased fatty acid oxidation through phosphorylation of ACC2, and also promotion of plasma membrane localization of GLUT4 in part through phosphorylation of TBC1D1 via an insulin-independent mechanism and thereby glucose uptake in skeletal muscle (*11–14*). These beneficial effects of AMPK activation prompted the pharmaceutical development of small-molecule activators of AMPK, resulting in the identification of several potent and specific allosteric activators, including A- 769662 (*15,16*), 991/MK-8722 (*17,18*), and PF-739 (*19*). These compounds all bind into a pocket termed the Allosteric Drug and Metabolite (ADaM) site (*7*) located between the α and β subunits (*17, 8*). To date, some ADaM-site binding compounds have demonstrated promising efficacy in reversing hyperglycemia in pre-clinical proof of concept studies (*18, 19*). Even though current allosteric pan-AMPK activators have the potential risk of causing cardiac hypertrophy, isoform-selective activators are being developed to mitigate safety issues (*8*). Moreover, these highly potent and selective allosteric activators serve as valuable research tools and have profoundly contributed to delineate AMPK functions.

In cancer, AMPK has a dual role, as a tumor promoter or suppressor, depending on for instance subcellular context, trimeric complex formation and upstream regulators (*20, 21*). Previously, AMPK was viewed as tumor suppressor, where upon activation, it was reported to inhibit anabolic processes and mediate the tumor suppressive effects of LKB1 such as promoting cell growth through the inactivation of mTORC1 (*22, 23, 24*). Conversely, more recent studies have supported that the increased activity of AMPK may help to promote tumor growth and survival, alongside conferring drug resistance and resilience under tumor hypoxia and nutrient deprivation (*25,20*). In such conditions in established tumors, AMPK inhibitors might be efficacious tools for cancer treatment. Specifically, they may be particularly effective in cases where the *PRKAA1* or *PRKAB2* genes are amplified, causing aberrantly high expression of AMPK, and when given in combination with genotoxic treatments such as etoposide or radiotherapy, thus reducing the viability of tumor cells during such therapies (*21, 26*).

In contrast to activators, the availability of so-called “selective AMPK inhibitors” is limited and with major drawbacks. The first and most used AMPK inhibitor, the pyrazolopyrimidine derivative compound C, was identified in an in vitro high-throughput screen, and even though its selectivity was poorly characterized, it was employed to probe for AMPK-dependent metabolic actions of the anti-diabetes drug metformin in hepatocytes (*27*). Compound C was “re-discovered” in an in vivo compound screen in zebrafish embryos as an inhibitor of the bone morphogenetic protein (BMP) pathway (e.g., BMP type 1 receptors ALK2, 3 and 6) and was re-named “dorsomorphin” as it induced dorsalization in embryos (*28*). Co-crystal structure studies have established that compound C/dorsomorphin binds to the highly conserved ATP-binding pocket in the kinase domain (KD) of AMPKα (*29*) and ALK2 (*30*).

Multiple in vitro kinase activity screens studying inhibitor selectivity have verified that compound C is not a selective inhibitor of AMPK nor BMP receptor kinases (*31–33*), as for example it has been reported to inhibit the activities of 34 out of 119 kinases more potently than AMPK at 1 μM (*32*). In line with this, numerous cellular studies to date have shown that compound C affects a diverse range of biological processes via AMPK independent mechanisms (*34*).

SBI-0206965, a 2-aminopyrimidine derivative, was originally identified through screening of a library of pyrimidine analogues and described as an ATP-competitive inhibitor of the autophagy initiator kinase ULK1, with the ability to inhibit ULK signaling and ULK1- mediated survival of cancer cells (*35*). Recent studies have reported that SBI-0206965 inhibits AMPK and ULK1/2 with a similar potency and in a cell-free assay, SBI-0206965 inhibited AMPK with an *IC*_50_ of 0.16 μM at 20 μM ATP, compared with a value of 1.9 μM with compound C (*33*). Even though SBI-0206965 has demonstrated much higher potency and selectivity towards AMPK in cell-free assays compared to compound C, several other kinases including NUAK1, FAK, MLK1/3, and MARK3/4, all of which contain a methionine in their gatekeeper position in the kinase domain, are also potently inhibited (*36*). Structures of the AMPKα2(T172D) KD bound to SBI-0206965 (*33*) and compound C (*29*) show that SBI-0206965 contacts mainly the ATP-binding pocket, hinge and gatekeeper region. In cellular studies, SBI-0206965 dose-dependently blocked 991-induced phosphorylation of ACC1 and inhibition of lipogenesis in mouse primary hepatocytes (*36*). Notably, SBI- 0206965 also inhibited glucose and nucleoside transport systems in adipocytes and myotubes.

More recently, a potent and selective AMPK inhibitor termed BAY-3827 was developed in a high throughput compound screen and underwent chemical optimization. In cell-free assays, BAY-3827 inhibited AMPK with *IC*_50_ values of 1.4 nM at low (10 μM) ATP, and 15 nM at high (2 mM) ATP concentrations (with 2 μM AMP under both conditions), although it inhibited 90 kDa ribosomal S6 kinase (RSK) isoforms at a similar potency. In cellular assays, BAY-3827 inhibited phosphorylation of ACC1 and showed anti-proliferative effects in a subset of tumor models, namely androgen-dependent prostate cancer and myeloma cell lines (*37*). However, it is unknown if this anti-proliferative effect of BAY-3827 is mediated through AMPK. To date, little or no information about the molecular mechanism of AMPK inhibition by BAY-3827 has been demonstrated. Here, we describe the molecular basis for BAY-3827’s inhibitory effect on AMPK through a co-crystal structure of the AMPKα2-KD with BAY-3827. In addition, we provide cellular potency and selectivity data of BAY-3827 with a specific emphasis on AMPK-dependent transcriptional and metabolic processes (fatty acid synthesis) in wild-type (WT) and AMPKα1α2 -null primary mouse hepatocytes.

## RESULTS

### Kinase inhibition selectivity profile of BAY-3827 and its structurally related BAY-974

We determined the selectivity of BAY-3827 alongside its inactive control compound BAY- 974, which shares a similar chemical scaffold with BAY-3827 by performing an in vitro kinase activity assay across a panel of 140 human protein kinases (**Fig. 1A, B**). Both compounds were tested at 0.1 μM in an assay and consistent with previous data, AMPK was the most inhibited (>90%) by BAY-3827 followed by PHK and RSK1 (∼80-85%) (**Fig. 1A, D**), with RSK1 reported previously to be among the top inhibited kinases by BAY-3827 at 1 μM in a different panel tested at 10 μM ATP (*37*). Aurora B, MINK1, TBK1 and IRAK1 were also inhibited (∼60 %) by BAY-3827 (**Fig. 1A, D**). The inhibitory profile of BAY-3827 in the tested human kinome (*38*) displayed a preference for the CAMK group (which includes AMPK and PHK) and the AGC group containing RSK family and MSK1. Furthermore, Aur family Aurora B and STE group kinases MINK1 and MST3 were also highly inhibited alongside TKL group belonging IRAK1. As anticipated, BAY-974 had almost no effect on AMPK, however it inhibited (∼55-90%) other kinases mainly from the TK and TKL groups, including SYK, YES1, IRAK1 and ULK1 from the ULK family (**Fig. 1Bg3., Table S1**). These results demonstrate that BAY-3827, but not its structurally related BAY-974, selectively inhibits AMPK. This can be attributed to specific chemical alterations, including a missing fluoride group in the indazole ring which is present in BAY-3827, a methyl group in the benzamide ring instead of an ethyl group and finally a dimethyl dihydropyridine ring compared to a trimethyl pyridine ring as seen in BAY-3827 (**Fig. 1C**). The inhibitory profiles between BAY-3827 and BAY-974 across this panel of kinases with ≤ 50% remaining activity revealed a minimal overlap (**Fig. 1D**), making BAY-974 a suitable control compound to help dissect the BAY-3827 specific effects.

**Figure 1.**
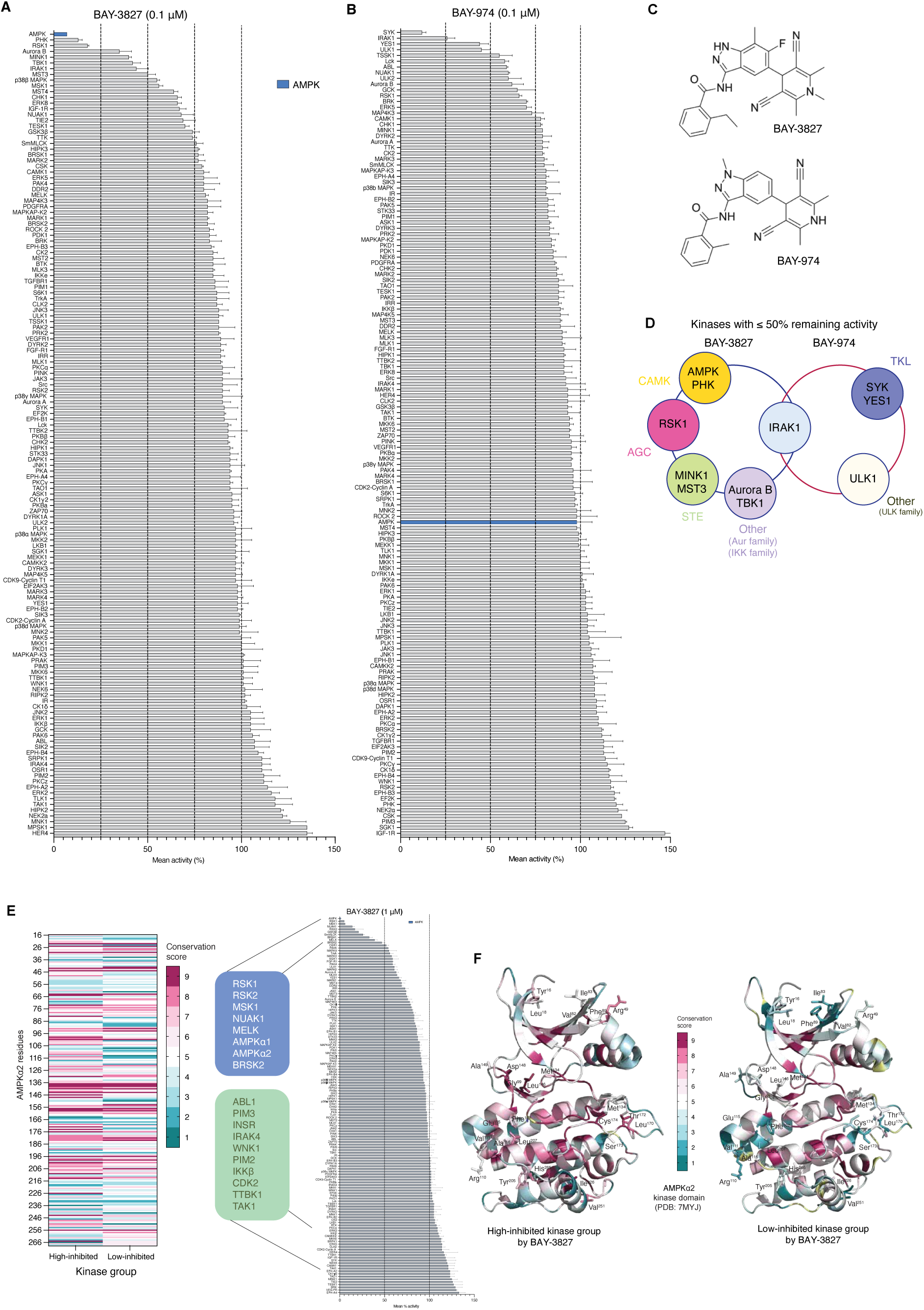
BAY-3827 but not BAY-974 selectively inhibits AMPK. Kinase activity assays were conducted across a panel of 140 human kinases and compound selectivity of **A)** BAY-3827 (0.1 μM) and **B)** BAY-974 (0.1 μM) was assessed, reporting remaining kinase activity (%) from n=2 in mean ± SEM. **C)** Chemical structures of BAY-3827 (*N*-[5-(3,5-dicyano-1,2,6-trimethyl-4*H*-pyridin-4-yl)-6- fluoro-7-methyl-1*H*-indazol-3-yl]-2-ethylbenzamide) and BAY-974 (*N*-[5-(3,5-dicyano-2,6-dimethyl- 1,4-dihydropyridin-4-yl)-1-methylindazol-3-yl]-2-methylbenzamide) drawn in Chemaxon Marvin (24.3.163). **D)** Schematic showing the inhibitory overlap between BAY-3827 and BAY-974 of kinases with ≤50% remaining activity as well as the corresponding kinase groups they belong to. **E)** In silico residue conservation analysis based on high-inhibited (≤50% remaining activity) and low-inhibited (≥99% remaining activity) kinases by BAY-3827 at 1 μM across a 140-human kinase panel cross- referenced with available data (*37*). **F)** Ribbon view of AMPKα2 kinase domain (PDB: 7MYJ, chain A) showing computed ConSurf conservation scale (1–9) colours (*39,40*) between high-inhibited kinase group (left) and low-inhibited kinase group (right) depicting the residues with a 3-point or higher scale difference between groups.

Achieving a specific kinase inhibitor often involves targeting residues which are unique to the target kinase or lowly conserved across the kinome. Based on the selectivity profile of BAY-3827, we attempted to identify the pattern of this compound’s preference by studying residue conservation in the kinase domain of highly inhibited kinases compared to those that were not inhibited by BAY-3827. To do this comprehensively, we screened a higher concentration of BAY-3827 (1 μM) across the same kinase panel, and this data was cross-referenced with available kinase screen data at the same concentration (*37*) (**Fig. 1E**). We defined a high-inhibited kinase group as those kinases which had 50% or less activity remaining after BAY-3827 treatment, and a low-inhibited kinase group representing those with 99% or more remaining activity after BAY-3827 treatment. AMPK, BRSK2, MELK, NUAK1, MSK1, RSK1 and RSK2 composed the high-inhibited group (**Fig. 1E**, shown in blue) and kinases ABL1, PIM3, INSR, IRAK4, WNK1, PIM2, IKKβ, CDK2, TTBK1 and

TAK1 were selected as representatives of the low-inhibited group (**Fig. 1E**, shown in green). A sequence alignment of human kinase domains was conducted in each group and their conservation was studied using AMPKα1 as a reference sequence, with the high-inhibited group having 30% or higher percentage sequence identity up to 47.8% with BRSK2, whereas the low-inhibited group ranged from 29.7% to 24.1% (**Table S2**). Based on residue conservation a score was computed using the ConSurf server (*39, 40*) (**Fig. 1E, F, Table S2**), revealing strikingly different residue conservation patterns between the two kinase groups (**Fig. S1 A, B**). We therefore hypothesized that highly conserved residues in the high- inhibited, but not in the low-inhibited kinase group, would be important for BAY-3827’s inhibitory mechanism. Residues with at least a three point-difference in the conservation scale (1–9) between kinase groups were displayed in the structure of AMPKα2. These were localised in four clusters: (1) in the N-terminal lobe such as Tyr^16^ and Arg^49^, near the ATP-binding pocket, including residues such as Gly^99^, Leu^146^ and Asp^148^, (2) in the C-terminal lobe helices such as His^265^ and Val^251^ and (3) in the activation loop, where Thr^172^ had a 4-point scale decrease in conservation in the low-inhibited group, and finally (4) Cys^174^ and Leu^170^ had a striking 6-point decrease compared to the high-inhibited kinase group (**Fig.1E, Table S2**). Collectively, these analyses suggest that there is a conservation basis for BAY-3827’s mechanism of inhibitory action that involves residues around the ATP-binding pocket and the activation segment.

### AMPKα2 binds BAY-3827 in a DFG-in inactive kinase conformation

To understand the precise molecular mechanism behind BAY-3827’s inhibition of AMPK, the co-crystal structure of BAY-3827 with wild type (WT) AMPKα2 kinase domain (KD) was obtained at a resolution of 2.5 Å (Table 1, **Fig.2A**). BAY-3827 binds into the conserved ATP-binding pocket between the N- and C-terminal lobes, as revealed by an unambiguous compound density map (**Fig.2B**), binding with a *K_d_* value of 1.23 μM as measured by a spectral shift binding assay (**Fig.2A, S1C**). In this structure, the activation loop (highlighted in magenta) instead of swinging out in an active conformation (**Fig.S1D**) (*41*) hovers over the kinase core occupying a space below the β1 sheet and interacts with the αD helix in the C-lobe through Cys^174^, constituting an inactive kinase conformation (**Fig. 2A**). The indazole ring in BAY-3827 hydrogen bonds with hinge region residues, with a single bond formed with Glu^94^ and two with Val^96^ alongside two additional hydrogen bonds are made with N-lobe residues Leu^22^ and Lys^45^ (**Fig. 2C, S1E**). Leu^22^ bonding to BAY-3827’s benzamide moiety is water-mediated and the conserved Lys^45^ in the β3 sheet interacts with the pyridine ring (**Fig. 2C**). Since BAY-3827 interacts directly with Lys^45^, the typical salt bridge with Glu^64^ in the αC helix is no longer possible, and with a Lys-Glu distance of 11.7 Å (**Table 2**), the αC helix has an out- conformation (*42*). Residues which accommodate and engage in hydrophobic contacts with BAY-3827 include hinge region residues Tyr^95^ and Ser^97^, as well as N-lobe residues Gly^99^, Val^30^ and Asp^166^. In the C-lobe, residues Glu^100^, Gly^23^, Ile^77^ and Ala^43^ and the gatekeeper Met^93^ also contribute to ligand hydrophobic contacts, as well as Leu^146^, which faces BAY-3827’s benzamine moiety. Moreover, activation loop residues Asn^144^ and Ala^156^, alongside Asn^162^, Met^164^ and Ser^165^ face towards BAY-3827’s pyridine ring and engage in hydrophobic contacts (**Fig. S1E**), with BAY-3827 occupying a space in the ATP-binding pocket which does not extend further past gatekeeper Met^93^. Focusing on the DFG motif, Asp^157^ presents with an out- conformation and DFG motif Phe^158^ residue has shifted down towards the C-lobe compared to the kinase active state in AMPK (**Fig.S1D**) (*41*). In the C-lobe, a key interaction occurs between αD helix Cys^106^ and activation loop residue Cys^174^ which form a disulfide bridge (**Fig. 2D**). Superimposition of AMPKα2KD-BAY-3827 with available compound C and SBI-0206965 structures (*29, 33*) showed a partial overlap with compound C’s binding site in the ATP-binding pocket, however the extension of BAY-3827 binding was shared with SBI-0206965’s binding site, whereas compound C further extends outside the pocket (**Fig. 2E, S1F**).

**Figure 2.**
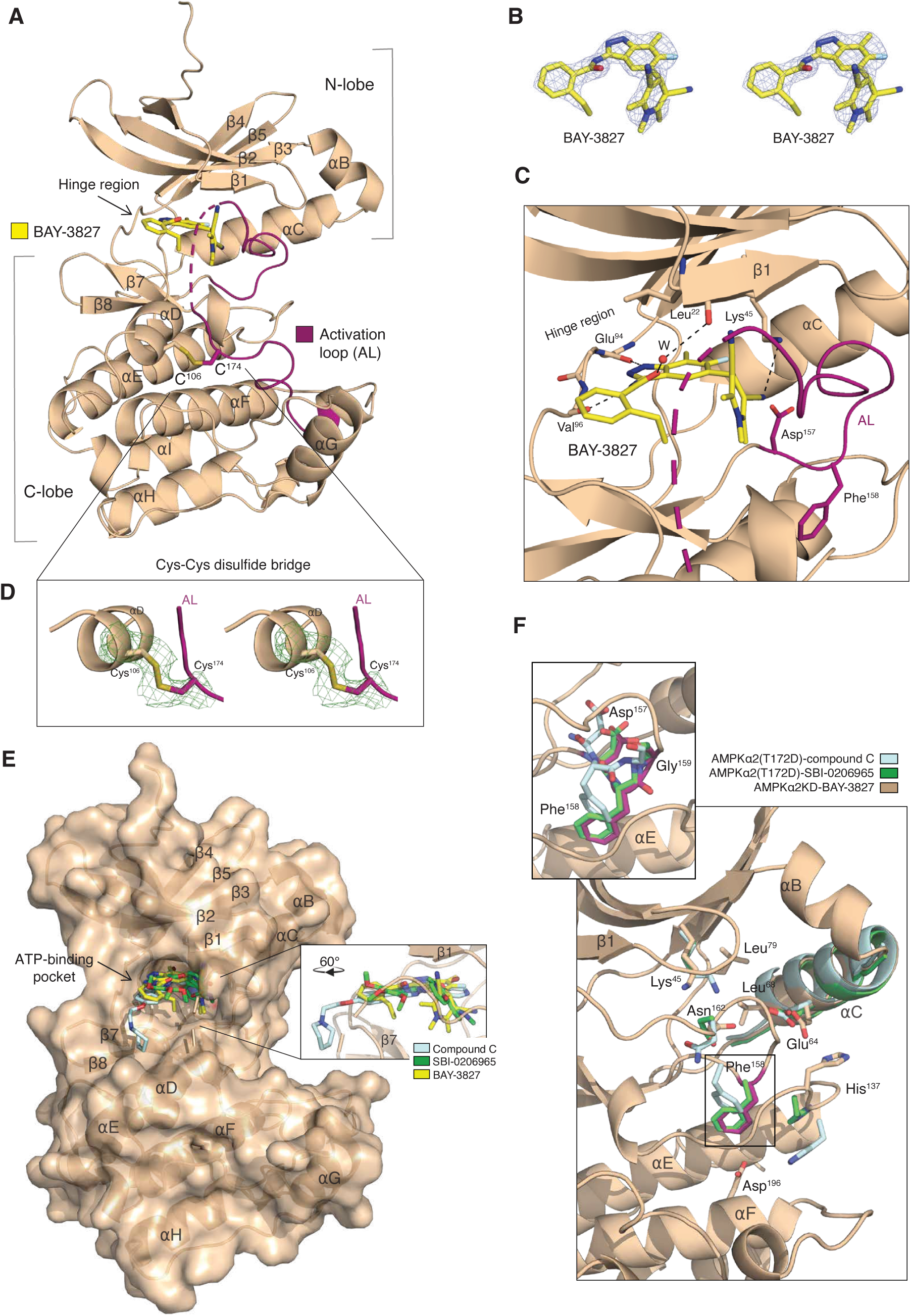
Structure of AMPKα2 kinase domain bound to BAY-3827. **A)** Co-crystal structure of AMPKα2 kinase domain (shown as ribbons) with BAY-3827 (shown as yellow sticks). The activation loop (AL) segment is coloured red. **B)** Stereo-image of the 2Fo-Fc omit map of BAY-3827 shown at 1σ. **C)** Magnified view of the BAY-3827 binding site with interactions indicated as grey dash lines. The side chains of Asp^157^ and Phe^158^ of the DFG motif are shown as sticks with Phe^158^ in an “FG down” conformation. Water molecule (W) is shown as a red sphere. A disulfide bond (Cys^106^-Cys^174^) is shown as sticks. **D)** Stereo image of the Fo-Fc polder map (Phenix) of the indicated disulfide bond between Cys^106^ from αD and Cys^174^ from activation loop (AL). **E)** Surface view of AMPKα2 kinase domain bound to BAY-3827 with superimposed compound C (light cyan, PDB:3AQV) and SBI-0206965 green, PDB:6BX6) inhibitors shown with sticks (*29, 33,77*). **F)** Zoomed-in view of the DFG motif superimposed with compound C (light cyan, PDB: 3AQV) and SBI-0206965 (green, PDB:6BX6), displaying the backbone (as ribbons) of AMPKα2 kinase domain (KD) bound to BAY- 3827 showing key regulatory spine residues.

**Table 1.** Crystallography data collection and refinement statistics. Crystallography data collection for AMPK⍺2 kinase domain (KD) with BAY-3827. Statistics for the highest- resolution shell are shown in parentheses.

| | α2KD-BAY3827 |
| --- | --- |
| Wavelength | 0.6199 Å |
| Resolution range | 58.89 - 2.5 (2.56 - 2.50) |
| Space group | P 21 21 2 |
| Unit cell | 59.176 117.778 38.327 90 90 90 |
| Total reflections | 65446 (6013) |
| Unique reflections | 10021 (890) |
| Multiplicity | 6.5 (6.8) |
| Completeness (%) | 99.9 (100.00) |
| Mean I/sigma(I) | 15.70 (4.6) |
| Wilson B-factor | 39.04 |
| R-merge | 0.074 (0.418) |
| R-meas | 0.081 (0.454) |
| R-pim | 0.0332 (0.174) |
| CC1/2 | 0.997 (0.944) |
| CC* | 0.999 (0.973) |
| Reflections used in refinement | 9799 (2397) |
| Reflections used for R-free | 497 (119) |
| R-work | 0.2049 (0.2606) |
| R-free | 0.2327 (0.3047) |
| Number of non-hydrogen atoms | 2256 |
| macromolecules | 2142 |
| ligands | 43 |
| solvent | 71 |
| Protein residues | 267 |
| RMS (bonds) | 0.002 |
| RMS (angles) | 0.52 |
| Ramachandran favoured (%) | 95.44 |
| Ramachandran allowed (%) | 3.04 |
| Ramachandran outliers (%) | 1.52 |
| Rotamer outliers (%) | 6.06 |
| Clash score | 5.52 |
| Average B-factor | 53.50 |
| macromolecules | 53.90 |
| ligands | 48.62 |
| solvent | 44.43 |

**Table 2.** Atomic distance measurements to define kinase conformation. Distance measurements performed as previously described (*42*).

| Structures | **D1: Leu<sup>68</sup>.CA-Phe<sup>158</sup>.A CZ** | **D2: Lys<sup>45</sup>.CA-Phe<sup>158</sup>.CZ** | **Lys<sup>45</sup>.CB-Glu<sup>64</sup>.CB** |
| --- | --- | --- | --- |
| AMPKα2-KD BAY-3827 | 12.4 Å | 18.4 Å | 11.7 Å |
| AMPKα2 (T172D)-KD-compound C (PDB: 3AQV) | 11.4 Å | 16.6 Å | 11.3 Å |
| AMPKα2 (T172D)-KD-SBI-0206965 (PDB: 6BX6) | 13.1 Å | 18.2 Å | 11.8 Å |

Recent classification of kinase structural states based on DFG motif conformation used the positions of the αC helix and the DFG Phe^158^ ring relative to Lys^45^ (D1 and D2 distances) (*42*). According to these measurements (**Table 2**), AMPKα2KD–BAY-3827, as well as the AMPKα2(T172D)–inhibitor structures of compound C and SBI-0206965 clustered together with other reported “DFG-in” structures (*42*). To further classify the DFG conformation, we measured Φ/Ψ angles of x-DFG-x motif residues of our AMPKα2KD–BAY-3827 structure (**Table 3**), however we could not place the structure in an established cluster, suggesting a DFG-in conformation in an inactive kinase state which has not yet been defined (*42*). Interestingly, SBI-0206965 (*33*) and BAY-3827–bound structures showed nearly identical angle measurements (**Table 3**) and a “Phe158-down” orientation toward the C-lobe compared to the active AMPK conformation (**Fig. 2F, S1 D, F**). In AMPKα2(T172D)–compound C (*29*), Phe^158^ also pointed down toward the C-lobe but adopted a different conformation than in BAY-3827/SBI-0206965 structures (**Fig. 2F**, **Table 3**).

**Table 3.** DFG motif Phe^158^ dihedral angle (Phi,Psi) measurements. Measurements were performed in PyMOL (*77*).

| Structures | **Ala<sup>156</sup>**<br>**($\Phi$, $\psi$)** | **Asp<sup>157</sup>**<br>**($\Phi$, $\psi$)** | **Phe<sup>158</sup>**<br>**($\Phi$, $\psi$)** | **Gly<sup>159</sup>**<br>**($\Phi$, $\psi$)** | **Leu<sup>160</sup>**<br>**($\Phi$, $\psi$)** |
| --- | --- | --- | --- | --- | --- |
| AMPKα2-KD BAY-3827 | -77.8°, 133.2° | -64.3°, 162.0° | -90.6°, 42.6° | -91.5°, 174.0° | -69.5°, -13.6° |
| AMPKα2 (T172D)-KD compound C (PDB: 3AQV) | -77.6°, 137.5° | -58.4°, 143.4° | -127.8°, 21.4° | -78.7°, -145.7° | -61.1°, -34.9° |
| AMPKα2 (T172D)-KD-SBI-0206965 (PDB: 6BX6) | -83.3°, 141.7° | -56.9°, 150.6° | -90.9°, 68.7° | -125.6°, 179.3° | -68.2°, -25.5° |

In AMPKα2KD–BAY-3827, activation-loop residue Asn^162^ occupies a position below Lys^45^ in the N-lobe, stabilized by hydrogen bonds with DFG residues Asp^157^ and Gly^159^ (**Fig. 2F**). This differs from its proximity to His^137^ in the C-lobe observed in active AMPK structures (**Fig. S1D**) (*41*). As a result, Phe^158^ is displaced downward toward the αE helix, and His^137^ is pushed outward—away from its typical stack on Asp^196^—resulting in a broken hydrophobic spine characteristic of an inactive kinase (**Fig. 2F, S1G**) (*44*). Notably, in BAY-3827–bound AMPKα2, His^137^ points toward the αC helix in the N-lobe, whereas in SBI-0206965– and compound C–bound structures it faces the C-lobe (**Fig. S1G, 2F**), highlighting different ways of breaking the regulatory spine. Superimposing these three inhibitors also revealed a preserved αC helix conformation (**Fig. 2F, S1D, F**). Overall, given their overlapping binding sites and DFG conformations (**Fig. 2E, F**), the “FG-down” arrangement may represent a pivotal DFG-in transitional state in AMPK structural dynamics that is exploited by small molecule inhibitors.

### BAY-3827 inhibits different AMPK trimeric complexes at similar potency

Given the diverse assemblies that AMPK complexes can take depending on their subunit isoforms, BAY-3827’s inhibitory action alongside BAY-974 control was studied using cell-free assays in different AMPK complexes, namely α2β1γ1, α1β1γ1 and α2β2γ1 (**Fig.3 A-E**). The α1β1γ1 and α2β2γ1 trimeric complexes showed similar *IC*_50_ values when tested at high (200 μM) ATP concentrations, with *IC*_50_ values of 31.2 nM and 36.5 nM respectively, whereas the α2β1γ1 had a moderately higher *IC*_50_ of 57 nM. Under lower ATP condition (20 μM), α1β1γ1 and α2β2γ1 had values of 1.8 nM and 2.8 nM, with the α2β1γ1 complex having an *IC*_50_ value of 4.1 nM (**Fig. 3E)**. Despite this modest difference, BAY-3827 has an inhibitory effect in the same order of magnitude and kinetics across all three tested AMPK complexes (**Fig. 3A-C**), whereas BAY-974 showed no effect (**Fig. 3D**). We also tested the effects of BAY-3827 and BAY-974 at increasing concentrations on selected kinases; ULK1 (known to be potently inhibited by SBI-0206965 (*36*)) and CaMKK2 (upstream kinase of AMPK) and confirmed no inhibitory effect (**Fig. S2A, B**).

**Figure 3.**
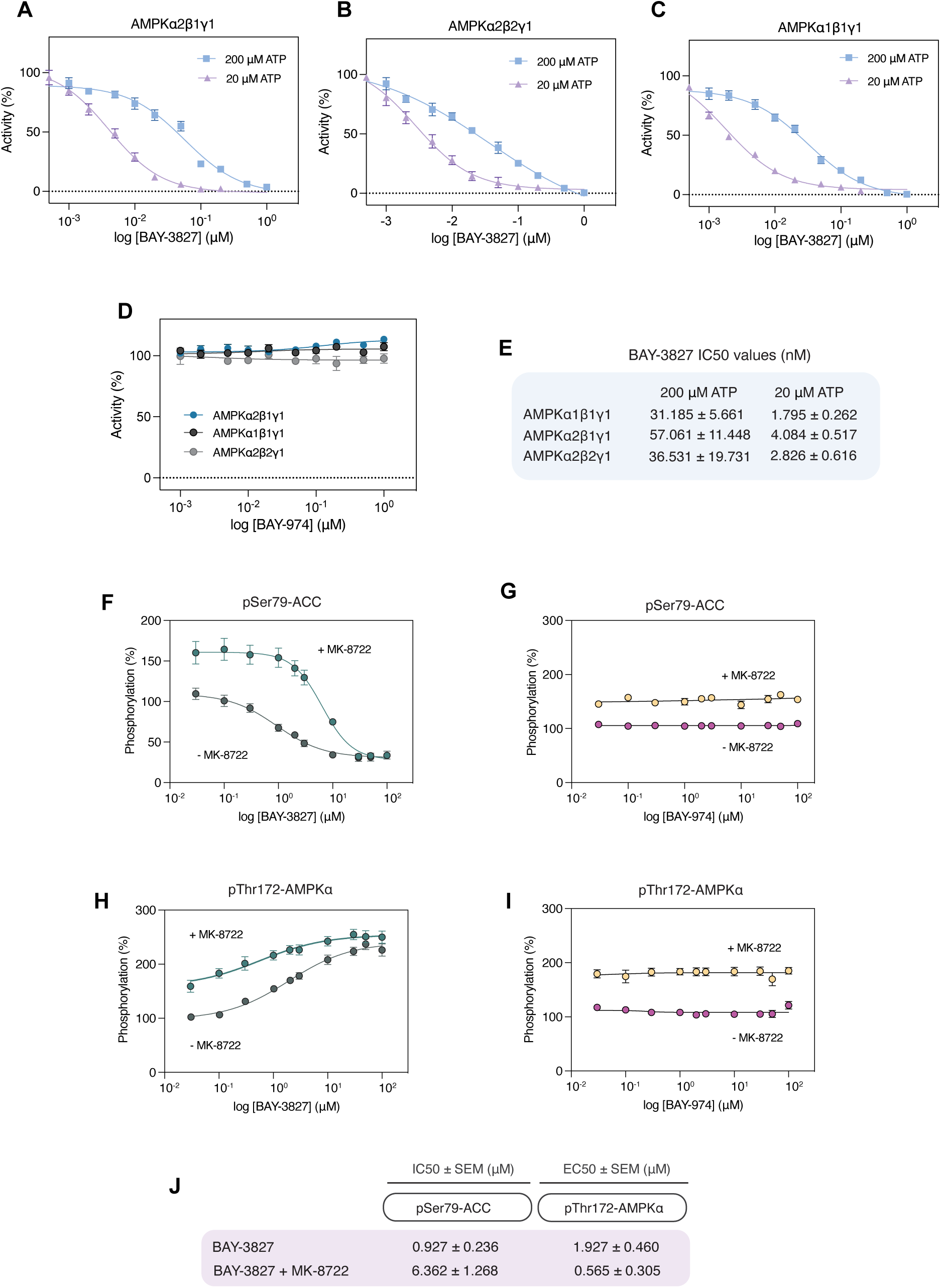
BAY-3827 and BAY-974 kinetics. A)-. **C)** Inhibition of AMPK complexes in cell free assays by BAY-3827 in high (200 μM) and low (20 μM) ATP concentrations and by **D)** BAY-974 at 200 μM. Data are n=3 represented as mean ± SEM. **E)** *IC*_50_ values (μM) of BAY-3827 inhibition of AMPK complexes in different ATP conditions. **F)-G)** HTRF assays in U2OS cells measuring % phosphorylation of Ser79-ACC following BAY-3827 or BAY-974 treatment and % phosphorylation of **H)-I)** Thr172-AMPK in the presence or absence of AMPK activator MK-8722 (10 μM). **J)** Estimated *IC*_50_ and *EC*_50_ values shown respectively from F)-I). Data is mean ± SEM with n=3 from three independent experiments.

### Cellular kinetics analysis of AMPK by BAY-3827

Prior to using BAY-3827 alongside BAY-974 control in cellular functional studies, their kinetics on AMPK activity were investigated in U2OS cells, an osteosarcoma cell line. Homogeneous time-resolved fluorescence (HTRF) is a high-throughput fluorescence resonance energy transfer (FRET)-based technique previously benchmarked as a sensitive and quantitative assay for detection of ACC1 Ser79 phosphorylation (*36, 45*) and is an established surrogate marker for cellular AMPK activation. HTRF assays were performed following treatment with BAY-3827 and BAY-974 in the presence and absence of allosteric AMPK activator MK-8722 (*18*), with vehicle compared to MK-8722 control treatment showing an expected significant increase in both pACC1 and pAMPK (**Fig. S2C, D**). ACC1 phosphorylation dose-dependently decreased upon BAY-3827 treatment, with an *IC*_50_ of 0.93 μM, which increased to a value of 6.36 μM in cells co-incubated with MK-8722. No inhibition was observed with BAY-974 even at the highest tested concentration of 100 μM (**Fig. 3F-G, J**). AMPK phosphorylation dose-dependently increased with BAY-3827, but not BAY-974, with an *EC50* of 1.93 μM and 0.57 μM in the presence of MK-8722 (**Fig. 3H-J**).

This observed increase in pAMPK is consistent with the reported effects of BAY-3827 on AMPK phosphorylation levels possibly due to its protective effect against Thr^172^ dephosphorylation (*46*).

### BAY-3827 inhibits AMPK and blocks its suppression effect on lipogenesis in primary hepatocytes

We next sought to establish if BAY-3827 inhibited an AMPK-dependent process in a more physiologically relevant cell system. Mouse primary hepatocytes were treated with increasing doses of BAY-3827 with or without MK-8722 (10 μM). Immunoblot analysis revealed a dose-dependent reduction in phosphorylation of ACC1 and Raptor, with full inhibition in the presence of MK-8722 achieved at 5 and 2.5 μM in pACC and pRaptor, respectively (**Fig. 4A- D**). Increased pThr^172^AMPK levels were only significant under MK-8722-treated conditions at 5 μM (**Fig. 4E**). BAY-974 (5 μM) showed no effect on levels of pSer^79^ACC1, pRaptor and pAMPK with or without MK-8722 (**Fig. 4C-E**).

**Figure 4.**
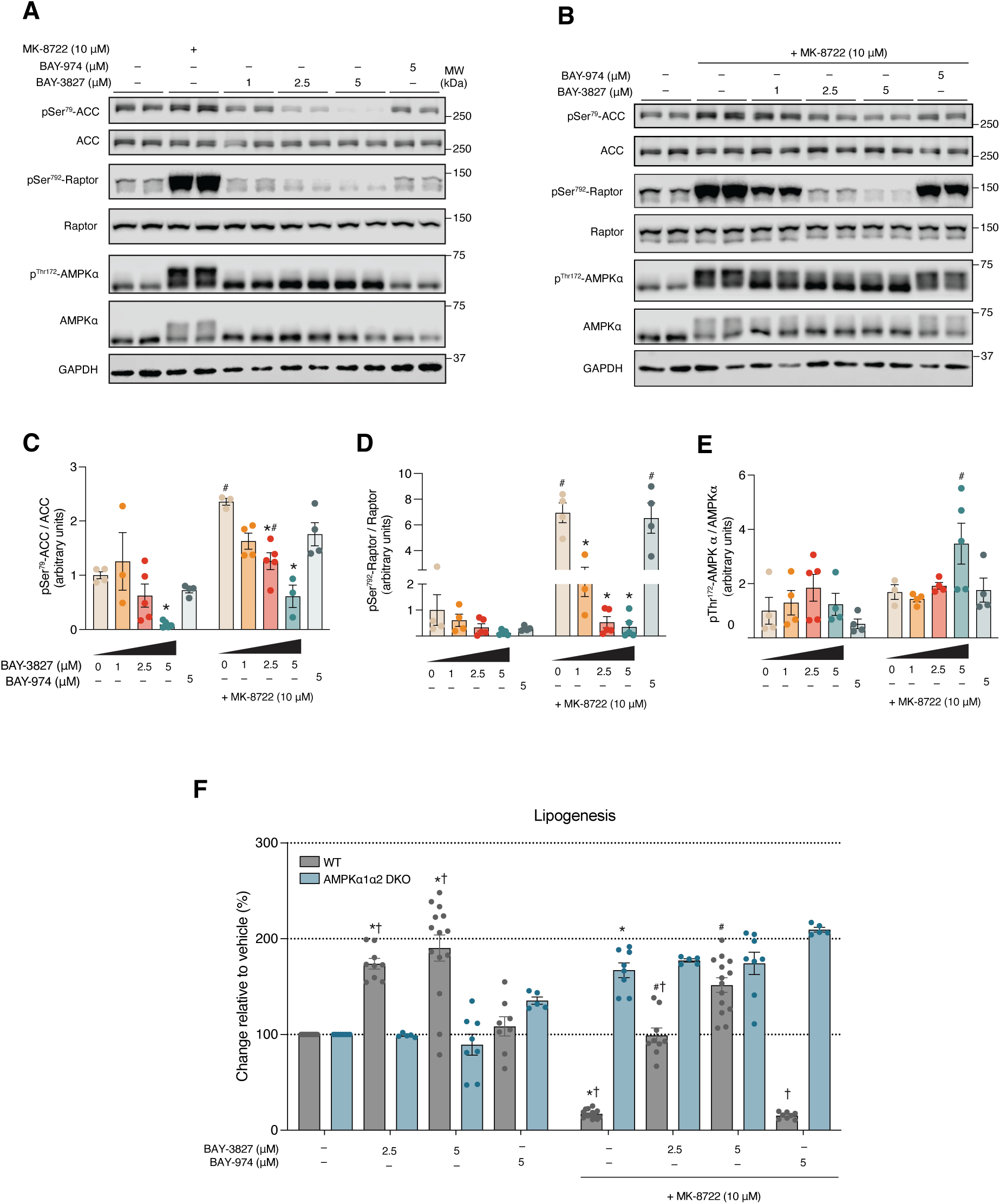
BAY-3827 but not BAY-974 dose-dependently reverse AMPK-inhibited lipogenesis in primary hepatocytes. **A)** Representative blots of wild-type hepatocytes treated with BAY-3827 and BAY-974 in basal conditions and **B)** treated with 10 μM MK-8722 after compound treatment for 15 min, with selected antibodies describing AMPK signalling. **C)-E)** Calculated phosphorylation ratios following band intensity quantification of data from (A, B) normalised to loading control showing mean ± SEM from four separate experiments with n=2-3 biological replicates. **F)** Lipogenesis assay of primary and AMPKα1 α2 double knockout (DKO) primary hepatocytes treated with BAY-3827 and inactive BAY-974, where n=3-5 from five separate experiments. *p <0.05 (vehicle vs indicated treatment), ^#^p <0.05 (vehicle + MK-8722 vs in indicated treatment) and ^†^p < 0.05 (WT vs KO treatments in basal and + MK-8722 conditions).

One of the best-characterized physiological consequences of AMPK activation is the suppression of hepatic lipogenesis through phosphorylation of Ser^79^ on ACC1 (*47, 48*). Mouse primary hepatocytes derived from WT and liver-specific AMPKα1α2 double knockout (DKO) mice were treated with BAY-3827 (2.5 and 5 μM) with or without MK-8722 (10 μM), and [^14^C]-acetate incorporation into fatty acids was assessed. Basal lipogenesis (without MK- 8722) was significantly higher in WT compared to AMPKα1α2 DKO hepatocytes at 2.5 and 5 μM treatments, with 50% and 70% increases respectively. As anticipated, MK-8722 robustly suppressed (>80%) lipogenesis in WT, but had no effect in AMPKα1α2 DKO cells. In WT, BAY-3827 at both doses significantly increased lipogenesis in the absence of MK- 8722 and restored the inhibitory effect of MK-8722. In contrast, BAY-3827 had no effect on lipogenesis in AMPKα1α2 DKO hepatocytes (**Fig. 4F**). BAY-974 at 5 μM treatment had no effect on lipogenesis under basal and MK-8722-treated conditions. We also tested if compound C was able to rescue MK-8722-mediated inhibition of hepatic lipogenesis and observed that it only negligibly (10%) restored at 25 μM, confirming its low potency in cells (**Fig. S2E**).

We have previously reported that SBI-0206965 not only inhibits AMPK-related kinases (NUAK1, MARK3/4) equally or more potently than AMPK or ULK1, but also critical cellular functions such as basal and insulin-stimulated glucose uptake in adipocytes and skeletal muscle (*36*). We treated mouse primary adipocytes and isolated mouse skeletal muscle ex vivo with varying doses of BAY-3827 with or without MK-8722 or insulin (**Fig. S2F-J**). We observed a dose dependent reduction in Raptor phosphorylation in response to BAY-3827 irrespective of MK-8722’s presence (**Fig.S2F, G**). Both basal and insulin-stimulated phosphorylation of Akt and TBC1D4, as well as glucose uptake were not significantly affected in adipocytes (**Fig.S2H-J**). In ex vivo skeletal muscle, BAY-3827 dose-dependently inhibited MK-8722-stimulated glucose uptake and a full inhibition was achieved at 5 μM (**Fig.S2K**). BAY-3827 attenuated insulin-stimulated (AMPK-independent) glucose uptake in ex vivo skeletal muscle, indicating a potential off-target effect.

### BAY-3827 diminishes drug-stimulated AMPK-dependent gene expression

To further examine BAY-3827’s specific effect on AMPK signaling and gene expression in an unbiased manner, we treated WT and AMPKα1α2 DKO mouse primary hepatocytes with 5 μM BAY-3827 in the presence or absence of MK-8722 and performed RNA-Seq analysis. We focused on 1841 MK-8722-stimulated genes that were significantly up-regulated compared to basal conditions in WT cells (**Fig. 5**). To confidently define them as AMPK-dependent genes, we filtered them with a list of significantly downregulated genes under AMPKα1α2-null, MK-8722-treated conditions which yielded 1973 genes. Among them, 845 genes were both MK-8722-stimulated and AMPK-dependent, and their expression was displayed as a heatmap (**Fig. 5A**). Upregulated gene expression in MK-8722-treated WT cells was almost completely abolished in AMPKα1α2 DKO cells (**Fig. 5A**). In hepatocytes treated with BAY-3827 and MK-8722, approximately 30% of MK-stimulated-AMPK genes were downregulated (524 genes out of 845 predefined MK-8722-AMPK genes) (**Fig. 5A, B, Fig. S3A**). In WT hepatocytes treated with BAY-3827 and MK-8722, 2,511 significant genes were differentially expressed compared to MK-8722, with 61.2% (1,539 genes) downregulated by BAY-3827 treatment when AMPK is active, with top significant (FC < -1.3, FDR < 0.05) downregulated genes including *PLCE1*, *ZFP36*, *FIGNL2*, *ADAMTSL4* and *TIMP3* (**Fig. 5C**). In AMPKα1α2 DKO cells compared to WT treated with MK-8722, *ADAMTSL4* and *ZFP36* were also amongst the top downregulated genes (**Fig. S3B**). Gene ontology enrichment analysis of biological processes was performed to query the functions of the downregulated genes where AMPK is inactivated either via genetic (AMPKα1α2 DKO) or pharmacological (BAY-3827) means in the presence of MK-8722 (**Fig. 5D, E**). The top 25 categories (FDR < 0.05) of downregulated genes in these two conditions showed a large overlap in known AMPK-regulated functions which BAY-3827 also inhibited, such as lipid oxidation and modulation, organic acid catabolic processes and response to nutrient levels and starvation (**Fig. 5F**).

**Figure 5.**
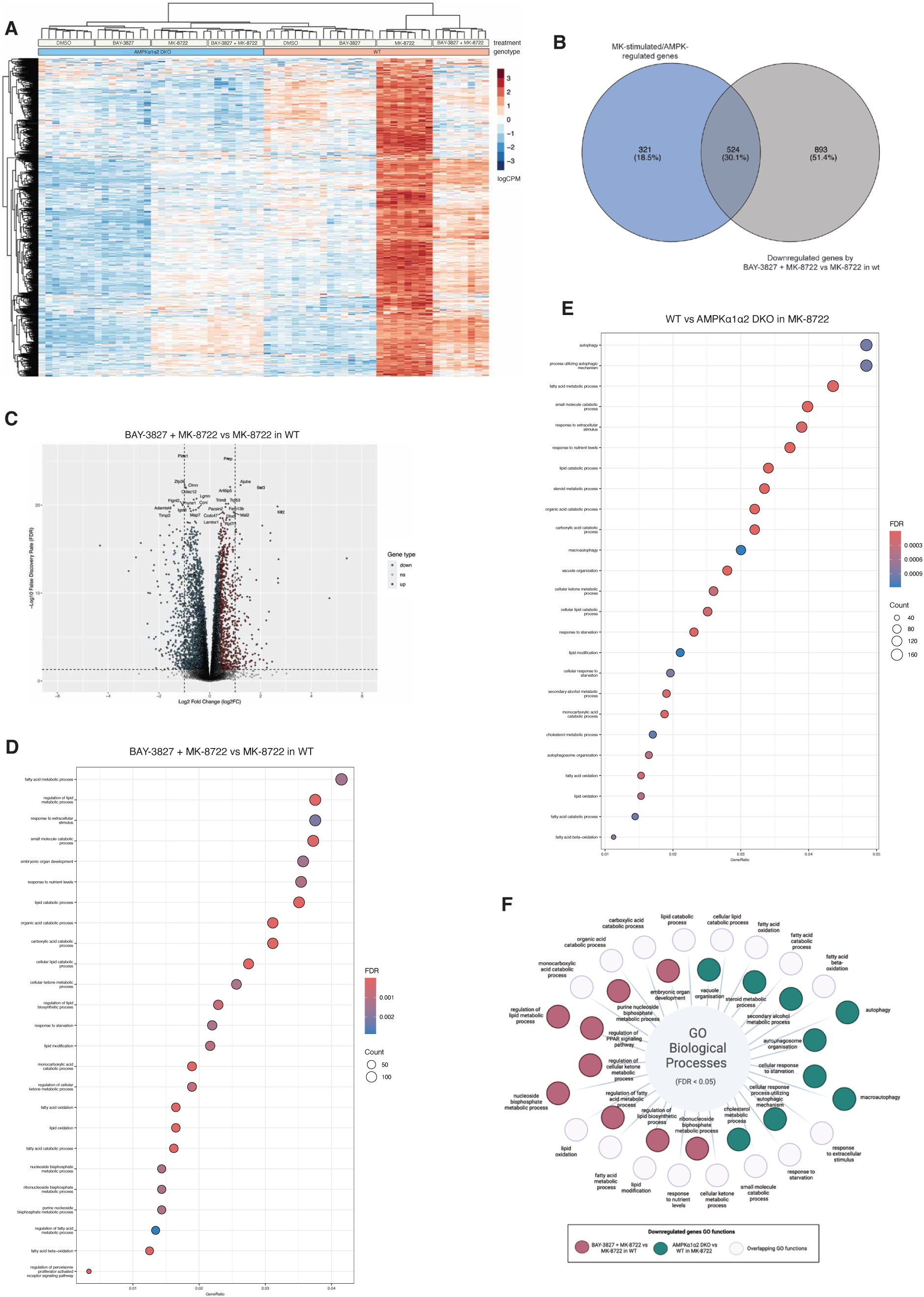
Unbiased transcriptome analysis of BAY-3827-treated wild-type and AMPKα1α2 DKO primary hepatocytes in combination with MK-8722. **A)** Heat map of gene expression of 845 significant genes which are stimulated by MK-8722 and are AMPK-dependent. **B)** Venn diagram showing the overlap between genes with a significantly reduced expression when treated by BAY- 3827 + MK-8722 in wild-type compared to MK-8722-stimulated/AMPK-regulated genes, with a 30.1% overlap. **C)** Volcano plot showing top significant (FDR < 0.05) upregulated (FC ≥ 1.3; red) and downregulated (FC < - 1.3; blue) genes by BAY-3827 + MK-8722 compared to MK-8722 treatment alone in wild-type cells. **D)** Gene ontology biological process enrichment analysis downregulated significant genes (FDR < 0.05) by BAY-3827 + MK-8722 treatment and **E)** by the loss of AMPKα1α2 in MK-8722 treatment. **F)** Schematic of the downregulated gene ontology (GO) biological processes showing the functional overlap between AMPKα1α2 DKO vs wild-type hepatocytes and BAY-3827 + MK-8722 vs MK-8722 in wild-type.

Specific functions in downregulated genes in wild-type cells treated with BAY-3827 and MK- 8722 compared to MK-8722 alone included nucleoside biphosphate, ketone metabolism, and regulation of peroxisome proliferator-activated receptor (PPAR) signaling pathway (**Fig. 5D and F**). Unique downregulated functions in MK-8722-treated AMPKα1α2 DKO compared to MK-8722-treated WT cells included autophagy-related functions alongside steroid and secondary alcohol metabolic processes (**Fig. 5E and F**). BAY-3827 treatment in the presence of MK-8722 resulted in a significant decrease of AMPK-dependent genes involving lipid metabolic genes which in MK-8722 alone conditions are upregulated to promote lipid oxidation and other catabolic processes (**Fig. S3C**). AMPK-dependent genes that were downregulated by BAY-3827 treatment included fatty acid metabolism genes *PLIN5*, *LPIN2*, *EHHADH*, *AVPR1A*, *ADORA2B* as well as liver metabolism regulator *SDS* and histidine catabolism gene *HAL*, establishing BAY-3827’s role in suppressing the expression of AMPK-regulated lipid oxidative genes (**Table 4, Fig. S3C**). Apart from lipid metabolic genes, BAY- 3827 also reduced the expression of AMPK-regulated transcription genes such as *SP1* which encodes a transcription factor proposed to be AMPK-regulated (*49*). Another example includes the *TBCD15* and *TBCD17* genes with roles in autophagy, with encoded Rab-GTPase activating protein TBCD17 recently described AMPK target (*50,51*) (**Table S3**).

**Table 4.**
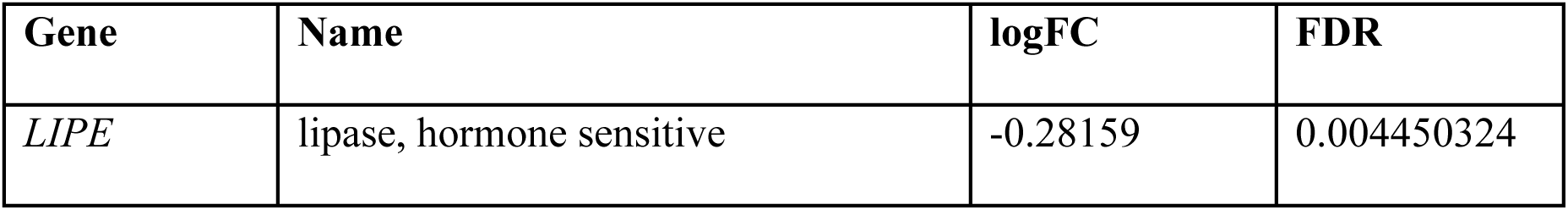
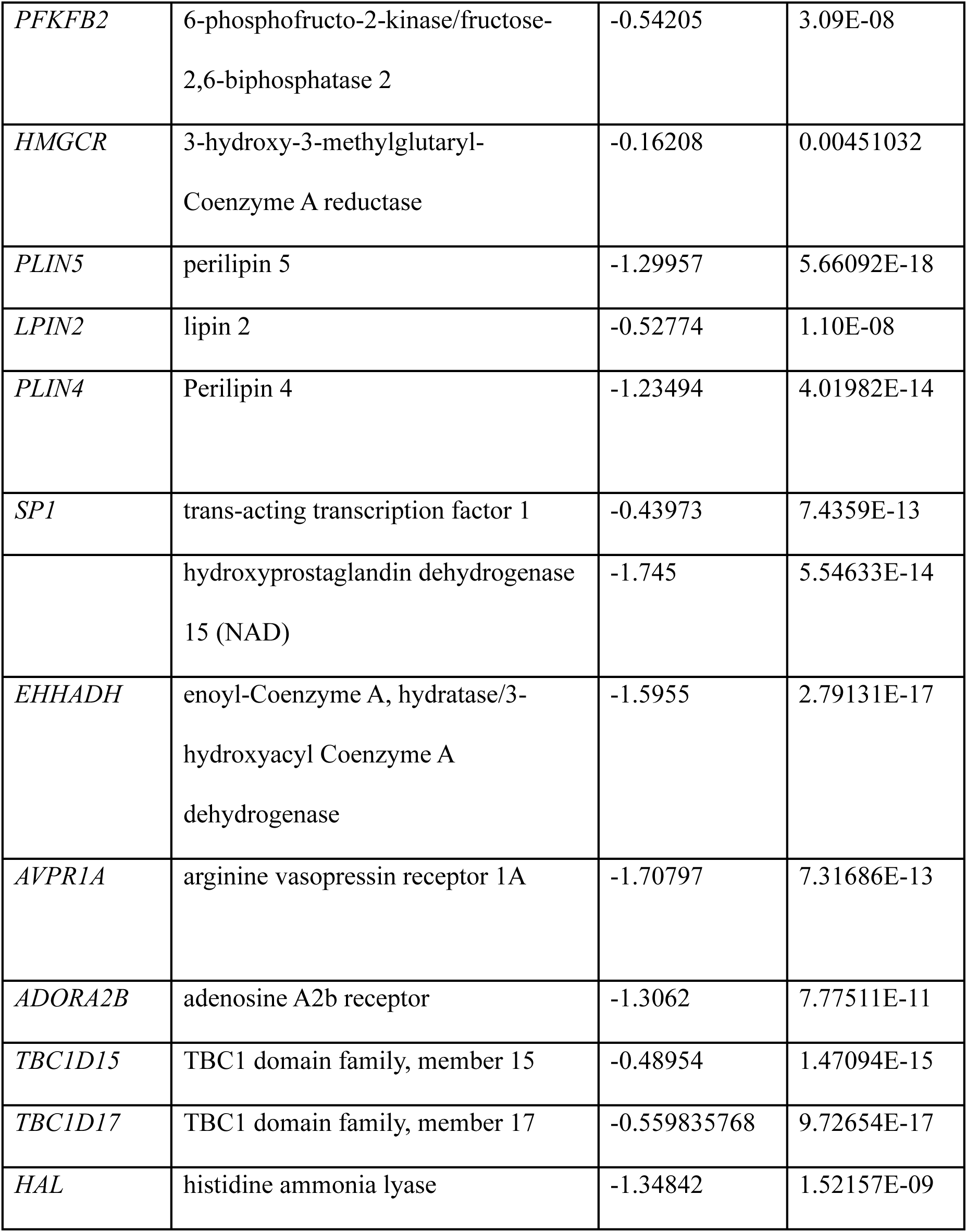
Examples of AMPK-dependent genes downregulated in BAY-3827 in combination with MK-8722-treated wild-type hepatocytes. Significant differentially expressed genes recorded in Table S3.

Yang *et al.* recently described RSK-isoform specific transcriptional profiles in cancer and RSK-specific genes using RSK1- and RSK2 knockout glioma-based cell lines (*52*). The most significant RSK-regulated genes found in our dataset in genes downregulated by BAY-3827 treatment in combination with MK-8722 compared to MK-8722 alone in WT cells were RSK2-proposed regulated genes *STAT5B* and *CALCOCO1* and RSK1-proposed regulated genes such as *NEK2, DSN1, MASTL* and *SPC25* (**Fig. S3D, Table S3**) (*52*).

### Cellular effect of BAY-3827 on RSK

Even though previous (*37*) and current in vitro (cell-free) data (**Fig. 1A**) have shown that BAY-3827 potently inhibits RSK isoforms, this has not been verified in a cellular context. We treated serum-starved HEK293 cells with varying concentrations of BAY-3827 or RSK inhibitor BI-D1870 (*53*) with or without phorbol 12-myristate 13 acetate (PMA), an activator of the ERK1/2-RSK isoforms, and assessed phosphorylation of GSK3α/β as a surrogate marker for cellular RSK activity (*53*). Since phosphorylation of GSK3α/β is also regulated by Akt, we included an Akt inhibitor MK2206 to specifically assess RSK-mediated (i.e., PMA-stimulated) phosphorylation of GSK3α/β. As anticipated, BAY-3827 treatment resulted in a dose-dependent reduction of ACC1 phosphorylation, while BI-D1870 treatment had no effect (**Fig. S4A and B**). MK-2206 treatment reduced basal pGSK3αβ. PMA robustly increased pERK1/2 and pGSK3αβ levels under MK-2206-treated condition (**Fig S4C and D**). PMA-stimulated pGSK3αβ (in the presence of MK-2206) was inhibited to the same extent by 10 μM BAY-3827 and BI-D1870 (**Fig. S4D**), indicating that BAY-3827 has a similar inhibitory potency to BI-D1870 in suppressing RSK in HEK293 cells. To explore the molecular basis for RSK inhibition by BAY-3827, a superimposition of the BAY-3827-AMPKα2-KD structure was conducted together with available RSK2 N-terminal (NT) kinase domain structures, with BI-D1870 and SL0101 inhibitors (*54,55*). Interestingly, both compounds showed overlapping binding sites with BAY-3827-AMPKα2-KD as well as having conserved interacting residues with AMPK (**Fig. S4E, F**). Additionally, both inhibitors have hydrogen bond contacts with hinge region Glu^94^ (Asp^148^ in RSK2) and Val^96^ (Leu^150^ in RSK2) where the latter in SL0101 is water-mediated (**Fig. S4E and F**), implicating that BAY-3827’s effect on RSK inhibition is likely derived from these conserved structural features and the compound binding mode.

## DISCUSSION

Here we provide the structure-based mechanism of action of BAY-3827 in inhibiting AMPK with distinct and new molecular features. We also have comprehensively evaluated the cellular potency and selectivity (i.e., on/off-target effects) of BAY-3827 alongside its inactive control BAY-974 in signaling, transcriptomic and metabolic output levels employing WT and AMPKα1α2 DKO primary hepatocytes. Consistent with a previous study (*37*), we show that BAY-3827 potently inhibits AGC group kinases RSK1 and RSK2 in cell-free assays, however to a lower extent than AMPK, as shown across a panel of 140 human protein kinases (**Fig 1A, Table S1**). Compound C and SBI-0206965, based on their reported selectivity profiles (*31, 33, 36*), did not affect the RSK family, however, other compounds reported to inhibit AMPK have also shown a preference for RSK family kinases. For instance, ATP-competitive PAK4 inhibitor PF-03758309 has been reported to potently inhibit AMPK with an *IC*_50_ of 1 nM in a cell-free assay. PF-03758309 is also known to potently inhibit RSK1 and RSK2 with an *IC*_50_ of 2 nM (*56*), and in silico docking to AMPK kinase domain predicted three hydrogen bonding interactions in the hinge region with Glu^94^ and Val^96^ residues (*57*) in the same fashion as in the current AMPKα2 KD-BAY-3827 structure. RSK inhibitors BI-D1870 and SL0101 both make contacts with these two hinge residues which correspond to Asp^148^ and Leu^150^ in the reported RSK2 structures (*54,55*) (**Fig. S4E and F**), displaying overlapping binding sites compared to BAY-3827, pointing to a structural basis for RSK inhibition. In line with this notion and with results from current and previous (*37*) cell-free assays, our cellular studies reveal that BAY-3827 treatment inhibited PMA-stimulated RSK activation to the same extent as BI-D1870 (**Fig. S4A and D**). Thus, RSK isoforms are validated off-targets of BAY-3827, and RSK inhibitors could be included as an additional control alongside BAY-974 (which does not inhibit RSKs) to mitigate the involvement of RSK isoforms.

The AMPKα2 KD-BAY-3827 co-crystal structure revealed that BAY-3827 binds in the ATP-binding pocket at a site overlapping with SBI-0206965, an observation consistent with compound C’s ATP-competitive nature as opposed to SBI-0206965 and BAY-3827 which have been reported as mixed-type inhibitors (*29,33,46*). Superimposition of AMPKα2 KD- BAY-3827 with SBI-0206965- and compound C-bound structures revealed an activation loop conformation characteristic of an inactive kinase, with a conserved αC-helix-out state (**Fig. S1F**). However, in BAY-3827-bound conformation, activation loop residue Asn^162^ is stabilized in a position which disrupts the AMPK regulatory spine and His^137^ is displaced (**Fig.S1G**). Phe^158^ residue faces down towards the C-lobe compared to the active state, with a conserved conformation compared to the SBI-0206965-bound structure (**Fig.2F, S1D**), which is likely due to their shared binding sites. The AMPKα2(T172D)-SBI-0206965 structure (*33*) was first described as a type IIB structure in a non-canonical DFG-out conformation, however distance and dihedral measurements following an updated kinase nomenclature (*42*) has revealed that it clusters together with the BAY-3827-bound structure with “DFG-in” states (**Table 2**). DFG-in structures are often associated with active kinases; however they are also found in inactive states (*42–44*). In these structures, an inactive state is achieved either due to αC helix movement or activation loop conformation (*44*). In the case of BAY-3827 binding, we find that both factors contribute to this, mainly by disrupting the regulatory spine which is otherwise uninterrupted in an active kinase. Given the conserved orientation of the Phe^158^ ring towards the C-lobe we propose that an “FG-down” conformation, known to be associated with “DFG-in” states (*43*), might be important in AMPK structural dynamics and inhibition.

We further observed a cysteine disulfide bridge unique to the AMPKα2-KD-BAY-3827 structure between AMPK-specific αD Cys^106^ and activation loop Cys^174^, which we hypothesize helps to stabilize activation loop conformation in an unproductive kinase state upon BAY-3827 binding (**Fig. 2D**). This interaction has not been observed previously in available AMPK KD-inhibitor co-structures, and Cys^174^ alongside the close-by Phe^102^ which is displaced as a result of this interaction were among the top residues predicted as highly conserved in BAY-3827 inhibited kinases but were lowly conserved in BAY-insensitive kinases (**Fig. 1F, Table S2**), suggesting that the A-segment stabilization is a contributing factor to BAY-3827 inhibition.

We also report that BAY-3827 increases AMPK-Thr^172^ phosphorylation in U2OS cells, mouse primary hepatocytes, and mouse primary adipocytes (**Fig. 4, S2**). It was recently suggested that the binding of BAY-3827 (and also SBI-0206965) (*33,46*) induces a protective effect towards phosphatase-meditated dephosphorylation of Thr^172^ (*46*). However, given the dose-dependent reduction in AMPK downstream target phosphorylation such as ACC1 and Raptor, we also find that it does not reflect functional activation status of AMPK. Rather BAY-3827 likely stabilizes an activation loop conformation which contributes to phosphatase access shielding as well as an inactive conformation for substrate engagement.

We have demonstrated that BAY-3827 is capable of fully reversing MK-8722-induced inhibition of fatty acid synthesis and downregulating >30% of MK-8722-stimulated AMPK-regulated genes (**Fig. 4B**) involved in known AMPK-governing functions such as fatty acid oxidation and lipid catabolism (**Fig. 4D-F**). Lemos et.al treated androgen-dependent prostate cancer cells (LNCaP) with BAY-3827 and reported a number of downregulated genes involved in lipid metabolism that we also observed were downregulated under BAY-3827 treatment in primary hepatocytes in the presence of AMPK activator MK-8722, namely *LIPE, PFKFB2* and *HMGCR* (*37*). Our analysis also uncovered other lipid metabolism related genes with expression dependent on the presence of functional AMPK, which were significantly downregulated following the treatment with BAY-3827 when AMPK had been pharmacologically activated (**Fig.S3A, C**). The RSK family encompasses a conserved group of serine/threonine protein kinases formed by human isoforms RSK1, RSK2, RSK3 and RSK4 alongside RSK-like mitogen- and stress-activated kinases 1 (MSK1) and 2 (MSK2), with key functions in cellular growth and proliferation such as cell survival promotion, protein translation and ribosome biogenesis (*58*). Because of their growth-promoting roles alongside ensuring cell survival, RSKs are also established targets in anticancer therapies and are associated with increased expression in certain types of cancer (*59*). We searched for RSK-regulated genes (*52*) in our dataset and found that BAY-3827 in combination with MK-8722 significantly downregulated some genes, with a larger number of previously defined RSK1 genes present in our dataset compared to RSK2 (**Fig.S3D**). This is consistent with our data showing higher inhibition of RSK1 compared to RSK2 isoform (**Table S1**) and warrants further investigations.

As new emerging roles and therapeutic implications of AMPK arise, it is critical to perform robust proof-of-concept experiments with multifaceted approaches. Even though there are well established preclinical genetic models (*60, 61*) and allosteric activators (*8*), availability of potent and selective inhibitors has been limited. We here delineated the molecular basis of the BAY-3827’s inhibition of AMPK and AMPK-regulated transcriptomic and metabolic functions, alongside building on the knowledge of AMPK-specific structural features. We have also described the current limitations of BAY-3827 (e.g., potent inhibition of RSK) and propose the use of inactive control BAY-974 and context dependent use of RSK inhibitors.

BAY-3827 offers greater potency and selectivity compared to prevalently used inhibitors (e.g., compound C, SBI-0206965), making it the preferred choice for cell-based studies. However, potent inhibition of RSK isoforms and poor pharmacokinetic properties limit its in vivo use, and these issues will need to be addressed in future efforts to identify AMPK inhibitors.

## MATERIALS AND METHODS

### Animals

Animal experiments were conducted in accordance with the European directive 2010/63/EU of the European Parliament and of the Council of the protection of animals used for scientific purposes. Ethical approval was given by the Danish Animal Experiments Inspectorate (license number #2021-15-0201-01059). For primary hepatocytes and ex vivo skeletal muscle incubation experiments, wild-type (WT) C57BL/ 6N Tac male mice (10-16 weeks old) were obtained from Taconic Biosciences and housed in the animal facility at the Faculty of Health and Medical Sciences (University of Copenhagen). Animals were anesthetized with Avertin [stock of 1 g/ml tribromoethanol (#T48402, Merck MilliporeSigma) in 2-methyl-2-butanol (#152463, Merck MilliporeSigma), diluted 1:20 in saline, and dosed at 10 µl/g body weight via intraperitoneal injection prior to the procedure.

For adipocyte experiments, animal experiments were approved by the Regional Ethical Committee on Animal Experiments in Malmö/Lund (approval number 5.8.18–19111/2023). WT C57Bl6/J male mice (10–11-week-old) were obtained from Taconic Biosciences and kept in the animal facility at the Biomedical Centre (Lund University, Sweden).

All the animals were kept and maintained according to local regulations under a light/dark cycle of 12 h, 22°C ± 1°C, and had free access to water and standard chow diet.

### Mouse primary hepatocyte isolation and lipogenesis assay

Mouse hepatocytes were isolated by collagenase perfusion as previously described (*48*) Hepatocytes were seeded in medium Eagle-199 (MEM-199) (#41150, Thermo Fisher Scientific) containing 100 U/ml penicillin G, 100 µg/ml streptomycin and 10% (vol/vol) FBS. Hepatocytes were left for attachment (3 - 4 h) and serum-starved overnight at 37°C with 5% CO_2_ in MEM-199 supplemented with 100 U/ml penicillin G, 100 µg/ml streptomycin 10 nM insulin, and 100 nM dexamethasone. Primary hepatocytes isolated from liver-specific AMPKα1α2 double knockout (DKO), or wild-type (WT) animals were seeded and after 16- 18h incubated for 3 h for signalling studies as previously described (*47*) or were incubated with MEM-199 media supplemented with 0.6 μCi/ml [^14^C]-acetate in the presence of BAY- 3827 or inactive BAY-974 co-treated with MK-8722 or vehicle (0.1% DMSO) respectively. Cells were then harvested in 0.5 ml PBS, transferred into 1 ml 40% KOH and 2 ml methanol followed by 1 hour incubation at 80°C. Lipids were saponified by acidifying the samples in 37% HCl and extracted with petroleum ether. Extracts were allowed to evaporate to dryness, then dissolved in Ultima Gold scintillation fluid for determination of [^14^C]-acetate incorporation into lipids.

### Mouse primary adipocyte isolation, compound treatment and glucose uptake

Epidydimal adipose tissue excised from mice was digested with 1 mg/ml collagenase in Krebs-Ringer medium (120 mM NaCl, 4.7 mM KCl, 1.2 mM KH_2_PO_4_, 1.2 mM MgSO_4_) containing 25 mM HEPES (pH 7.4), 200 nM adenosine, 2 mM glucose and 3% (wt./vol.) BSA (KRH buffer) at 37°C in a shaking incubator. Adipocytes were isolated by filtering and washing in KRH buffer. For preparation of protein extracts for western blot, adipocyte suspensions (0.7-1ml of 10% (v/v) cells) were pre-incubated with BAY-3827/BAY-974 compounds or 0.1% DMSO for 1 hour at 37°C in a shaking (80 rpm) water bath followed by either 1 hour incubation with 10 µM MK-8722 or 30 min with 0.1 nM insulin in KRH buffer, as indicated in the figure legends. Cells were then washed in KRH buffer without BSA and lysed in 50 mM Tris HCl (pH 7.5), 0.27 M sucrose, 1 mM EDTA, 1 mM EGTA, 5 mM sodium pyrophosphate, 1 mM sodium orthovanadate, 50 mM sodium fluoride, 1 mM dithiothreitol, 1 % (wt./vol.) NP-40 and complete protease inhibitor cocktail (one tablet/50 ml) included in lysis buffer. Lysates were centrifuged at 13,000 g for 15 min (4°C) and supernatant protein concentration was determined by the Bradford assay.

Freshly isolated mouse adipocytes were washed in glucose-free buffer containing 30 mM HEPES pH 7.4, 120 mM NaCl, 4 mM KH2OPO4, 1 mM MgSO4, 0.75 mM CaCl2, 10 mM NaHCO3, 200 nM adenosine and 3% (w/v) BSA (KRBH buffer). Adipocyte suspensions (400 µl of 3.75% (v/v) cells) were pre-incubated with BAY compounds or 0.1% DMSO for 1 hour at 37 °C in a shaking (80 rpm) water bath, before being stimulated with 0.1 insulin or 10 µM cytochalasin B for 30 min. Subsequently, 100 µl KRBH buffer containing 0.25 µl (0.025 µCi) [14C]-glucose (275 mCi/mmol glucose; final glucose concentration = 0.18 µM) was added and cells were incubated for a further 30 min. Reactions were stopped by aliquoting 300 μl of the total 500 μl adipocyte suspension to Beckman microtubes containing 75 μl of dinonylphtalate. The adipocyte suspension was centrifuged at 6000g and frozen at −20°C before adipocytes were collected and subjected to scintillation counting. The assay was performed in triplicate for each condition.

### Ex vivo skeletal muscle incubation for signaling and glucose uptake assays

Extensor digitorum longus (EDL) muscles were rapidly dissected and mounted in oxygenated (95% O_2_ and 5% CO_2_), and warmed (30°C) Krebs Ringer buffer (KRB) supplemented with 2 mM pyruvate in the presence of the BAY-3827 or vehicle (0.1% DMSO) for 30 min. The muscles were then co-incubated for 50 min with BAY-3827 and ± MK-8722 (10 µM) or insulin (100 nM).The muscles were further incubated for 10 min in the presence of vehicle, drugs or insulin in glucose uptake buffer (KRB buffer containing 1.5 mCi/ml [^3^H]-2-deoxy-D-glucose, 1 mM 2-deoxy-D-glucose, 0.45 mCi/ml [^14^C]-mannitol, 7 mM mannitol). At the end of incubation period, muscles were frozen in liquid nitrogen and subsequently processed for glucose uptake (as described in *62*) and western blot analysis.

### Kinase selectivity assays

BAY-3827 and BAY-974 were assayed against a 140-human kinase panel at 0.1 μM concentration in a [γ-^33^P] ATP-based radioactive filter binding assay (*63, 31*) setup in duplicates with reported mean remaining kinase percentage activity alongside standard deviation (**Table S1**). Assays were performed at either 5, 20, or 50 μM in order to be at or below the Km for ATP for each enzyme and conducted by the services offered by the International Centre for Kinase Profiling at MRC Protein Phosphorylation and Ubiquitylation Unit, University of Dundee (https://www.kinase-screen.mrc.ac.uk/services/premier-screen).

### Protein production and kinase assays

Heterotrimeric human AMPK FLAG-α1β1γ1, Flag-α2β1γ1 and FLAG-α2β2γ1, as well as human FLAG-CaMKK2 and FLAG-ULK1, were produced in mammalian HEK293 cells as described (*64,65*). For AMPK expression, cells were triply transfected at 60% confluency using FuGene HD (Roche Applied Science) and 1 μg of pcDNA3 plasmid expression constructs for AMPK Flag-α1, β1-Myc, β2-Myc and HA-γ1. For CaMKK2 expression, the cells were transfected with 1 μg of pcDNA3 Flag-CaMKK2 plasmid. For ULK1 expression, the cells were transfected with 2 μg of pcDNA3 FLAG-ULK1 plasmid. After 48 h, the transfected cells were harvested by rinsing with ice-cold PBS, followed by rapid lysis using 500 μl of lysis buffer. AMPK, CaMKK2 and ULK1 kinase activities were determined by phosphorylation of synthetic peptide substrates: SAMS (HMRSAMSGLHLVKRR), CaMKKtide (LSNLYHQGKFLQTFCGAPLYRRR) and S108tide.

(KLPLTRSHNNFVARRR), respectively. Briefly, recombinant AMPK, CaMKK2 or ULK1 were immunoprecipitated from 10 μg of transfected cultured cell lysate using 10 μl of anti-FLAG M2 agarose beads (50% (v/v)) (Merck MilliporeSigma), washed and then added to a 25 μl reaction containing assay buffer (50 mM HEPES-NaOH, pH 7.4, 1 mM DTT, and 0.02% (v/v) Brij-35), 200 μM synthetic peptide substrate (SAMS, CaMKKtide or S108tide), 200 μM or 20 μM [γ-^32^P]-ATP (PerkinElmer), 5 mM MgCl_2_, in the presence of BAY-3827 (0–1 μM) or BAY-974 (0–1 μM). Reactions were performed at 30°C and terminated after 10 min by spotting 15 μl onto phosphocellulose paper. Radioactivity was quantified by liquid scintillation counting. Data visualisation and non-linear regression fitting was performed in GraphPad Prism v10.

### HEK293 cell culture

HEK293 cells were seeded at a density of 500,000 cells/well in 6 well cell culture plates (Corning Incorporated Costar #3516) in Dulbecco’s Modified Eagle Medium (DMEM) high glucose media + GlutaMAX^TM^ Supplement, pyruvate (Thermofisher CAT# 31966-021) containing 100 U/ml penicillin G, 100 µg/ml streptomycin and 10% (vol/vol) FBS. Once attached, the next day the media was changed to serum-free and the cells were starved for 16 hours followed by compound treatment with 2X BAY-3827 (MedChemExpress, #HY- 112083), BI-D1870 (MedChemExpress, #HY-10510) ± 10 µM MK-2206 dihydrochloride (MedChemExpress, #HY-10358) with 30 min incubation at 37°C followed by co-incubation with 0.1 PMA µM (MedChemExpress, #HY-18739) or vehicle for additional 30 min.

### Western blotting

Following compound treatment, cells were washed with room temperature PBS and lysed on ice with the addition of lysis buffer (50 mM Tris-HCl, pH 7.5, 150 mM NaCl, 1 mM EDTA, 1 mM EGTA, 0.27 M sucrose, 1% w/v Triton X-100, 20 mM glycerol-2 phosphate disodium, 50 mM NaF, 5 mM Na_4_P_2_O_7_.10 H_2_O) containing proteases and phosphatase inhibitors (0.5 mM PMSF, 1 mM benzamidine HCl, 1 µg/ml leupeptin, 1 µg/ml pepstatin A, 1 µM microcystin-LR, 1 mM Na_3_VO_4_ and 1 mM DTT). Lysates were centrifuged at 10000 g, 4°C for 10 min and protein concentration of the supernatant was determined with the Pierce™ Bradford Protein Assay Kit (Thermo Scientific™, #23200) and read on the Hidex™ Plate Reader (Software Version 0.5.64.0). Samples were prepared in Laemmli Sample Buffer (Tris 200 mM, 8% SDS, EDTA 2 mM, 40% Glycerol) supplemented with 10% DTT and boiled for 5 min at 100°C. 10 µg of protein per sample was run on a Criterion™ Tris-glycine polyacrylamide gel (Bio-Rad, #5678095), transferred to a nitrocellulose membrane. For cell signalling experiments in Hek293 cells, samples were run on a Criterion TGX Stain-Free Precast gel (Bio-Rad, CAT #5678085) with transfer to a PVDF membrane performed in BioRad Criterion blotter (#1704071) using Tris-glycine transfer buffer (Bio-Rad, #1610734) following manufacturer’s instructions. Membranes were probed at 4°C overnight in a 4% BSA TBS-T solution containing 0.02% sodium azide with the following antibodies where appropriate. Phospho-Acetyl-CoA Carboxylase (Ser79) (1:1000, Cell Signaling Technology, #3661S), Acetyl-CoA Carboxylase (C83B10) (1:1000, Cell Signaling Technology, #3676S), Phospho-Raptor (Ser792) (1:1000, Cell Signaling Technology, #2083S), Raptor (24C12) (1:1000, Cell Signaling Technology, #2280S), GSK3αβ (1:2000, Sigma, #04-903), Phospho- GSK3αβ (Ser21/9) (D17D2) (1:1000, Cell Signaling Technology, #8566S), ERK1/2 (137F5) (1:1000, Cell Signaling Technology, #4695), Phospho-ERK1/2 (Thr202/Tyr204) (D13.14.4E) (1:1000, Cell Signaling Technology, #4370), Phospho-AMPKα (Thr172) (40H9) (1:1000, Cell Signaling Technology, #2535S), AMPKα (1:1000, Cell Signaling Technology, #2532S), Anti-GAPDH antibody (1:5000, Cell Signaling Technology, #G8795) and vinculin (1:1000, Cell Signaling Technology, #13901T). Membranes were scanned using the Odyssey® Fc LiCor system (exposure time 30s) with densitometry analysis performed using LiCor Image Studio™ Lite normalising to control loading protein. Figures were made using GraphPad Prism v10.

For adipocyte experiments, cell lysates (25 μg protein) were heated at 95°C for 3 min in LDS sample buffer, subjected to polyacrylamide gel electrophoresis on precast Novex gradient gels (Thermo Fisher Scientific), and electrotransferred to nitrocellulose membranes.

Membranes were blocked for 30 min in 50 mM Tris-HCl pH 7.6, 137 mM NaCl, and 0.1% (wt/vol) Tween 20 (TBS-T) containing 10% (wt/vol) skimmed milk and then probed with the indicated antibodies in TBS-T containing 5% (wt/vol) BSA for 16 h at 4°C. The following antibodies were used: anti-AMPK (#2603, 1:1000), anti-AMPK-pT172 (#2535, 1:1000), anti-Raptor (#2280, 1:1000), anti-Raptor-pS792 (#2083, 1:1000), anti-ACC (#3662, 1:1000), anti-ACC-pS79 (#3661, 1:1000), anti-PKB/Akt (#9272, 1:1000), anti-PKB/Akt-pT308 (#9275,1:1000) and anti-GSK3α/β-pS21/9 (#9331, 1:1000) were all purchased from Cell Signaling Technology (Danvers, United States). Anti-AS160 (#07-741, 1:1000) was from MerckMilliporeSigma, anti-AS160-pT649 (#44-1071G, 1:1000) from Thermo Fisher Scientific and anti-HSP90 from BD Biosciences (San Jose, CA, United States). Detection of primary antibodies was performed using horseradish peroxidase (HRP)-conjugated secondary antibodies, and SuperSignal® West Pico or Femto Chemiluminescent Substrates (Thermo Fisher Scientific). Luminescence signals were visualized in a ChemiDoc XRS+ (Bio-Rad) and quantified by densitometry using the software Image LabTM 5.1 (Bio-Rad). Signals were normalized to the loading control HSP90.

### Homogeneous Time-Resolved Fluorescence (HTRF) Assays

Human osteosarcoma U2OS wild-type cells were cultured in Dulbecco’s Modified Eagle Medium (DMEM) high glucose media + GlutaMAX^TM^ Supplement, pyruvate (Thermofisher CAT# 31966-021) containing 100 U/ml penicillin G, 100 µg/ml streptomycin and 10% (vol/vol) FBS seeding with 50,000 cells per well in a 50 µl volume in culture 96-well cell culture plates (Corning Incorporated costar, #3599). The next day, the media was changed to 50 µl of serum-free media and 50 µl of serum-free media containing 2X BAY-3827 or BAY- 974 was dispensed and left to incubate at 37°C for 30 min. Media was aspirated and replaced with 50 µl of serum-free media containing either 2X MK-8722 (10 µM) or vehicle and incubated at 37°C for 1 h, with a final DMSO assay concentration of 1.03%. HTRF assays using either phospho-Ser79-ACC kit (Revvity, #64ACCPEG) or phospho-Thr172-AMPK kit (Revvity, #64AMPKPEG) were performed in triplicate in HTRF 96-well low volume white plates (Revvity, #66PL96001) following manufacturer’s instructions. Sealed plates incubated overnight in darkness conditions were scanned at room temperature in Multimode Plate Reader EnVision (PerkinElmer) machine following HTRF setup recommendations for EnVision (Revvity), equipped with a top mirror LANCE/DELFIA single mirror (#412), a UV2 (TRF) 320 excitation filter (#111), emission filter APC 665 (#205) and second emission filter Europium 615 (#203), with a 2000 μs cycle and 60 μs delay, with the number of flashes set to 100 as well as for the 2^nd^ detector. Results were viewed in EnVision Manager software (Version 1.13.3009.1401) where 665/615 ratios were calculated, and data was normalised to control condition as %. All assay ratios between control lysate signal/ non-specific signal were greater than 2 as expected of a successful assay following manufacturer instructions.

Data was visualised in GraphPad Prism v10.

### Primary hepatocyte RNA isolation, sequencing and bioinformatics analysis

Primary hepatocytes were pre-treated for 15 min with BAY-3827 5 μM at 37°C in MEM-199 supplemented with 100 U/ml penicillin G, 100 μg/ml streptomycin and 100 nM dexamethasone followed by addition of MK-8722 10 μM treatment. 6h-post-treatment, cells were washed twice with room-temperature PBS. RNA was isolated using QIAwave RNAeasy kit (CAT #74104) with on-column DNAse treatment using Qiagen RNase-Free DNase set (CAT #79256) following manufacturer’s protocol. Messenger RNA sequencing was performed by the Single-Cell Omics platform at the Novo Nordisk Foundation Center for Basic Metabolic Research. Libraries were prepared using the Universal Plus mRNA-seq protocol (Tecan) as recommended by the manufacturer. Libraries were quantified with NuQuant using the CLARIOstar Plate Reader (BMG Labtech), quality checked using a TapeStation instrument (Agilent Technologies) and subjected to 52-bp paired-end sequencing on a NovaSeq 6000 (Illumina). The sequencing analysis pipeline nf-core (*66*) was used, with alignment and quantification selection of STAR and Salmon, and mouse GRCm38 set as the reference genome. UMI-based deduplication was performed using UMI-tools and sorting and index alignments were completed with SAMtools as part of the pipeline. Differential expression analysis was carried out using the R package edgeR (*67*) and a batch-effect removal was performed in limma (*68*) to account for the variability induced by two mice (291 + 295) from which cells with the same treatments had been pooled to obtain the required cell confluence. Venn diagrams were produced using the R package ggvenn (*69*). Volcano plots were produced using the R package ggplot2 and significant upregulated or downregulated genes were described as having a fold change (FC) ≥1.3 and false discovery rate (FDR) < 0.05. Heatmaps of log2 counts per million (logCPM) normalised counts were produced using R package pheatmap (*70*). Gene ontology (GO) enrichment analysis was performed using the R package clusterProfiler version 4.6.2 (*71*) setting FDR < 0.05 with reference organism specified as mouse, pAdjustMethod was set to BH and ontology category set to biological process (BP), with a qvalueCutoff of 0.05. Visualisation analyses were conducted in RStudio version 4.2.0 (*72*).

### AMPKα2-KD protein expression and purification

The gene sequence of human AMPKα2 kinase domain (α2KD; 6-280 aa) with a cleavable 6x- His and Avi-tag was custom synthesized, and codon optimized for bacterial expression and cloned in pET-14b plasmid. For protein expression the pET14b-⍺2KD construct was transformed in Bl21-DE3-RIL cells and induced with 0.2 µM IPTG at an optical density of 0.7 and incubated overnight in a shaker maintained at 18℃. For protein expression, the cell pellet was resuspended in buffer containing Tris 25 mM pH 8.0, NaCl 500 mM, Glycerol 10%, Imidazole 20 mM, TCEP 2 mM, MgCl2-5mM, PMSF 0.2 mM, benzamidine 1 mM, Lysozyme 0.3 mg/ml and sonicated for 10 min with 20s:40s on/off cycle with sample kept on ice all the time. The lysed cells were centrifuged at 30, 000 g to remove insoluble fraction and the clear lysate was filtered through 4 µM filter before loading onto the His-Trap column.

The protein bound column was washed with 10 CV of wash buffer containing Tris 25 mM, NaCl 500 mM, Glycerol 10%, Imidazole 20 mM, TCEP 2 mM, MgCl_2_-5mM pH 8.0. The protein elution was done using AKTA with a linear gradient of imidazole from 20 mM to 300 mM over 35 CV. The eluted protein fractions were pooled together and mixed with 1:50 molar ratio of TEV protease and dialyzed overnight in buffer containing Tris 25 mM, NaCl 200 mM, Glycerol 10%, 2 mM β-mercaptoethanol, MgCl2-5mM, pH 8.0. This step of dialysis and TEV-cleavage was skipped for α2KD protein preparation that was used for Dianthus assays. TEV-cleaved protein was subjected to gel filtration chromatography in buffer containing Tris 25 mM pH 8.0, NaCl 200 mM, Glycerol 10% TCEP 1 mM, using Superdex-S200-16/600 column. The pure fractions of the protein were concentrated to 5 mg/ml and flash frozen for storage for further use.

### Crystallization and structure determination

The crystallization screening was set up with various commercially available crystal screens with a protein concentration of 5 mg/ml in a sitting-drop vapour diffusion setup. 400 nl drops were setup with 1:1 ratio of protein to reservoir solution in a 96 well 2-drop MRC plates. For α2KD_BAY-3827 co-crystallization, the protein at 0.5 mg/ml was mixed with the BAY-3827 compound at a final concentration of 0.1 mM, which was then concentrated to 5 mg/ml. The apo-crystals of AMPKα2 kinase domain (α2KD) were grown in a condition containing 12.5% w/v PEG 1000, 12.5% w/v PEG 3350, 12.5% v/v MPD, 0.03 M each of magnesium chloride and calcium chloride, 0.1 M bicine/Trizma base pH 8.5. The α2KD_BAY-3827 co-crystals were grown in a condition containing 0.1 M ammonium sulphate, 0.3 M sodium formate, 0.1 M sodium cacodylate pH 6.5, 3% (w/v) γ-PGA, 3% (w/v) PEG 20000. The crystals were harvested and flash frozen in liquid nitrogen. Diffraction data were collected at 100K at I24 beamline, Diamond light source (DLS). The data was processed by autoPROC (*73*) in space group P21212. The structure of α2KD_BAY-3827 was solved by molecular replacement using the program PHASER (*74*) with a search model 2H6D (Apo-α2KD structure). The structure was built in coot (*75*) and further refined in phenix-refine (*76*). The figures were generated in PyMOL (*77*).

### Dianthus binding assays

The binding affinity of BAY-3827 to AMPK-⍺2 kinase (⍺2_KD) was measured by Dianthus (Nano Temper Technologies). BAY-3827 was diluted from a 100 mM stock in DMSO to 0.5 mM working concentration in the assay buffer with a final DMSO concentration of 2.5%, which was kept constant throughout the concentration gradient of the compound. His-⍺2_KD was labelled with His-tag labelling dye (His-Tag Labelling Kit RED-tris-NTA 2nd Generation). The ratio of fluorescence intensities (670/650 nm) over BAY-3827 concentration gradient ranging from 500 µM down to 0.015 µM. The concentration gradient was prepared by 1:1 serial dilution over 16 steps, with 3 technical repeats for each concentration. The ratio of fluorescence intensities was plotted against concentration gradient in Prism GraphPad using non-linear regression (one site-total binding). The graph of one of the representative experiments is shown above with a *K_d_* value of 1.227 µM (*K_d_* value of 0.8642 to 1.742 over 95% confidence interval). The binding experiment was repeated 3 times and the mean *K_d_* values of the inhibitor was calculated as 1.01 ± 0.18 µM.

### Statistical Analyses

Data points are plotted as mean ± standard error of the mean (SEM). Data was inputted and all statistical tests were performed using GraphPad Prism, version 10.1.0. One- or two-way analysis of variance (ANOVA) statistical test were conducted using Tukey’s correction for multiple comparisons, specific test will be stated in the figure legend. Significance was accepted as p <0.05. N number and statistical details for each experiment reported in each figure legend. For lipogenesis generated data, a one-way Welch and Brown-Forsythe ANOVA test was conducted to assess statistical significance which doesn’t assume that all the groups were sampled from populations with equal variances (skew in data), with Sunnett T3 multiple comparisons correction. To estimate half-maximal inhibitory concentration (*IC_50_*) values non-linear regression fitting was conducted using the model [Inhibitor] vs. response – Variable slope in GraphPad Prism (Version 10.1.0) and estimating values reported in nM.

## Supplementary Materials

Fig. S1. Sequence conservation and structural basis for BAY-3827 inhibition.

Fig. S2. BAY-3827 but not BAY-974 inhibit AMPK signaling in adipocytes and ex vivo skeletal muscle.

Fig. S3. Unbiased transcriptome sequencing of AMPKα1α2 null and wild-type primary hepatocytes treated with BAY-3827 ± MK-8722.

Fig. S4. Ribosomal S6 kinase (RSK) signaling and BAY-3827.

Table S1. BAY-3827 and BAY-974 kinase selectivity data at 0.1 μM across a panel of 140 human kinases.

Table S2. Kinase residue conservation scores calculated based sequence similarity based on in vitro kinase activity data of highly- and lowly-inhibited kinases by BAY-3827.

Table S3. Significantly differentially expressed genes by BAY-3827 in the presence of MK- 8722 in wild-type primary hepatocytes.

## Acknowledgements

We thank the Scientific Genome Consortium (SGC)-Frankfurt for their generous donation of chemical probe BAY-974. We acknowledge Lars R. Ingerslev, Christian Grønbæk, Mie Mechta and the staff at The Single-Cell Omics platform at the Novo Nordisk Foundation Center for Basic Metabolic Research (CBMR) for technical and computational expertise and support. We acknowledge Thomas Michael Frimurer and Michael Lückmann at the Computational Chemistry Unit at CBMR for their computational expertise and training support in virtual docking simulations. We acknowledge assistance and support from computational resources and infrastructure via the Health Data Science Sandbox (https://hds-sandbox.github.io), funded by the Novo Nordisk Foundation grant NNF20OC0063268.

## Author contributions

Conceptualization: KS, EZ, CFB, MSA Experimental design: KS, EZ, CFB, MSA, JWS, OG Investigation: CFB, MSA, JWS, ML, JC, ABA, HL

Data analysis: CFB, MSA, JC, ML, ABA, OG, JWS, EZ, KS

Research model development and establishment: M.F. Visualization: CFB, MSA

Project administration: CFB, KS Supervision: KS, EZ, OG Writing – original draft: CFB, KS

Writing – review & editing: All authors

## Data and materials availability

RNAseq data pertaining to this project has been deposited in the Gene Omnibus (GEO) repository under accession GSE290022. X-ray crystallography data has been deposited in the Worldwide Protein Data Bank (PDB) under accession 9IC2.

## SUPPLEMENTARY MATERIALS

**Figure S1.**
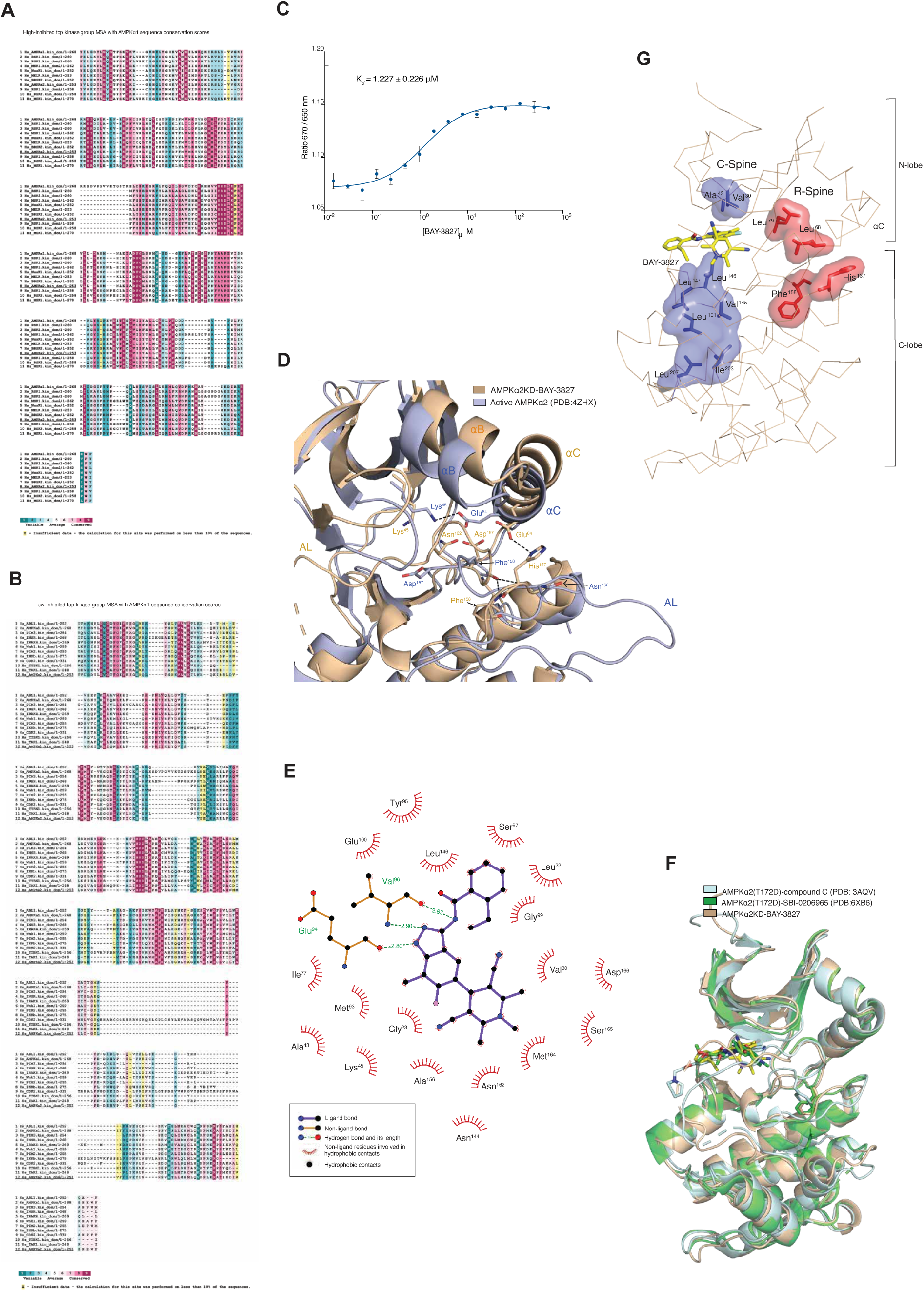
Sequence conservation and structural basis for BAY-3827 inhibition. **A)** Multiple sequence alignment of highly inhibited kinase group (≤50% remaining activity) by BAY-3827 showing computed ConSurf (*39,40*) residue conservation (1–9) reported in AMPKα2 numbering and in the low inhibited kinase group (≥99% remaining activity) shown in **B)** with yellow colors representing insufficient data to calculate a conservation score. **C)** Binding analysis of BAY-3827 to AMPKα2 kinase domain. Data is n=3 reported in mean ± SEM with *K_d_* value reported in μM. **D)** Superimposition of AMPKα2-KD-BAY-3827 structure (tan) with active AMPKα2-KD (purple, PDB: 4ZHX) with labelled displayed residues and features. **E)** Ligand plot of BAY-3827-interacting residues with AMPKα2 kinase domain showing hydrogen bonds and their lengths and ligand-protein hydrophobic contacts. **E)** Superimposition of compound C (light cyan, PDB:3AQV) and SBI-0206965 (green, PDB:6BX6)- AMPKα2(T172D) structures with AMPKα2 kinase domain (KD) bound to BAY- 3827. **G)** Catalytic (C) and regulatory (R) hydrophobic spines in AMPKα2-KD-BAY-3827 structure shown in purple and red respectively, alongside the composing residues. Visualizations were conducted in PyMOL (*77*).

**Figure S2.**
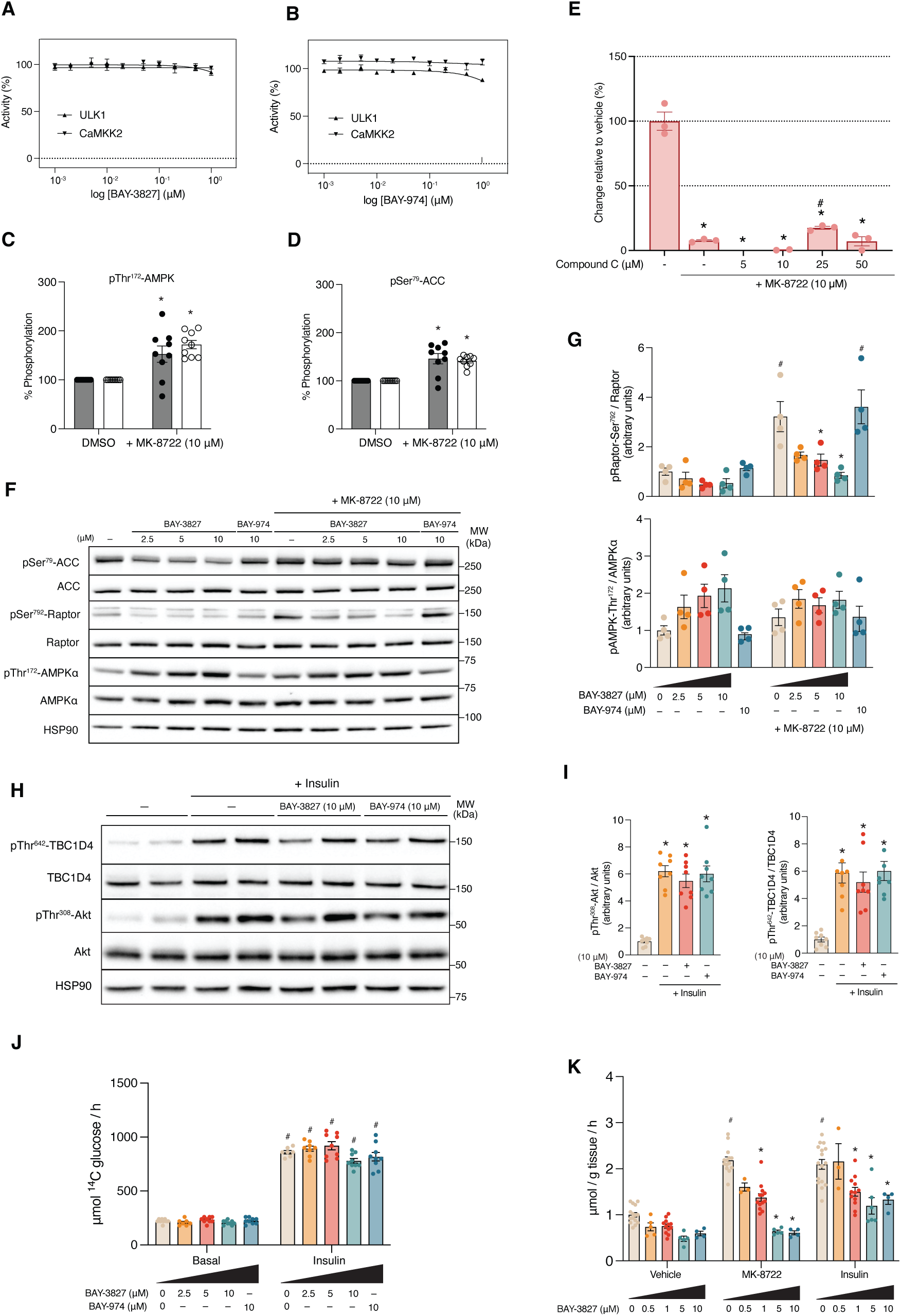
BAY-3827 but not BAY-974 inhibit AMPK signaling in adipocytes and ex vivo skeletal muscle. A) and. **B)** ULK1 and CaMKK2 kinase activity (%) assays in response to BAY-3827 or BAY-974 treatment from n=3. **C)** HTRF assay controls vehicle (DMSO) and MK-8722 10 μM in (C) pThr^172^-AMPK and **D)** pSer^79^-ACC kits. Data are n=3 from three independent experiments. **E)** Lipogenesis assay from n=6 mice seeded in n=3 technical triplicate wells. *p < 0.05 vehicle vs + MK- 8722 conditions; ^#^p < 0.05 vehicle + MK-8722 vs compound C + MK-8722 conditions. **F)** -**G)** Representative western blots of AMPK signalling in adipocytes treated with BAY-3827/BAY-974 in basal and MK-8722 (10 μM) conditions and **H)** Phospho/total ratios of pSer^792^-Raptor and pThr^172^- AMPK calculated from band intensities normalised to loading control. **H)** Representative western blot of insulin signalling in primary adipocytes treated with BAY-3827 or BAY-974 ± insulin and the corresponding calculated **I)** pThr^308^-Akt and pThr^642^-TBC1D4 phosho/total ratios based on data from (H). **J)** Glucose uptake in primary adipocytes treated with increasing BAY-3827 concentrations in the presence of MK-8722 or insulin. Data are n=8-9 from three separate experiments. **K)** Ex-vivo glucose uptake of EDL muscle treated with BAY-3827 ± MK-8722 in the presence or absence of insulin. Data is n=3-5 from four separate experiments. *p <0.05 (veh vs treatment) and ^#^p < 0.05 (veh vs veh+MK- 8722; veh vs veh +insulin). All data points shown as mean ± SEM.

**Figure S3.**
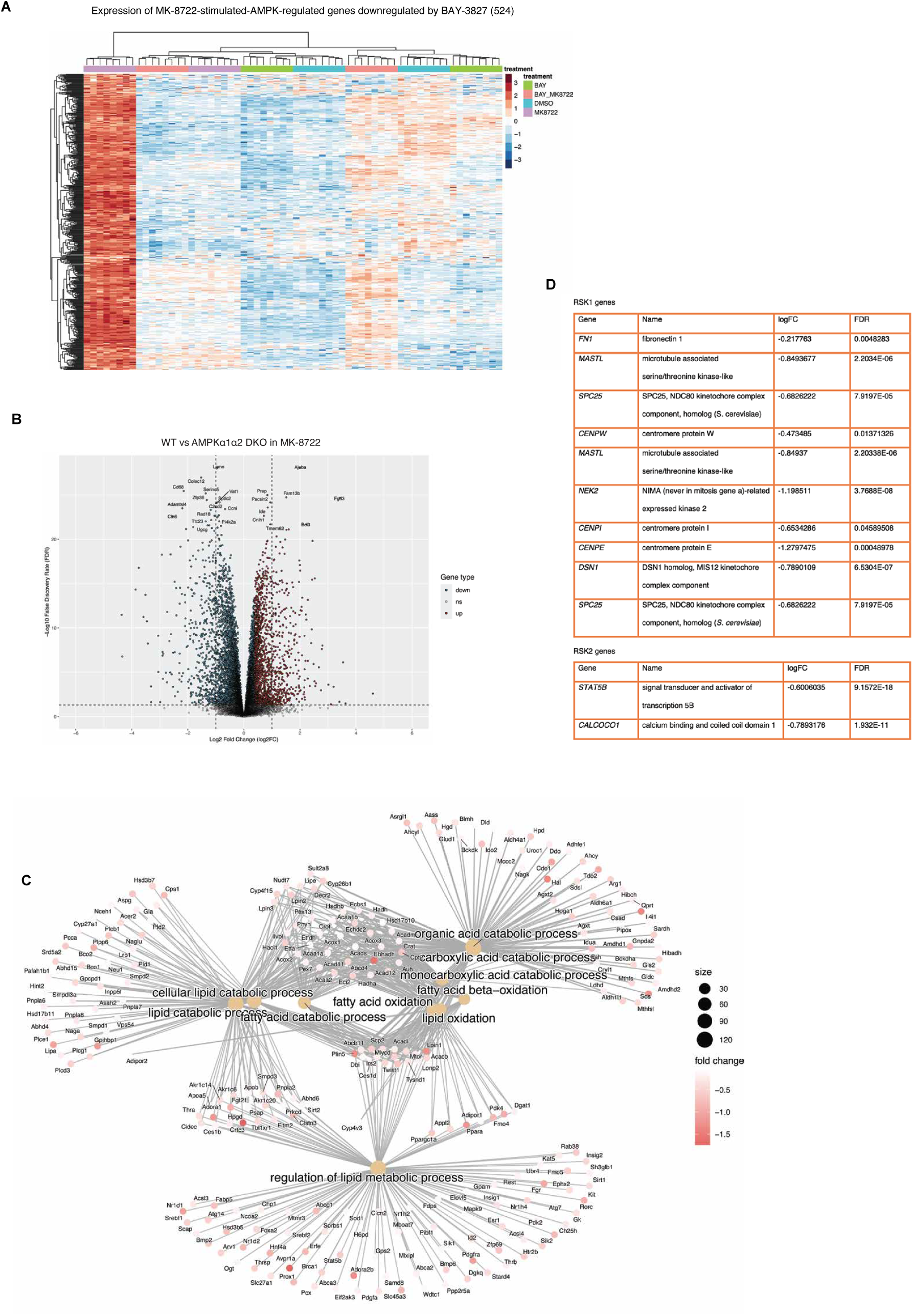
Unbiased transcriptome sequencing of AMPKα1α2 null and wild-type primary hepatocytes treated with BAY-3827 ± MK-8722. **A)** Heatmap representation of the gene expression of the 524 significant MK-8722-stimulated genes downregulated by BAY-3827 with shown treatments. **B)** Volcano plot showing top significant (FC ≥ 1.3, FDR < 0.05) upregulated (red) and downregulated (blue) genes by loss of AMPKα1α2 vs wild-type in MK-8722 treatment cells. **C)** Selected genes proposed as Ribosomal S6 kinase (RSK) isoform specific genes (*52*) downregulated in BAY-3827 in combination with MK-8722-treated wild-type hepatocytes. **D)** Cnetplot of top gene ontology biological process categories of downregulated genes by BAY-3827 + MK-8722 vs MK- 8722 alone in wild-type.

**Figure S4.**
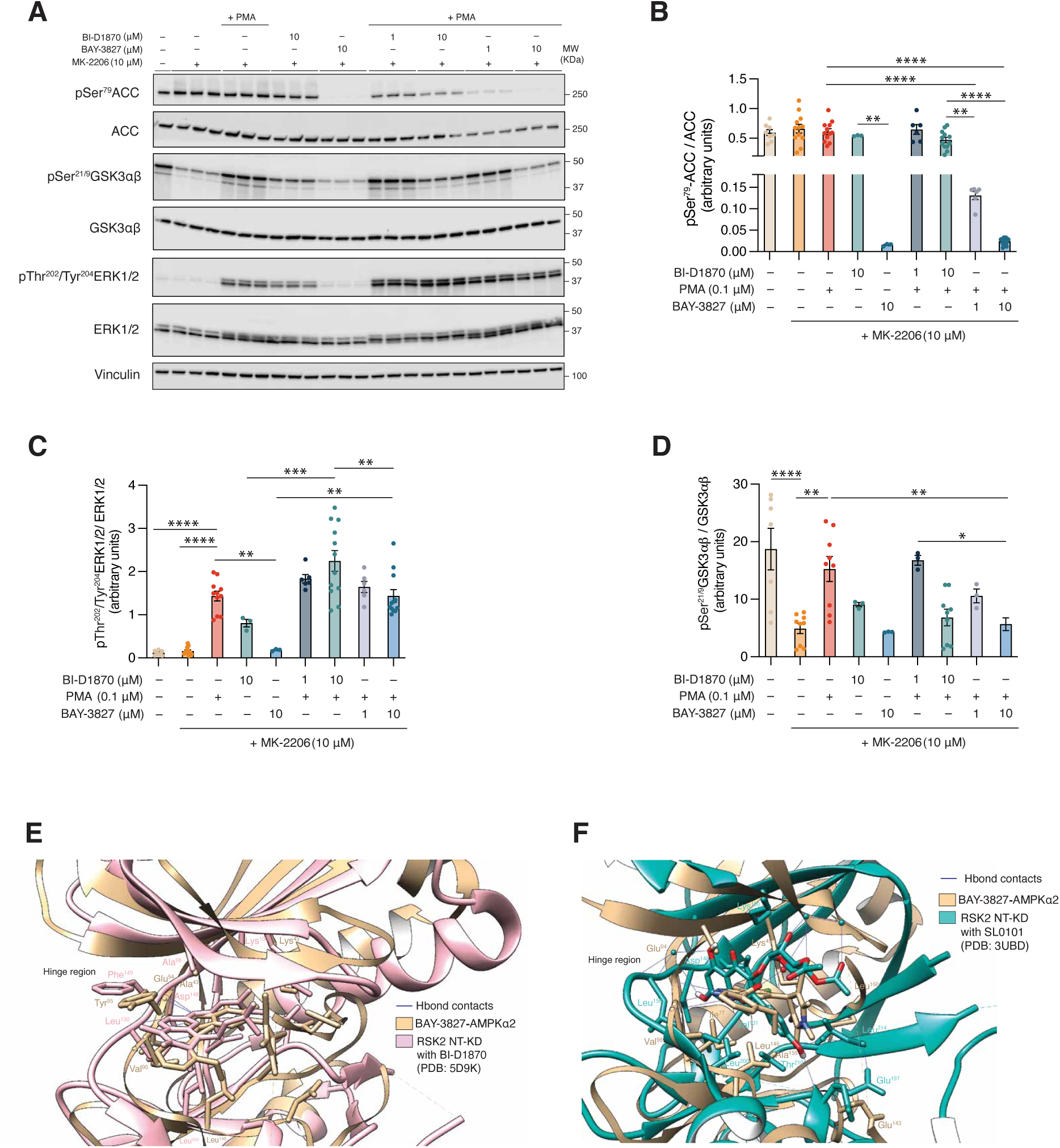
Ribosomal S6 kinase (RSK) signaling and BAY-3827. **A)** Representative western blot of RSK signaling in HEK293 cells treated with BAY-3827 or RSK inhibitor BI-D1870 in the presence of Akt inhibitor MK-2206 in the presence or absence of phorbol 12-myristate 13-acetate (PMA) ERK1/2-RSK activator. **B)** Calculated phospho/total ratios following (A) band quantification of pSer^79^-ACC **C)** pSer^21/9^GSK3αβ and **D)** pThr^202^/Tyr^204^ERK1/2. Data are n=3 from 3 independent experiments shown as mean ± SEM where *p < 0.05, **p < 0.002 and ***p < 0.0002 ****p <0.0001. **E)** RSK2 NT-KD structure solved with BI-D1870 inhibitor (PDB: 5D9K) and F) SL0101 inhibitor (PDB: 3UBD) (*55,56*) superimposed with BAY-3827-AMPKα2-KD structure showing key interacting residues, with hydrogen bonds shown in blue. Visualizations were conducted in Chimera (*78*).

**Table S1.**
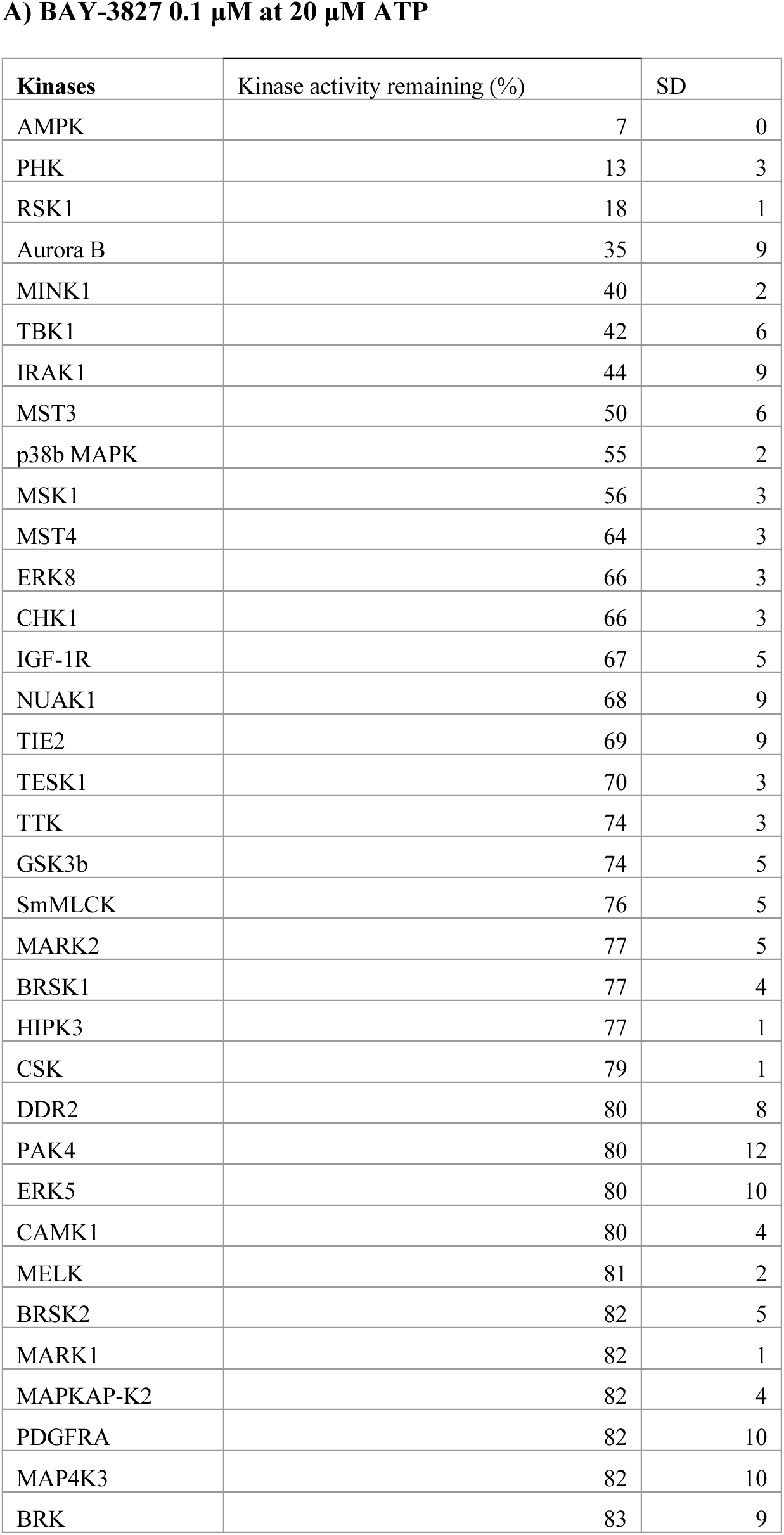
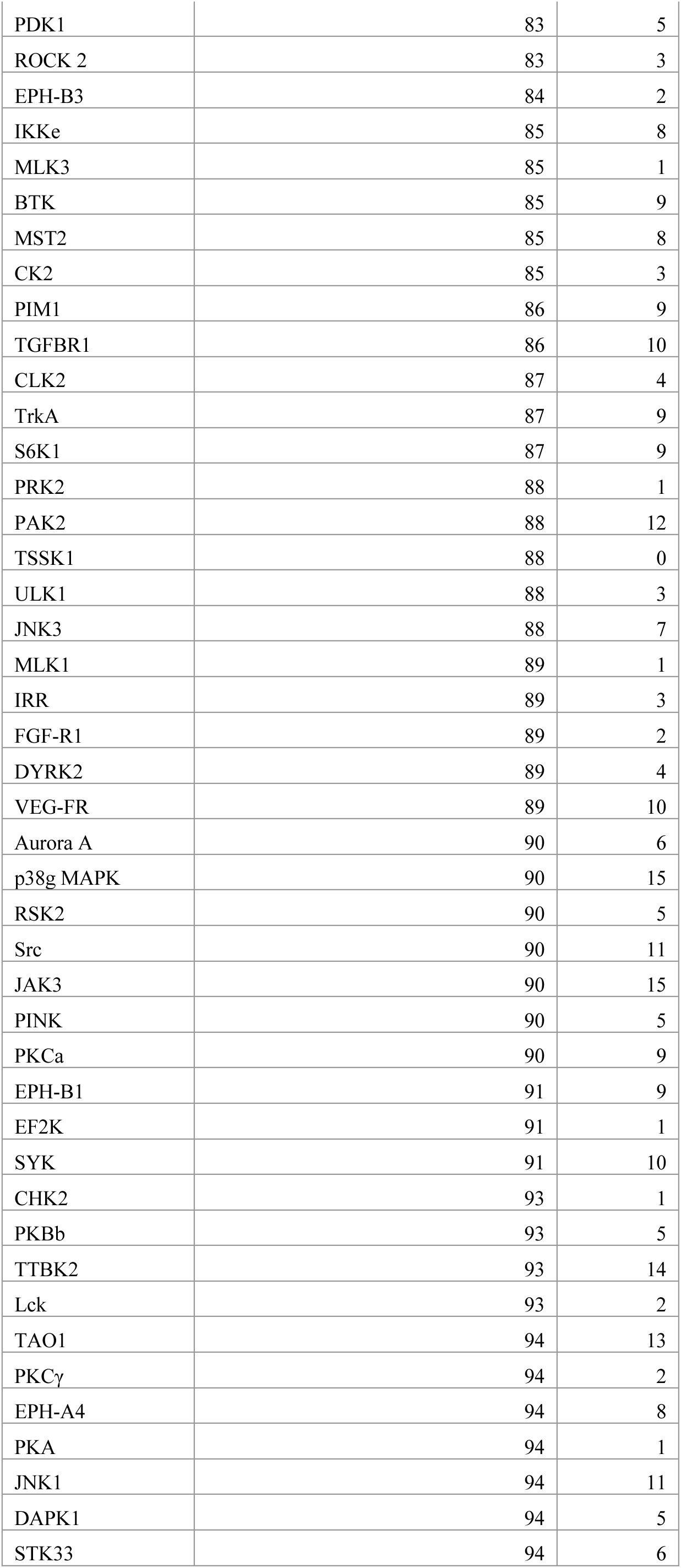
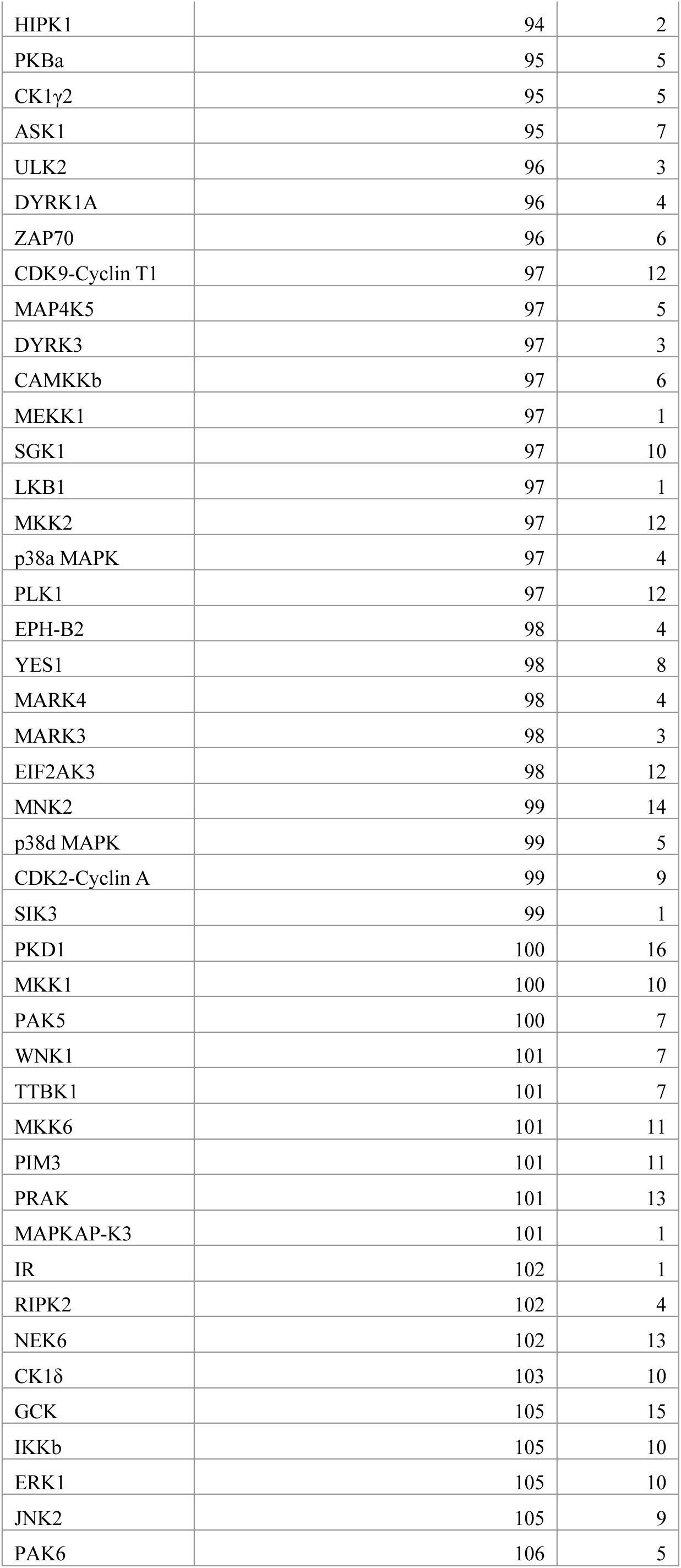
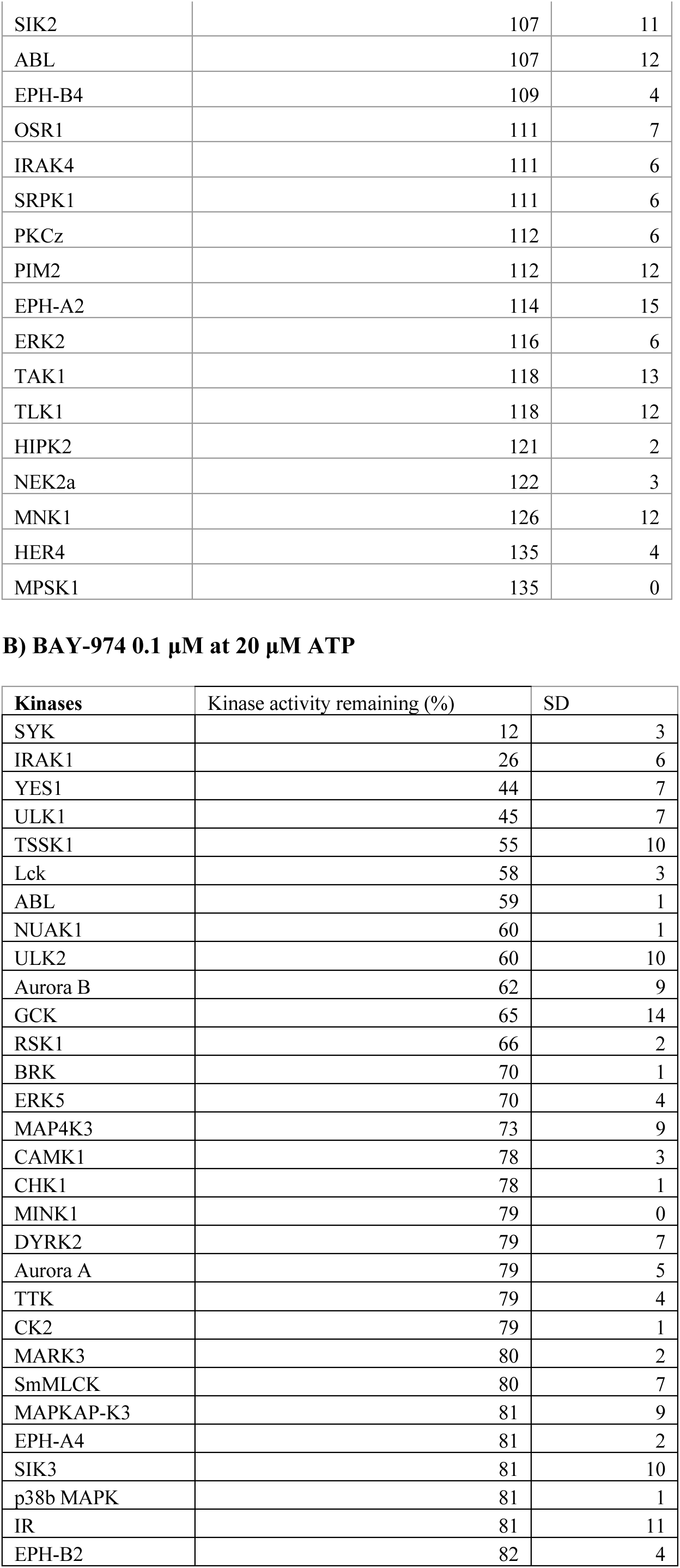
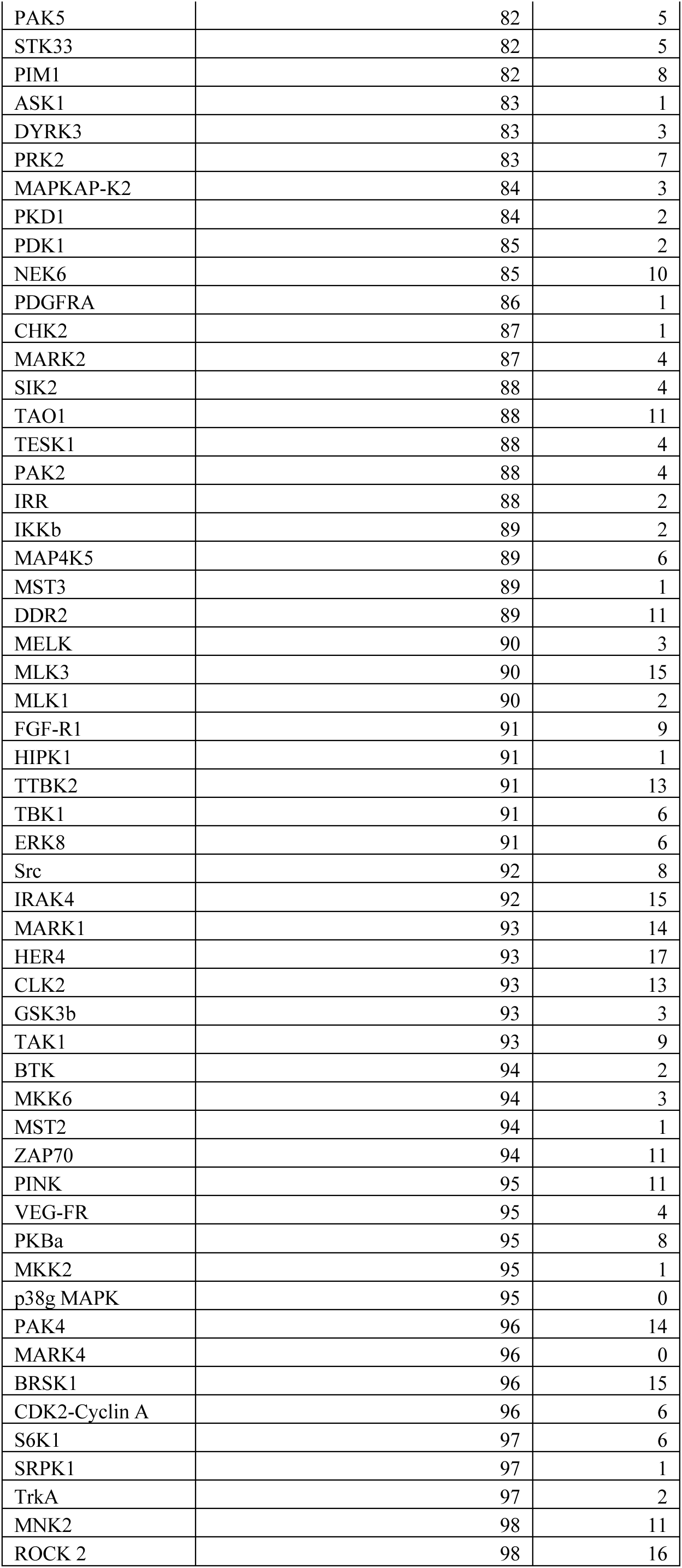
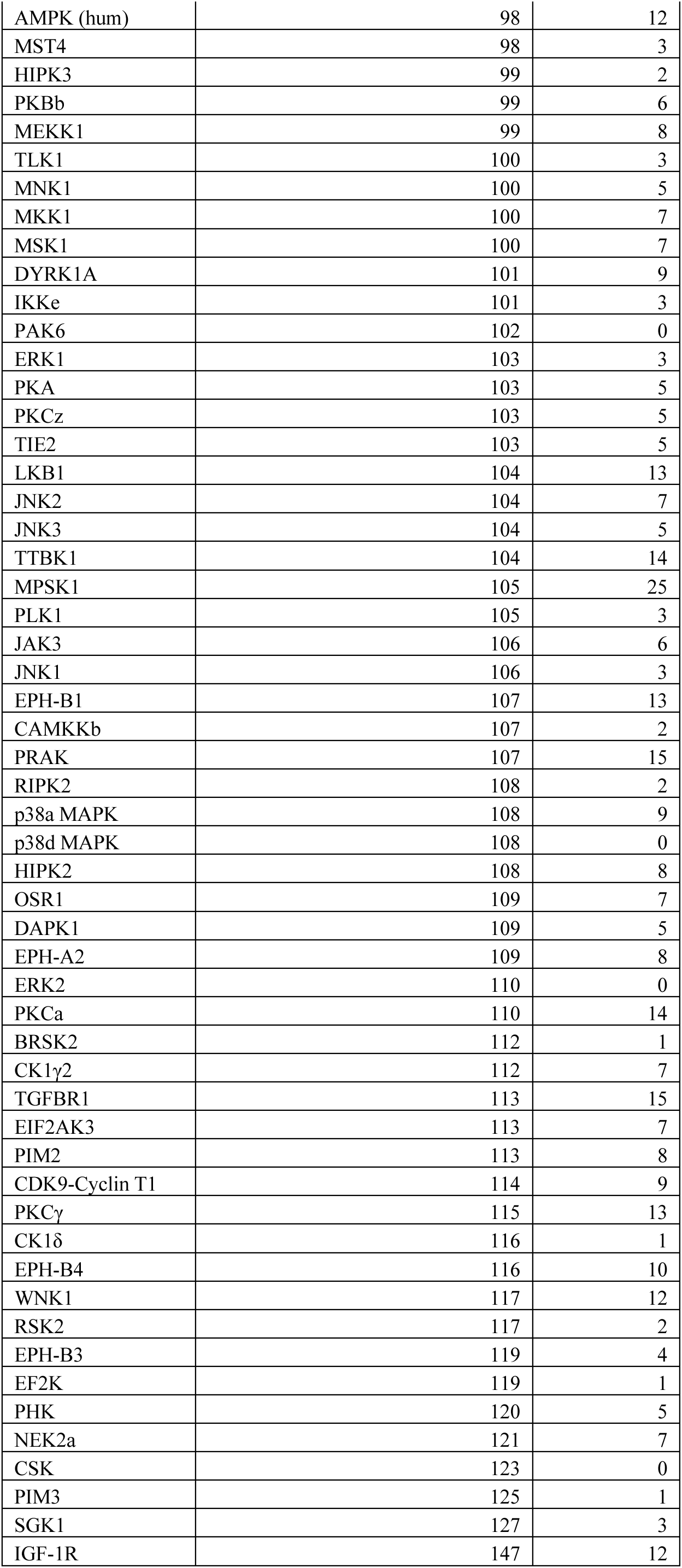
BAY-3827 (A) and BAY-974 (B) kinase selectivity data at 0.1 μM across a panel of 140 human kinases. From duplicate conditions reported as mean kinase activity remaining (%) and standard deviation (SD) values.

A) BAY-3827 0.1 μM at 20 μM ATP
| Kinases | Kinase activity remaining (%) | SD |
| --- | --- | --- |
| AMPK | 7 | 0 |
| PHK | 13 | 3 |
| RSK1 | 18 | 1 |
| Aurora B | 35 | 9 |
| MINK1 | 40 | 2 |
| TBK1 | 42 | 6 |
| IRAK1 | 44 | 9 |
| MST3 | 50 | 6 |
| p38b MAPK | 55 | 2 |
| MSK1 | 56 | 3 |
| MST4 | 64 | 3 |
| ERK8 | 66 | 3 |
| CHK1 | 66 | 3 |
| IGF-1R | 67 | 5 |
| NUAK1 | 68 | 9 |
| TIE2 | 69 | 9 |
| TESK1 | 70 | 3 |
| TTK | 74 | 3 |
| GSK3b | 74 | 5 |
| SmMLCK | 76 | 5 |
| MARK2 | 77 | 5 |
| BRSK1 | 77 | 4 |
| HIPK3 | 77 | 1 |
| CSK | 79 | 1 |
| DDR2 | 80 | 8 |
| PAK4 | 80 | 12 |
| ERK5 | 80 | 10 |
| CAMK1 | 80 | 4 |
| MELK | 81 | 2 |
| BRSK2 | 82 | 5 |
| MARK1 | 82 | 1 |
| MAPKAP-K2 | 82 | 4 |
| PDGFRA | 82 | 10 |
| MAP4K3 | 82 | 10 |
| BRK | 83 | 9 |
| PDK1 | 83 | 5 |
| ROCK 2 | 83 | 3 |
| EPH-B3 | 84 | 2 |
| IKKε | 85 | 8 |
| MLK3 | 85 | 1 |
| BTK | 85 | 9 |
| MST2 | 85 | 8 |
| CK2 | 85 | 3 |
| PIM1 | 86 | 9 |
| TGFBR1 | 86 | 10 |
| CLK2 | 87 | 4 |
| TrkA | 87 | 9 |
| S6K1 | 87 | 9 |
| PRK2 | 88 | 1 |
| PAK2 | 88 | 12 |
| TSSK1 | 88 | 0 |
| ULK1 | 88 | 3 |
| JNK3 | 88 | 7 |
| MLK1 | 89 | 1 |
| IRR | 89 | 3 |
| FGF-R1 | 89 | 2 |
| DYRK2 | 89 | 4 |
| VEG-FR | 89 | 10 |
| Aurora A | 90 | 6 |
| p38g MAPK | 90 | 15 |
| RSK2 | 90 | 5 |
| Src | 90 | 11 |
| JAK3 | 90 | 15 |
| PINK | 90 | 5 |
| PKCa | 90 | 9 |
| EPH-B1 | 91 | 9 |
| EF2K | 91 | 1 |
| SYK | 91 | 10 |
| CHK2 | 93 | 1 |
| PKBb | 93 | 5 |
| TTBK2 | 93 | 14 |
| Lck | 93 | 2 |
| TAO1 | 94 | 13 |
| PKCγ | 94 | 2 |
| EPH-A4 | 94 | 8 |
| PKA | 94 | 1 |
| JNK1 | 94 | 11 |
| DAPK1 | 94 | 5 |
| STK33 | 94 | 6 |
| HIPK1 | 94 | 2 |
| PKBa | 95 | 5 |
| CK1γ2 | 95 | 5 |
| ASK1 | 95 | 7 |
| ULK2 | 96 | 3 |
| DYRK1A | 96 | 4 |
| ZAP70 | 96 | 6 |
| CDK9-Cyclin T1 | 97 | 12 |
| MAP4K5 | 97 | 5 |
| DYRK3 | 97 | 3 |
| CAMKKb | 97 | 6 |
| MEKK1 | 97 | 1 |
| SGK1 | 97 | 10 |
| LKB1 | 97 | 1 |
| MKK2 | 97 | 12 |
| p38a MAPK | 97 | 4 |
| PLK1 | 97 | 12 |
| EPH-B2 | 98 | 4 |
| YES1 | 98 | 8 |
| MARK4 | 98 | 4 |
| MARK3 | 98 | 3 |
| EIF2AK3 | 98 | 12 |
| MNK2 | 99 | 14 |
| p38d MAPK | 99 | 5 |
| CDK2-Cyclin A | 99 | 9 |
| SIK3 | 99 | 1 |
| PKD1 | 100 | 16 |
| MKK1 | 100 | 10 |
| PAK5 | 100 | 7 |
| WNK1 | 101 | 7 |
| TTBK1 | 101 | 7 |
| MKK6 | 101 | 11 |
| PIM3 | 101 | 11 |
| PRAK | 101 | 13 |
| MAPKAP-K3 | 101 | 1 |
| IR | 102 | 1 |
| RIPK2 | 102 | 4 |
| NEK6 | 102 | 13 |
| CK1δ | 103 | 10 |
| GCK | 105 | 15 |
| IKKb | 105 | 10 |
| ERK1 | 105 | 10 |
| JNK2 | 105 | 9 |
| PAK6 | 106 | 5 |
| SIK2 | 107 | 11 |
| ABL | 107 | 12 |
| EPH-B4 | 109 | 4 |
| OSR1 | 111 | 7 |
| IRAK4 | 111 | 6 |
| SRPK1 | 111 | 6 |
| PKCz | 112 | 6 |
| PIM2 | 112 | 12 |
| EPH-A2 | 114 | 15 |
| ERK2 | 116 | 6 |
| TAK1 | 118 | 13 |
| TLK1 | 118 | 12 |
| HIPK2 | 121 | 2 |
| NEK2a | 122 | 3 |
| MNK1 | 126 | 12 |
| HER4 | 135 | 4 |
| MPSK1 | 135 | 0 |

B) BAY-974 0.1 μM at 20 μM ATP
| Kinases | Kinase activity remaining (%) | SD |
| --- | --- | --- |
| SYK | 12 | 3 |
| IRAK1 | 26 | 6 |
| YES1 | 44 | 7 |
| ULK1 | 45 | 7 |
| TSSK1 | 55 | 10 |
| Lck | 58 | 3 |
| ABL | 59 | 1 |
| NUAK1 | 60 | 1 |
| ULK2 | 60 | 10 |
| Aurora B | 62 | 9 |
| GCK | 65 | 14 |
| RSK1 | 66 | 2 |
| BRK | 70 | 1 |
| ERK5 | 70 | 4 |
| MAP4K3 | 73 | 9 |
| CAMK1 | 78 | 3 |
| CHK1 | 78 | 1 |
| MINK1 | 79 | 0 |
| DYRK2 | 79 | 7 |
| Aurora A | 79 | 5 |
| TTK | 79 | 4 |
| CK2 | 79 | 1 |
| MARK3 | 80 | 2 |
| SmMLCK | 80 | 7 |
| MAPKAP-K3 | 81 | 9 |
| EPH-A4 | 81 | 2 |
| SIK3 | 81 | 10 |
| p38b MAPK | 81 | 1 |
| IR | 81 | 11 |
| EPH-B2 | 82 | 4 |
| PAK5 | 82 | 5 |
| STK33 | 82 | 5 |
| PIM1 | 82 | 8 |
| ASK1 | 83 | 1 |
| DYRK3 | 83 | 3 |
| PRK2 | 83 | 7 |
| MAPKAP-K2 | 84 | 3 |
| PKD1 | 84 | 2 |
| PDK1 | 85 | 2 |
| NEK6 | 85 | 10 |
| PDGFRA | 86 | 1 |
| CHK2 | 87 | 1 |
| MARK2 | 87 | 4 |
| SIK2 | 88 | 4 |
| TAO1 | 88 | 11 |
| TESK1 | 88 | 4 |
| PAK2 | 88 | 4 |
| IRR | 88 | 2 |
| IKKb | 89 | 2 |
| MAP4K5 | 89 | 6 |
| MST3 | 89 | 1 |
| DDR2 | 89 | 11 |
| MELK | 90 | 3 |
| MLK3 | 90 | 15 |
| MLK1 | 90 | 2 |
| FGF-R1 | 91 | 9 |
| HIPK1 | 91 | 1 |
| TTBK2 | 91 | 13 |
| TBK1 | 91 | 6 |
| ERK8 | 91 | 6 |
| Src | 92 | 8 |
| IRAK4 | 92 | 15 |
| MARK1 | 93 | 14 |
| HER4 | 93 | 17 |
| CLK2 | 93 | 13 |
| GSK3b | 93 | 3 |
| TAK1 | 93 | 9 |
| BTK | 94 | 2 |
| MKK6 | 94 | 3 |
| MST2 | 94 | 1 |
| ZAP70 | 94 | 11 |
| PINK | 95 | 11 |
| VEG-FR | 95 | 4 |
| PKBa | 95 | 8 |
| MKK2 | 95 | 1 |
| p38g MAPK | 95 | 0 |
| PAK4 | 96 | 14 |
| MARK4 | 96 | 0 |
| BRSK1 | 96 | 15 |
| CDK2-Cyclin A | 96 | 6 |
| S6K1 | 97 | 6 |
| SRPK1 | 97 | 1 |
| TrkA | 97 | 2 |
| MNK2 | 98 | 11 |
| ROCK 2 | 98 | 16 |
| AMPK (hum) | 98 | 12 |
| MST4 | 98 | 3 |
| HIPK3 | 99 | 2 |
| PKBb | 99 | 6 |
| MEKK1 | 99 | 8 |
| TLK1 | 100 | 3 |
| MNK1 | 100 | 5 |
| MKK1 | 100 | 7 |
| MSK1 | 100 | 7 |
| DYRK1A | 101 | 9 |
| IKKe | 101 | 3 |
| PAK6 | 102 | 0 |
| ERK1 | 103 | 3 |
| PKA | 103 | 5 |
| PKCz | 103 | 5 |
| TIE2 | 103 | 5 |
| LKB1 | 104 | 13 |
| JNK2 | 104 | 7 |
| JNK3 | 104 | 5 |
| TTBK1 | 104 | 14 |
| MPSK1 | 105 | 25 |
| PLK1 | 105 | 3 |
| JAK3 | 106 | 6 |
| JNK1 | 106 | 3 |
| EPH-B1 | 107 | 13 |
| CAMKKb | 107 | 2 |
| PRAK | 107 | 15 |
| RIPK2 | 108 | 2 |
| p38a MAPK | 108 | 9 |
| p38d MAPK | 108 | 0 |
| HIPK2 | 108 | 8 |
| OSR1 | 109 | 7 |
| DAPK1 | 109 | 5 |
| EPH-A2 | 109 | 8 |
| ERK2 | 110 | 0 |
| PKCa | 110 | 14 |
| BRSK2 | 112 | 1 |
| CK1γ2 | 112 | 7 |
| TGFBR1 | 113 | 15 |
| EIF2AK3 | 113 | 7 |
| PIM2 | 113 | 8 |
| CDK9-Cyclin T1 | 114 | 9 |
| PKCγ | 115 | 13 |
| CK1δ | 116 | 1 |
| EPH-B4 | 116 | 10 |
| WNK1 | 117 | 12 |
| RSK2 | 117 | 2 |
| EPH-B3 | 119 | 4 |
| EF2K | 119 | 1 |
| PHK | 120 | 5 |
| NEK2a | 121 | 7 |
| CSK | 123 | 0 |
| PIM3 | 125 | 1 |
| SGK1 | 127 | 3 |
| IGF-1R | 147 | 12 |

**Table S2.**
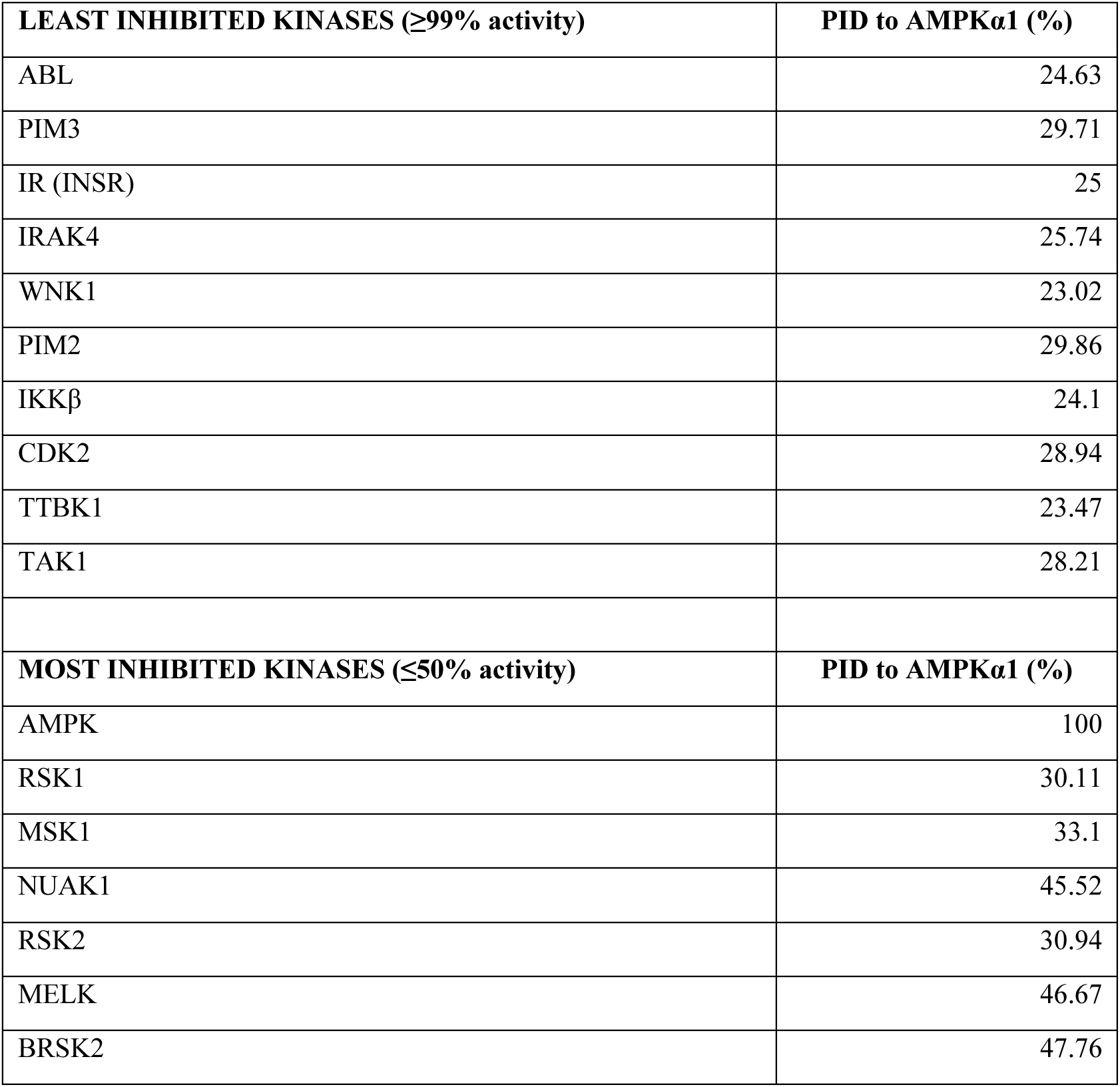
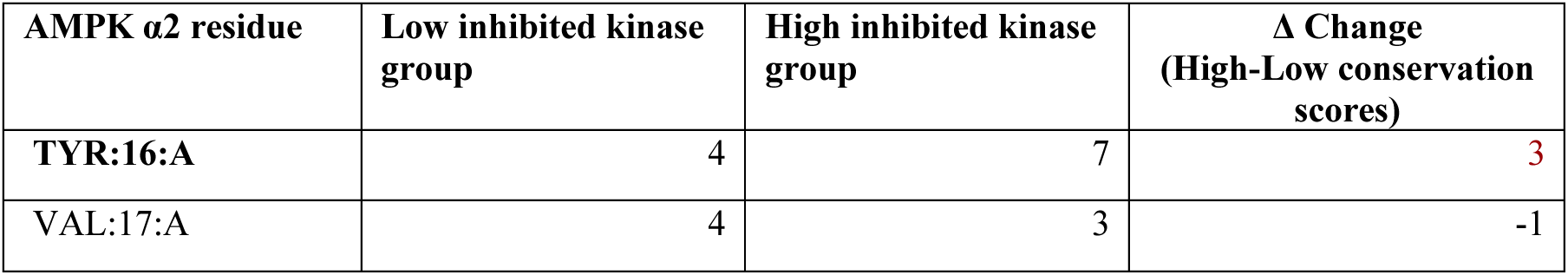
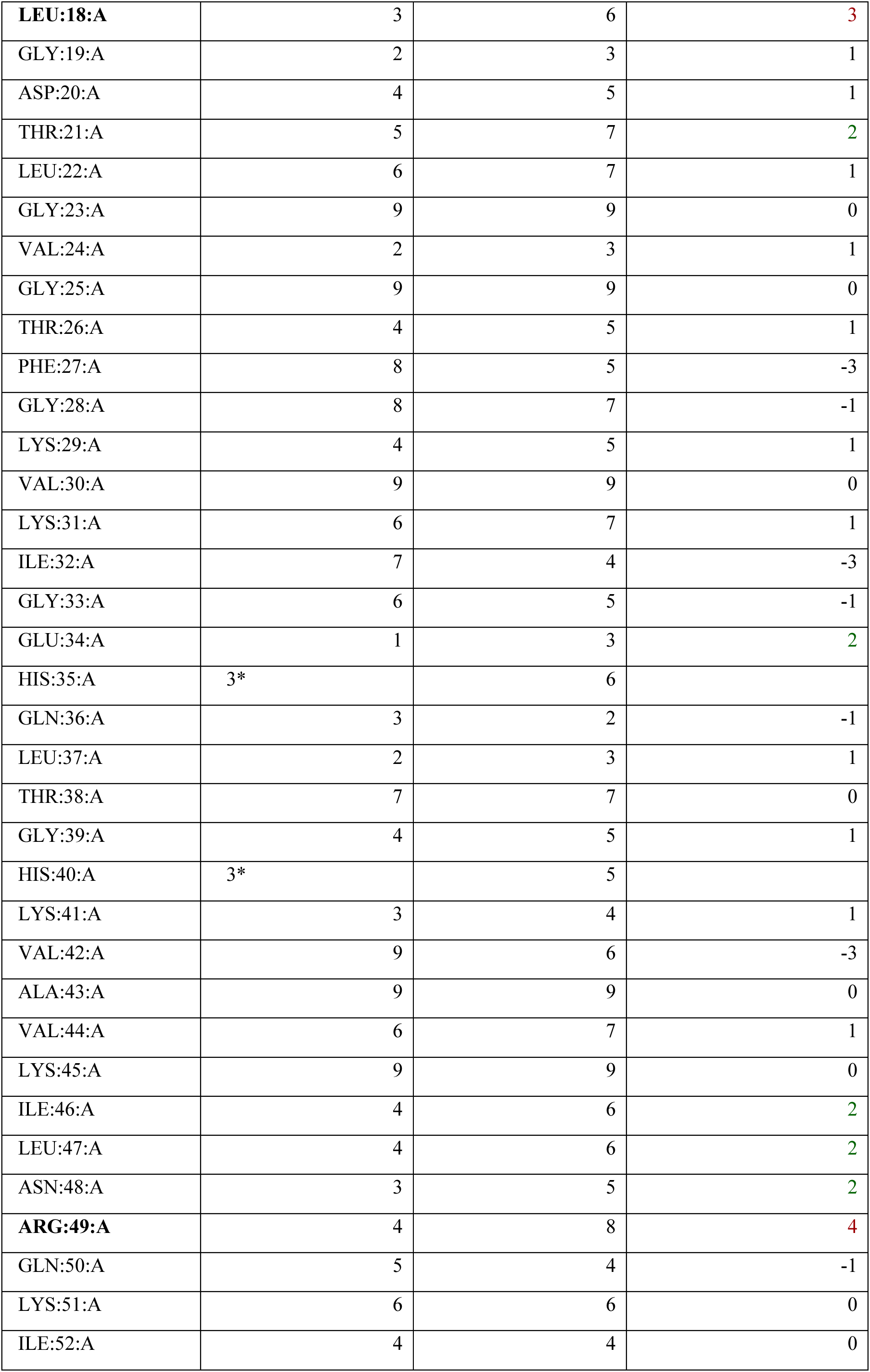
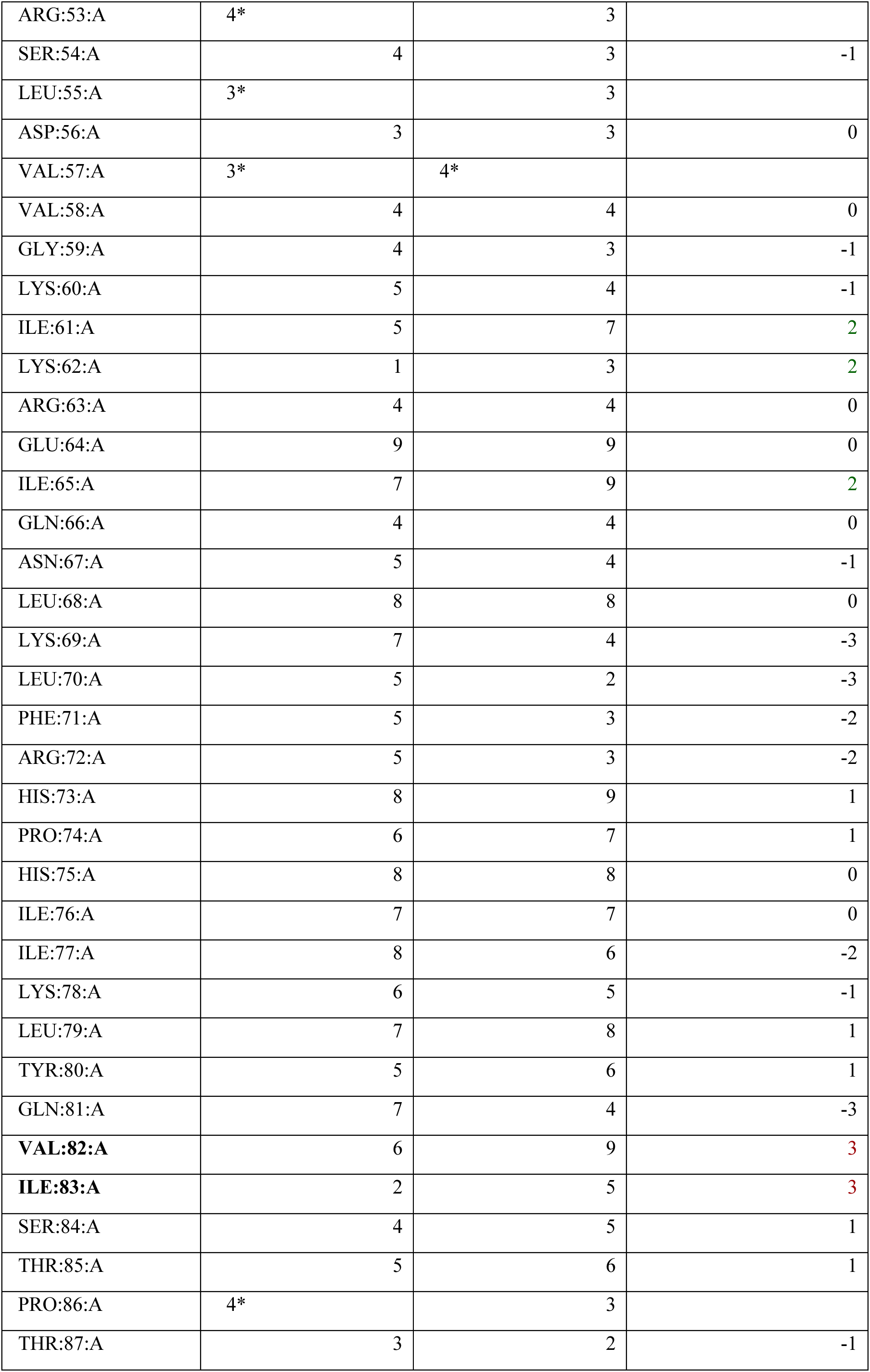
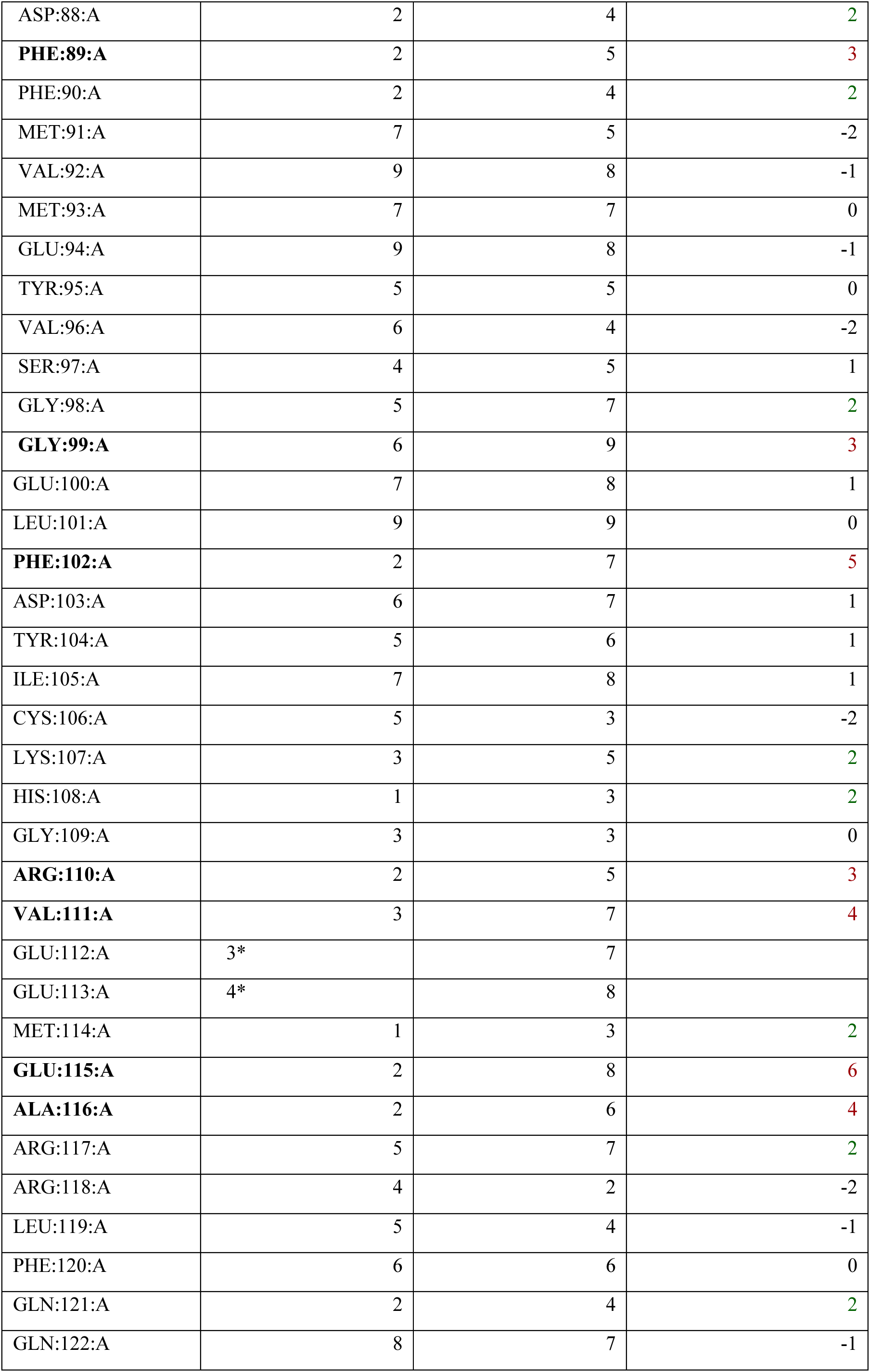
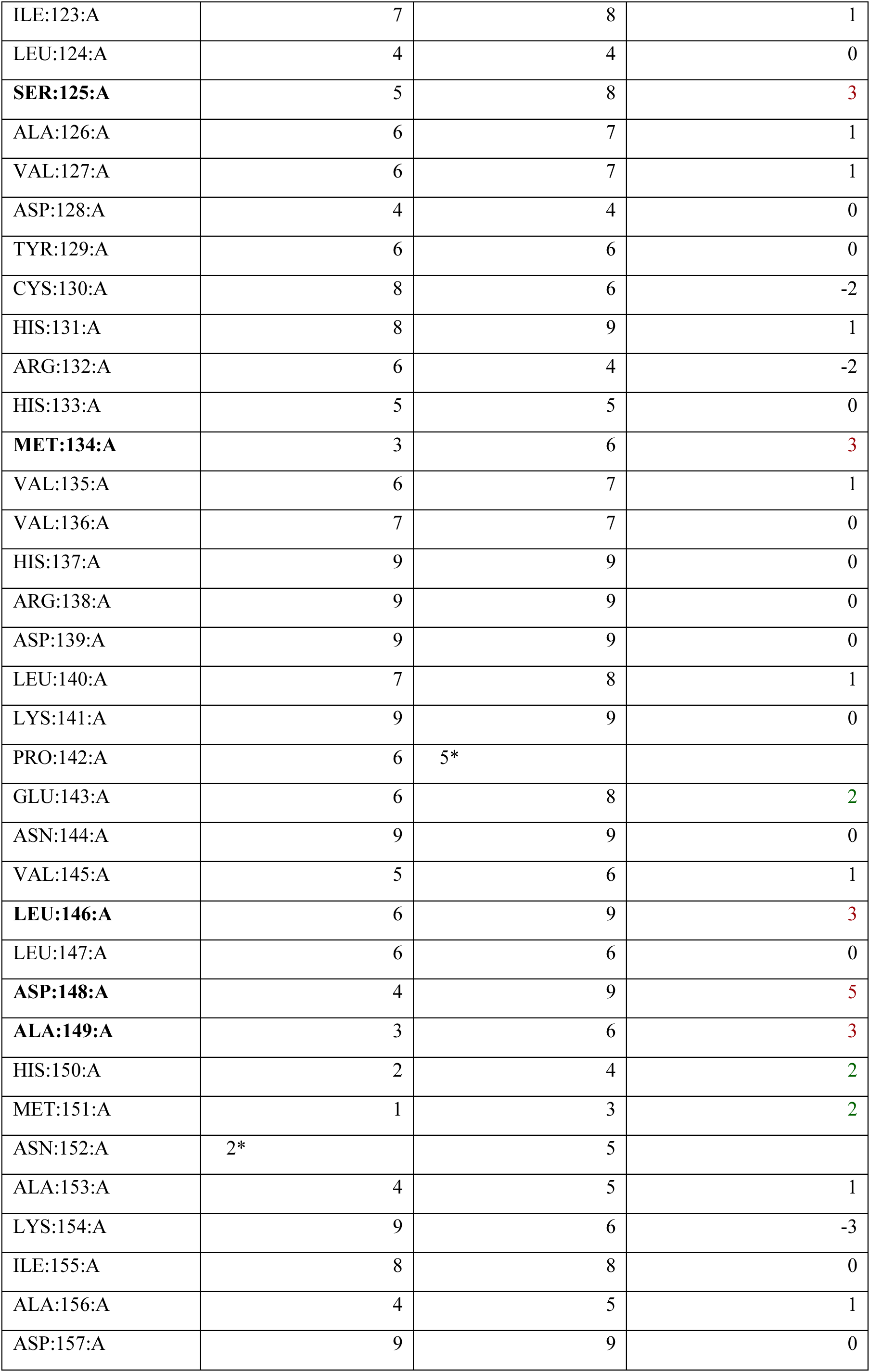
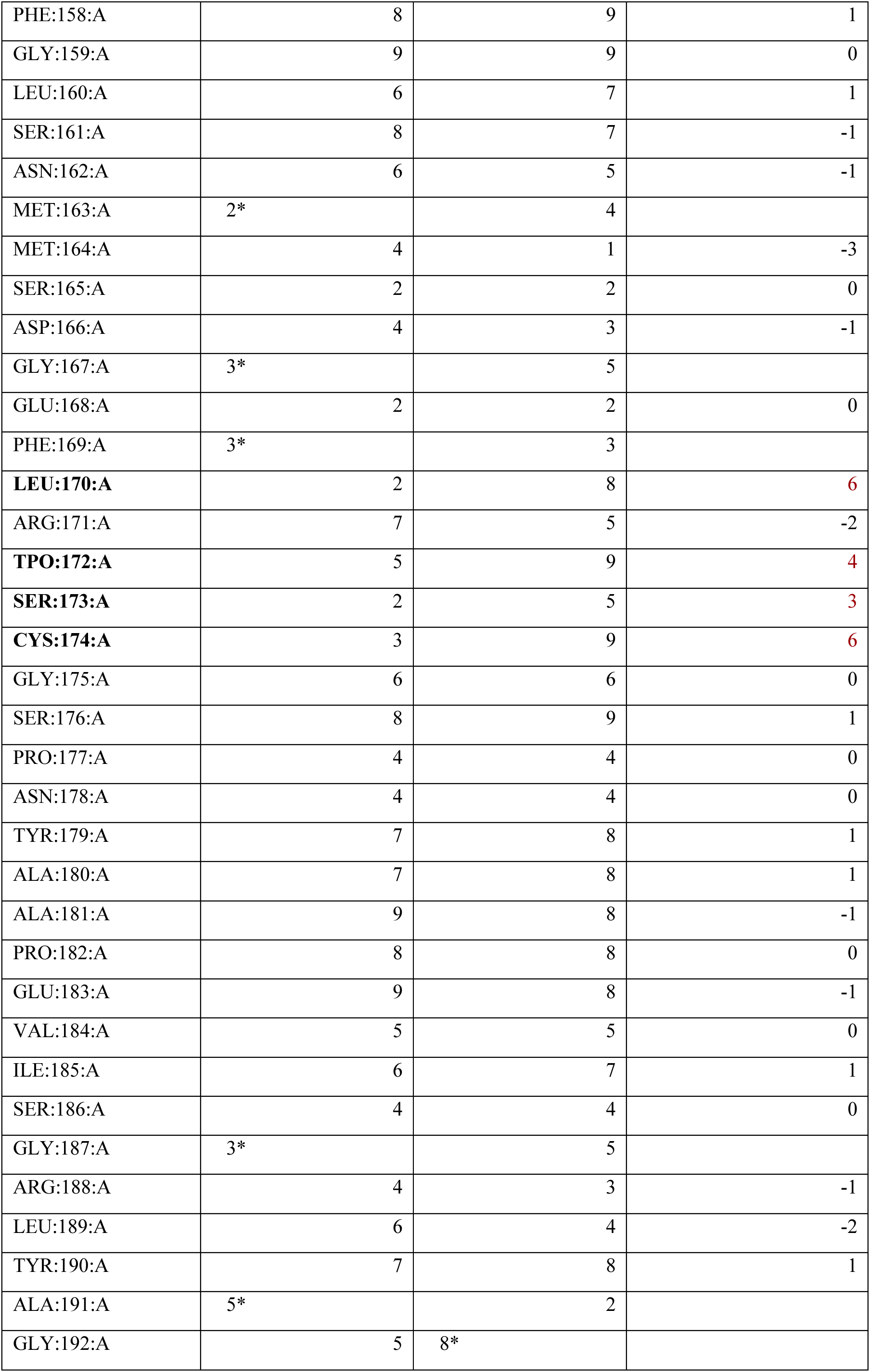
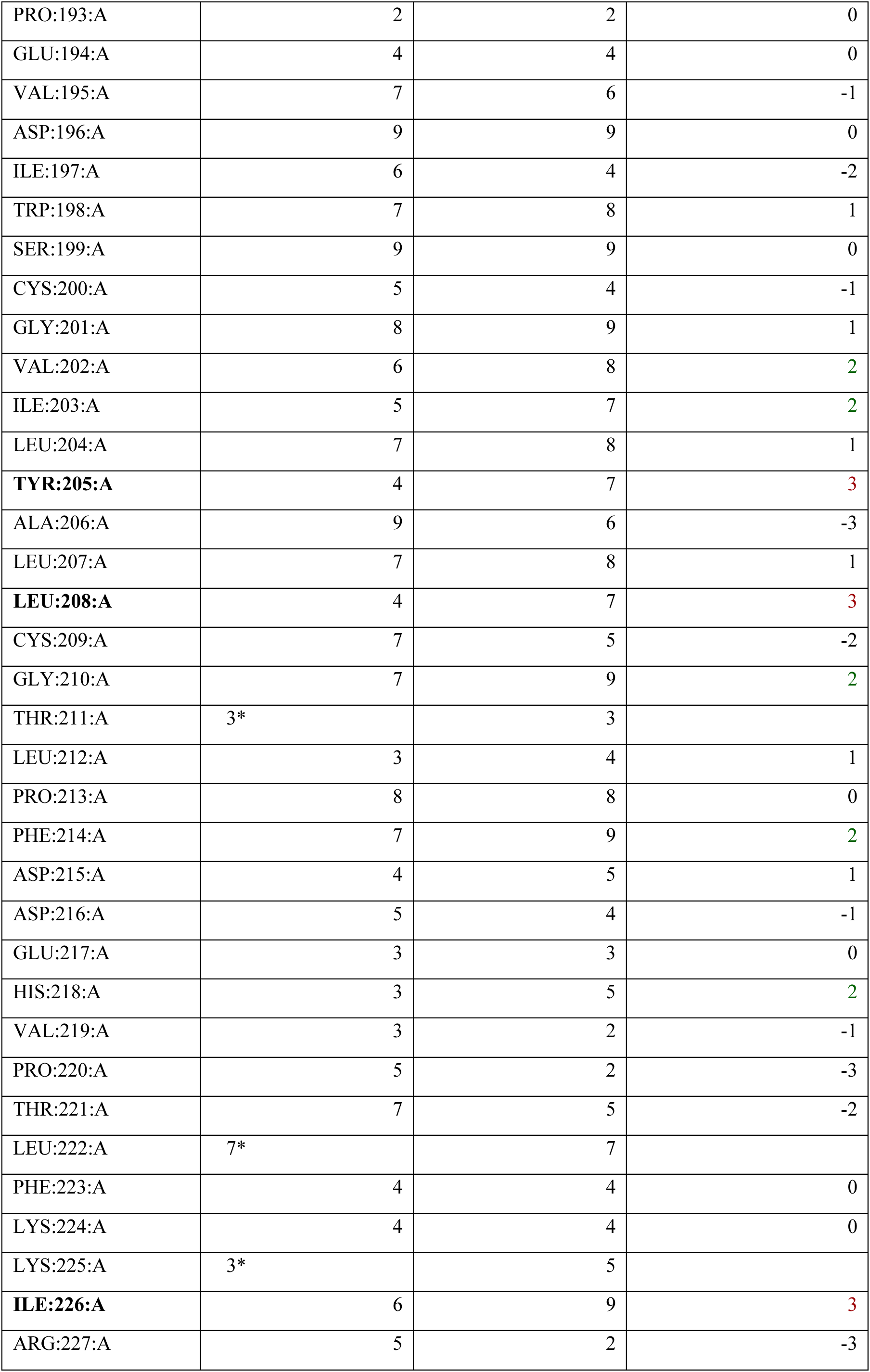
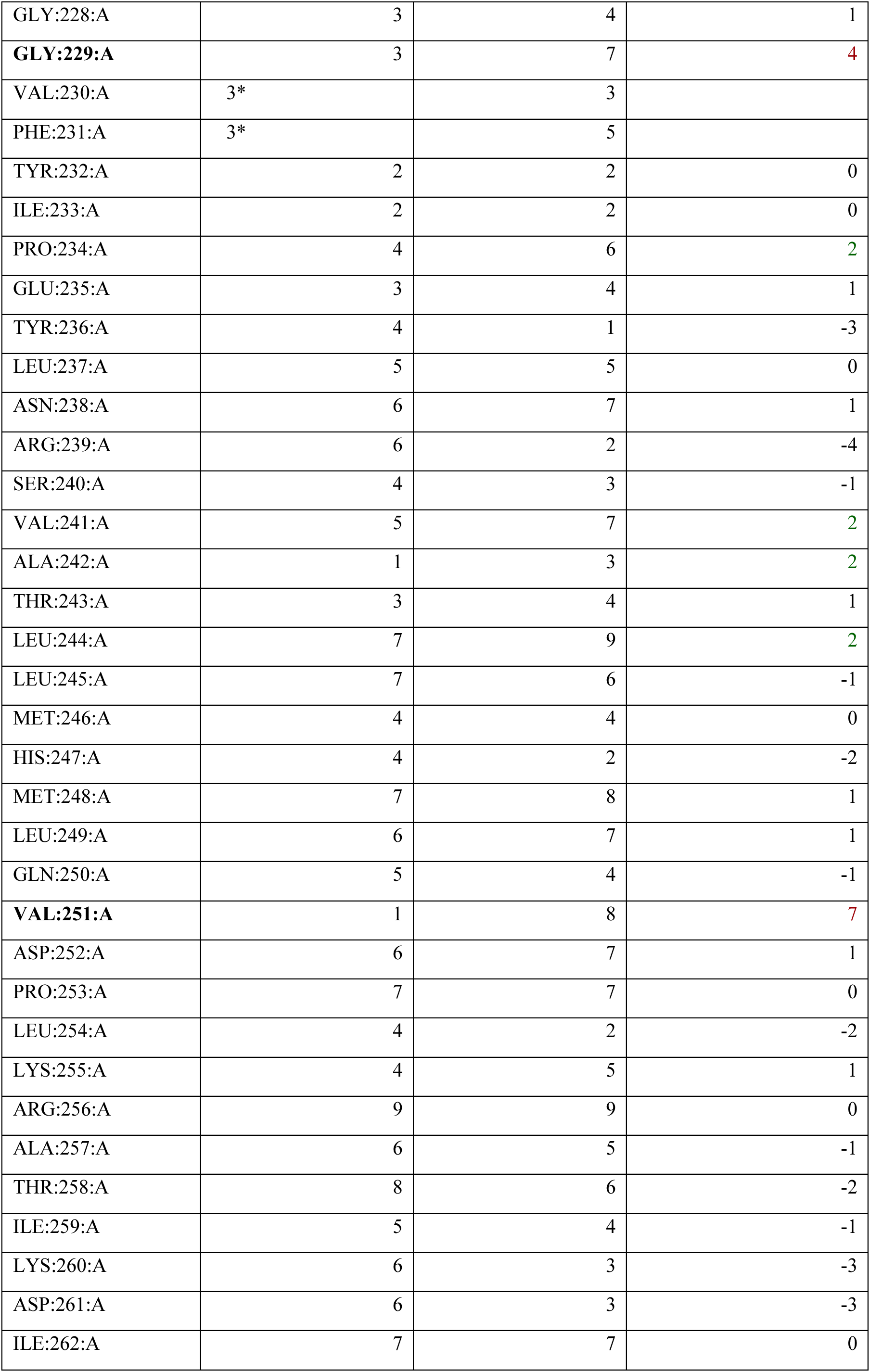
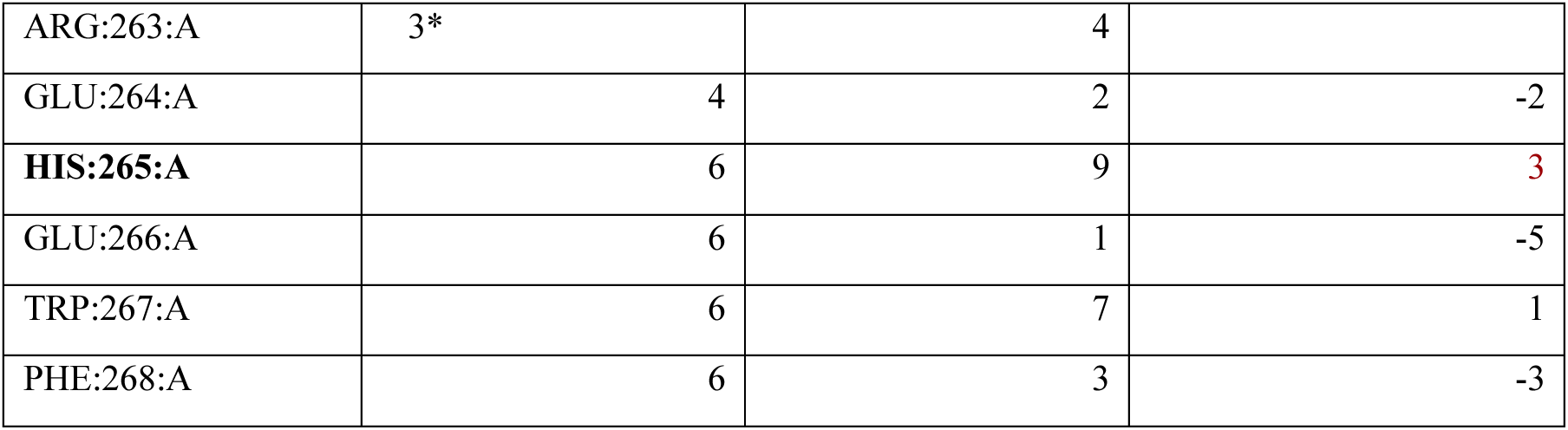
Kinase residue conservation scores calculated based sequence similarity based on in vitro kinase activity data of highly- and lowly-inhibited kinases by BAY-3827. In vitro kinase selectivity data at 1 μM was cross-referenced with past data (*37*). A) Calculated percentage identity (PID) values (%) based on a sequence alignment of selected group kinases with AMPKα1 as the reference. B) Calculated Consurf (39, 40) residue conservation scores (1-9). *low confidence scoring. A score difference between high and low-inhibited kinase groups represented by change (Δ) = 2 shown in green; ≥ 3 shown in red.

| **LEAST INHIBITED KINASES ($\geq 99\%$ activity)** | **PID to AMPKα1 (%)** |
| --- | --- |
| ABL | 24.63 |
| PIM3 | 29.71 |
| IR (INSR) | 25 |
| IRAK4 | 25.74 |
| WNK1 | 23.02 |
| PIM2 | 29.86 |
| IKKβ | 24.1 |
| CDK2 | 28.94 |
| TTBK1 | 23.47 |
| TAK1 | 28.21 |
| **MOST INHIBITED KINASES ($\leq 50\%$ activity)** | **PID to AMPKα1 (%)** |
| AMPK | 100 |
| RSK1 | 30.11 |
| MSK1 | 33.1 |
| NUAK1 | 45.52 |
| RSK2 | 30.94 |
| MELK | 46.67 |
| BRSK2 | 47.76 |

| **AMPK α2 residue** | **Low inhibited kinase group** | **High inhibited kinase group** | **$\Delta$ Change (High-Low conservation scores)** |
| --- | --- | --- | --- |
| **TYR:16:A** | 4 | 7 | 3 |
| **VAL:17:A** | 4 | 3 | -1 |

|  |  |  |  |
| --- | --- | --- | --- |
| **LEU:18:A** | 3 | 6 | 3 |
| GLY:19:A | 2 | 3 | 1 |
| ASP:20:A | 4 | 5 | 1 |
| THR:21:A | 5 | 7 | 2 |
| LEU:22:A | 6 | 7 | 1 |
| GLY:23:A | 9 | 9 | 0 |
| VAL:24:A | 2 | 3 | 1 |
| GLY:25:A | 9 | 9 | 0 |
| THR:26:A | 4 | 5 | 1 |
| PHE:27:A | 8 | 5 | -3 |
| GLY:28:A | 8 | 7 | -1 |
| LYS:29:A | 4 | 5 | 1 |
| VAL:30:A | 9 | 9 | 0 |
| LYS:31:A | 6 | 7 | 1 |
| ILE:32:A | 7 | 4 | -3 |
| GLY:33:A | 6 | 5 | -1 |
| GLU:34:A | 1 | 3 | 2 |
| HIS:35:A | 3* | 6 |  |
| GLN:36:A | 3 | 2 | -1 |
| LEU:37:A | 2 | 3 | 1 |
| THR:38:A | 7 | 7 | 0 |
| GLY:39:A | 4 | 5 | 1 |
| HIS:40:A | 3* | 5 |  |
| LYS:41:A | 3 | 4 | 1 |
| VAL:42:A | 9 | 6 | -3 |
| ALA:43:A | 9 | 9 | 0 |
| VAL:44:A | 6 | 7 | 1 |
| LYS:45:A | 9 | 9 | 0 |
| ILE:46:A | 4 | 6 | 2 |
| LEU:47:A | 4 | 6 | 2 |
| ASN:48:A | 3 | 5 | 2 |
| **ARG:49:A** | 4 | 8 | 4 |
| GLN:50:A | 5 | 4 | -1 |
| LYS:51:A | 6 | 6 | 0 |
| ILE:52:A | 4 | 4 | 0 |

|  |  |  |  |
| --- | --- | --- | --- |
| ARG:53:A | 4* | 3 |  |
| SER:54:A | 4 | 3 | -1 |
| LEU:55:A | 3* | 3 |  |
| ASP:56:A | 3 | 3 | 0 |
| VAL:57:A | 3* | 4* |  |
| VAL:58:A | 4 | 4 | 0 |
| GLY:59:A | 4 | 3 | -1 |
| LYS:60:A | 5 | 4 | -1 |
| ILE:61:A | 5 | 7 | 2 |
| LYS:62:A | 1 | 3 | 2 |
| ARG:63:A | 4 | 4 | 0 |
| GLU:64:A | 9 | 9 | 0 |
| ILE:65:A | 7 | 9 | 2 |
| GLN:66:A | 4 | 4 | 0 |
| ASN:67:A | 5 | 4 | -1 |
| LEU:68:A | 8 | 8 | 0 |
| LYS:69:A | 7 | 4 | -3 |
| LEU:70:A | 5 | 2 | -3 |
| PHE:71:A | 5 | 3 | -2 |
| ARG:72:A | 5 | 3 | -2 |
| HIS:73:A | 8 | 9 | 1 |
| PRO:74:A | 6 | 7 | 1 |
| HIS:75:A | 8 | 8 | 0 |
| ILE:76:A | 7 | 7 | 0 |
| ILE:77:A | 8 | 6 | -2 |
| LYS:78:A | 6 | 5 | -1 |
| LEU:79:A | 7 | 8 | 1 |
| TYR:80:A | 5 | 6 | 1 |
| GLN:81:A | 7 | 4 | -3 |
| **VAL:82:A** | 6 | 9 | 3 |
| **ILE:83:A** | 2 | 5 | 3 |
| SER:84:A | 4 | 5 | 1 |
| THR:85:A | 5 | 6 | 1 |
| PRO:86:A | 4* | 3 |  |
| THR:87:A | 3 | 2 | -1 |

|  |  |  |  |
| --- | --- | --- | --- |
| ASP:88:A | 2 | 4 | 2 |
| **PHE:89:A** | 2 | 5 | 3 |
| PHE:90:A | 2 | 4 | 2 |
| MET:91:A | 7 | 5 | -2 |
| VAL:92:A | 9 | 8 | -1 |
| MET:93:A | 7 | 7 | 0 |
| GLU:94:A | 9 | 8 | -1 |
| TYR:95:A | 5 | 5 | 0 |
| VAL:96:A | 6 | 4 | -2 |
| SER:97:A | 4 | 5 | 1 |
| GLY:98:A | 5 | 7 | 2 |
| **GLY:99:A** | 6 | 9 | 3 |
| GLU:100:A | 7 | 8 | 1 |
| LEU:101:A | 9 | 9 | 0 |
| **PHE:102:A** | 2 | 7 | 5 |
| ASP:103:A | 6 | 7 | 1 |
| TYR:104:A | 5 | 6 | 1 |
| ILE:105:A | 7 | 8 | 1 |
| CYS:106:A | 5 | 3 | -2 |
| LYS:107:A | 3 | 5 | 2 |
| HIS:108:A | 1 | 3 | 2 |
| GLY:109:A | 3 | 3 | 0 |
| **ARG:110:A** | 2 | 5 | 3 |
| **VAL:111:A** | 3 | 7 | 4 |
| GLU:112:A | 3* | 7 |  |
| GLU:113:A | 4* | 8 |  |
| MET:114:A | 1 | 3 | 2 |
| **GLU:115:A** | 2 | 8 | 6 |
| **ALA:116:A** | 2 | 6 | 4 |
| ARG:117:A | 5 | 7 | 2 |
| ARG:118:A | 4 | 2 | -2 |
| LEU:119:A | 5 | 4 | -1 |
| PHE:120:A | 6 | 6 | 0 |
| GLN:121:A | 2 | 4 | 2 |
| GLN:122:A | 8 | 7 | -1 |

|  |  |  |  |
| --- | --- | --- | --- |
| ILE:123:A | 7 | 8 | 1 |
| LEU:124:A | 4 | 4 | 0 |
| **SER:125:A** | 5 | 8 | 3 |
| ALA:126:A | 6 | 7 | 1 |
| VAL:127:A | 6 | 7 | 1 |
| ASP:128:A | 4 | 4 | 0 |
| TYR:129:A | 6 | 6 | 0 |
| CYS:130:A | 8 | 6 | -2 |
| HIS:131:A | 8 | 9 | 1 |
| ARG:132:A | 6 | 4 | -2 |
| HIS:133:A | 5 | 5 | 0 |
| **MET:134:A** | 3 | 6 | 3 |
| VAL:135:A | 6 | 7 | 1 |
| VAL:136:A | 7 | 7 | 0 |
| HIS:137:A | 9 | 9 | 0 |
| ARG:138:A | 9 | 9 | 0 |
| ASP:139:A | 9 | 9 | 0 |
| LEU:140:A | 7 | 8 | 1 |
| LYS:141:A | 9 | 9 | 0 |
| PRO:142:A | 6 | 5* |  |
| GLU:143:A | 6 | 8 | 2 |
| ASN:144:A | 9 | 9 | 0 |
| VAL:145:A | 5 | 6 | 1 |
| **LEU:146:A** | 6 | 9 | 3 |
| LEU:147:A | 6 | 6 | 0 |
| **ASP:148:A** | 4 | 9 | 5 |
| **ALA:149:A** | 3 | 6 | 3 |
| HIS:150:A | 2 | 4 | 2 |
| MET:151:A | 1 | 3 | 2 |
| ASN:152:A | 2* | 5 |  |
| ALA:153:A | 4 | 5 | 1 |
| LYS:154:A | 9 | 6 | -3 |
| ILE:155:A | 8 | 8 | 0 |
| ALA:156:A | 4 | 5 | 1 |
| ASP:157:A | 9 | 9 | 0 |

|  |  |  |  |
| --- | --- | --- | --- |
| PHE:158:A | 8 | 9 | 1 |
| GLY:159:A | 9 | 9 | 0 |
| LEU:160:A | 6 | 7 | 1 |
| SER:161:A | 8 | 7 | -1 |
| ASN:162:A | 6 | 5 | -1 |
| MET:163:A | 2* | 4 |  |
| MET:164:A | 4 | 1 | -3 |
| SER:165:A | 2 | 2 | 0 |
| ASP:166:A | 4 | 3 | -1 |
| GLY:167:A | 3* | 5 |  |
| GLU:168:A | 2 | 2 | 0 |
| PHE:169:A | 3* | 3 |  |
| **LEU:170:A** | 2 | 8 | **6** |
| ARG:171:A | 7 | 5 | -2 |
| **TPO:172:A** | 5 | 9 | **4** |
| **SER:173:A** | 2 | 5 | **3** |
| **CYS:174:A** | 3 | 9 | **6** |
| GLY:175:A | 6 | 6 | 0 |
| SER:176:A | 8 | 9 | 1 |
| PRO:177:A | 4 | 4 | 0 |
| ASN:178:A | 4 | 4 | 0 |
| TYR:179:A | 7 | 8 | 1 |
| ALA:180:A | 7 | 8 | 1 |
| ALA:181:A | 9 | 8 | -1 |
| PRO:182:A | 8 | 8 | 0 |
| GLU:183:A | 9 | 8 | -1 |
| VAL:184:A | 5 | 5 | 0 |
| ILE:185:A | 6 | 7 | 1 |
| SER:186:A | 4 | 4 | 0 |
| GLY:187:A | 3* | 5 |  |
| ARG:188:A | 4 | 3 | -1 |
| LEU:189:A | 6 | 4 | -2 |
| TYR:190:A | 7 | 8 | 1 |
| ALA:191:A | 5* | 2 |  |
| GLY:192:A | 5 | 8* |  |

|  |  |  |  |
| --- | --- | --- | --- |
| PRO:193:A | 2 | 2 | 0 |
| GLU:194:A | 4 | 4 | 0 |
| VAL:195:A | 7 | 6 | -1 |
| ASP:196:A | 9 | 9 | 0 |
| ILE:197:A | 6 | 4 | -2 |
| TRP:198:A | 7 | 8 | 1 |
| SER:199:A | 9 | 9 | 0 |
| CYS:200:A | 5 | 4 | -1 |
| GLY:201:A | 8 | 9 | 1 |
| VAL:202:A | 6 | 8 | 2 |
| ILE:203:A | 5 | 7 | 2 |
| LEU:204:A | 7 | 8 | 1 |
| **TYR:205:A** | 4 | 7 | 3 |
| ALA:206:A | 9 | 6 | -3 |
| LEU:207:A | 7 | 8 | 1 |
| **LEU:208:A** | 4 | 7 | 3 |
| CYS:209:A | 7 | 5 | -2 |
| GLY:210:A | 7 | 9 | 2 |
| THR:211:A | 3* | 3 |  |
| LEU:212:A | 3 | 4 | 1 |
| PRO:213:A | 8 | 8 | 0 |
| PHE:214:A | 7 | 9 | 2 |
| ASP:215:A | 4 | 5 | 1 |
| ASP:216:A | 5 | 4 | -1 |
| GLU:217:A | 3 | 3 | 0 |
| HIS:218:A | 3 | 5 | 2 |
| VAL:219:A | 3 | 2 | -1 |
| PRO:220:A | 5 | 2 | -3 |
| THR:221:A | 7 | 5 | -2 |
| LEU:222:A | 7* | 7 |  |
| PHE:223:A | 4 | 4 | 0 |
| LYS:224:A | 4 | 4 | 0 |
| LYS:225:A | 3* | 5 |  |
| **ILE:226:A** | 6 | 9 | 3 |
| ARG:227:A | 5 | 2 | -3 |

|  |  |  |  |
| --- | --- | --- | --- |
| GLY:228:A | 3 | 4 | 1 |
| **GLY:229:A** | 3 | 7 | 4 |
| VAL:230:A | 3* | 3 |  |
| PHE:231:A | 3* | 5 |  |
| TYR:232:A | 2 | 2 | 0 |
| ILE:233:A | 2 | 2 | 0 |
| PRO:234:A | 4 | 6 | 2 |
| GLU:235:A | 3 | 4 | 1 |
| TYR:236:A | 4 | 1 | -3 |
| LEU:237:A | 5 | 5 | 0 |
| ASN:238:A | 6 | 7 | 1 |
| ARG:239:A | 6 | 2 | -4 |
| SER:240:A | 4 | 3 | -1 |
| VAL:241:A | 5 | 7 | 2 |
| ALA:242:A | 1 | 3 | 2 |
| THR:243:A | 3 | 4 | 1 |
| LEU:244:A | 7 | 9 | 2 |
| LEU:245:A | 7 | 6 | -1 |
| MET:246:A | 4 | 4 | 0 |
| HIS:247:A | 4 | 2 | -2 |
| MET:248:A | 7 | 8 | 1 |
| LEU:249:A | 6 | 7 | 1 |
| GLN:250:A | 5 | 4 | -1 |
| **VAL:251:A** | 1 | 8 | 7 |
| ASP:252:A | 6 | 7 | 1 |
| PRO:253:A | 7 | 7 | 0 |
| LEU:254:A | 4 | 2 | -2 |
| LYS:255:A | 4 | 5 | 1 |
| ARG:256:A | 9 | 9 | 0 |
| ALA:257:A | 6 | 5 | -1 |
| THR:258:A | 8 | 6 | -2 |
| ILE:259:A | 5 | 4 | -1 |
| LYS:260:A | 6 | 3 | -3 |
| ASP:261:A | 6 | 3 | -3 |
| ILE:262:A | 7 | 7 | 0 |

|  |  |  |  |
| --- | --- | --- | --- |
| ARG:263:A | 3* | 4 |  |
| GLU:264:A | 4 | 2 | -2 |
| **HIS:265:A** | 6 | 9 | **3** |
| GLU:266:A | 6 | 1 | -5 |
| TRP:267:A | 6 | 7 | 1 |
| PHE:268:A | 6 | 3 | -3 |

**Table S3.**
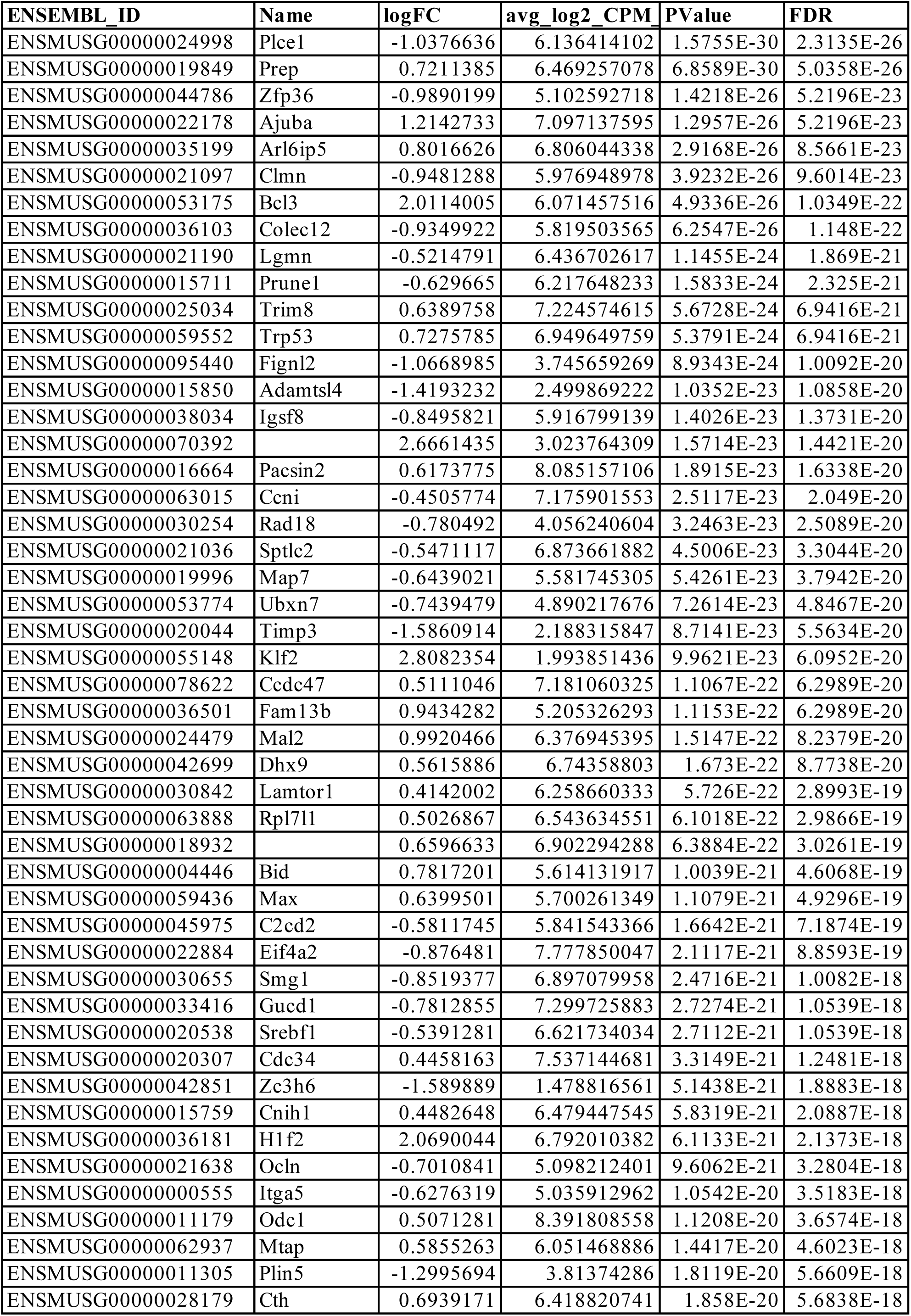
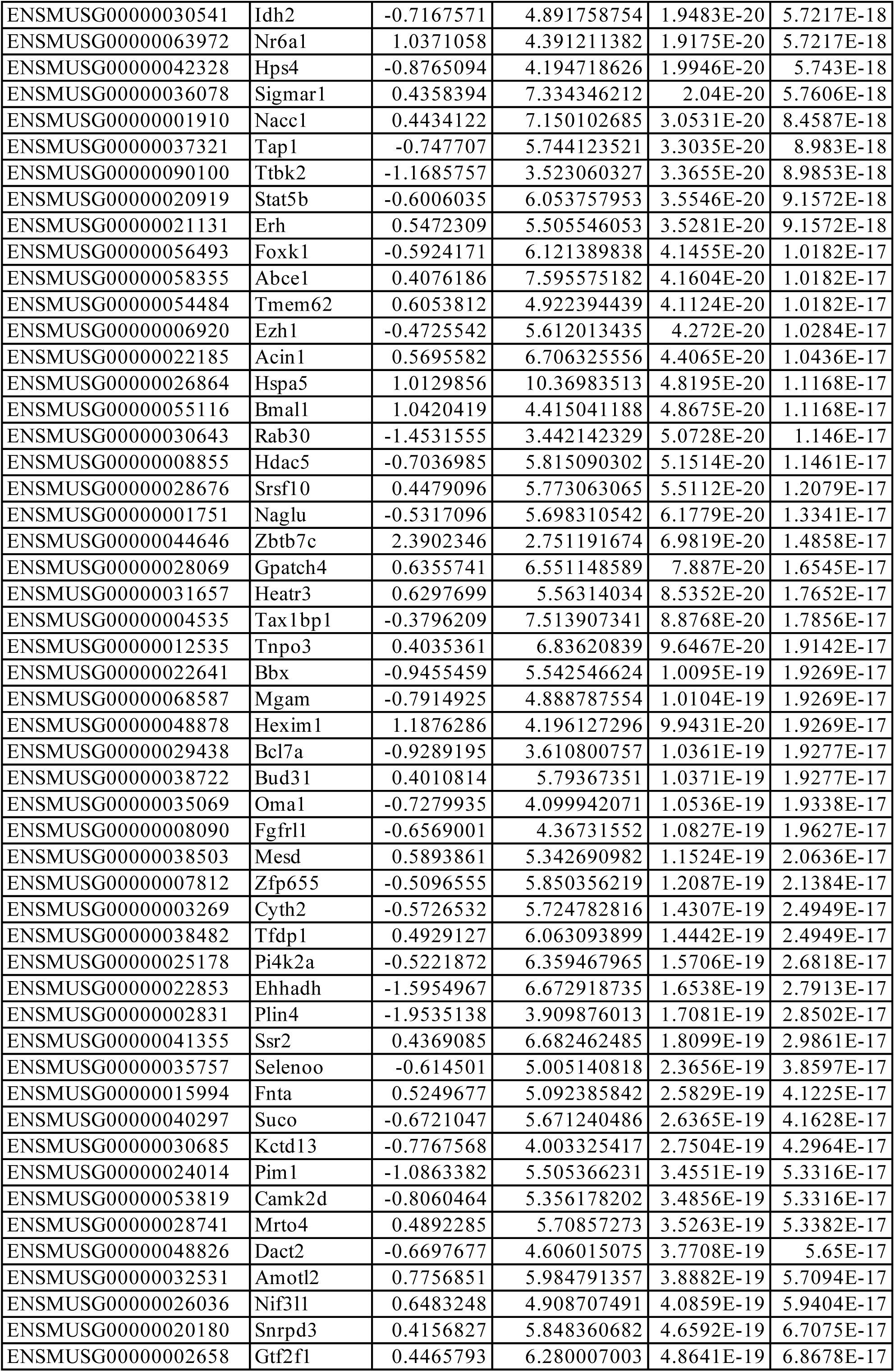
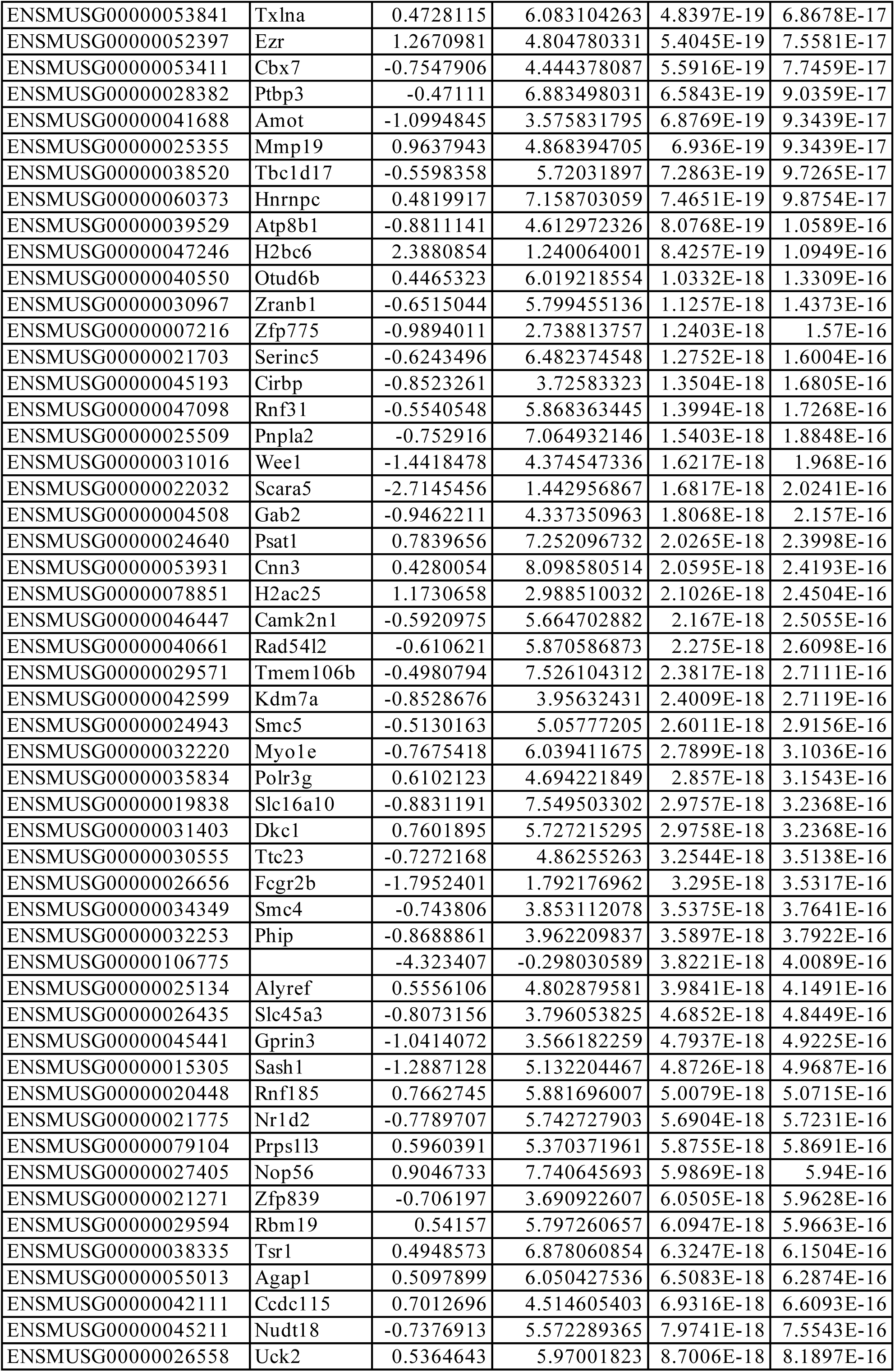
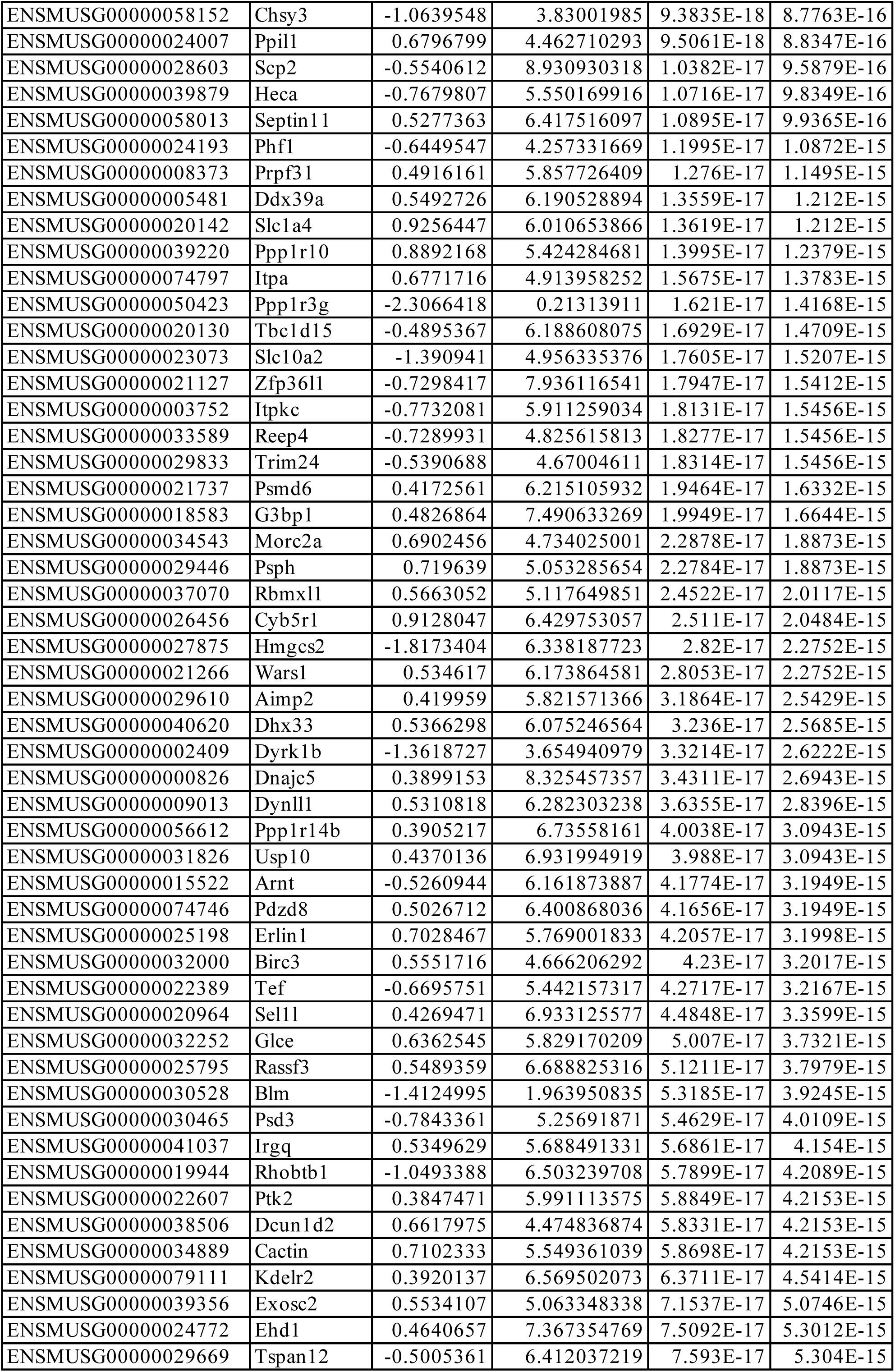
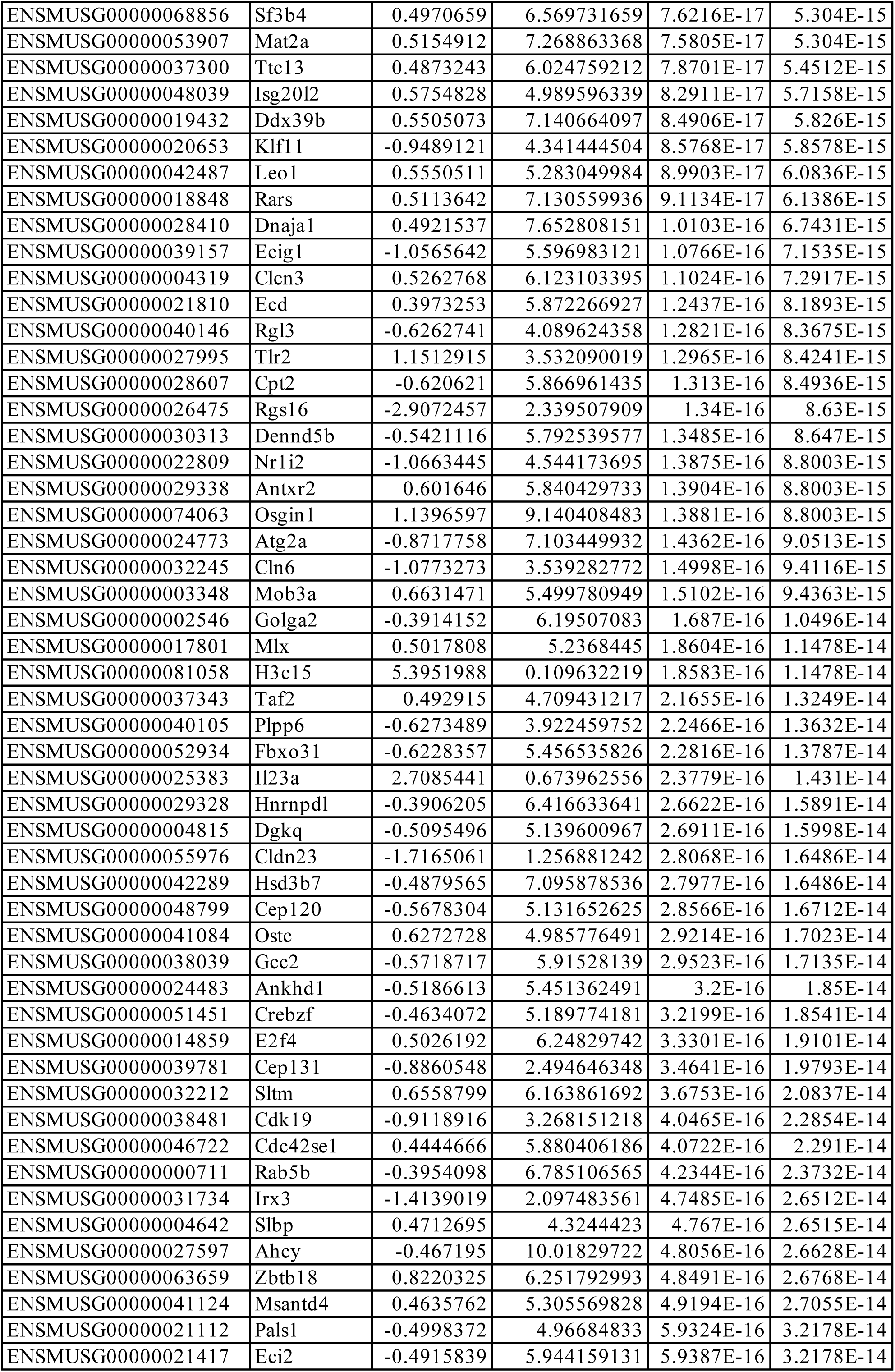
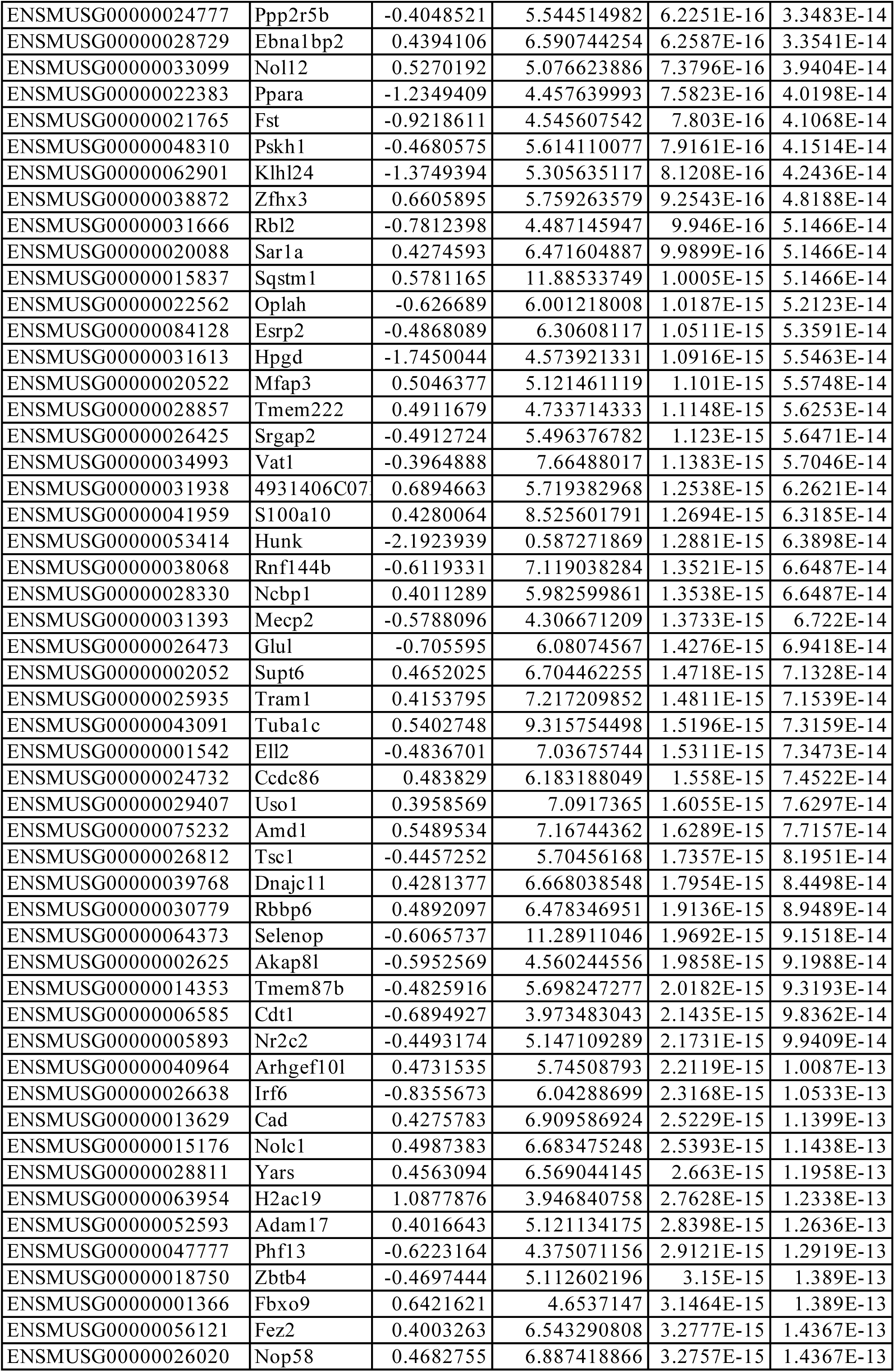
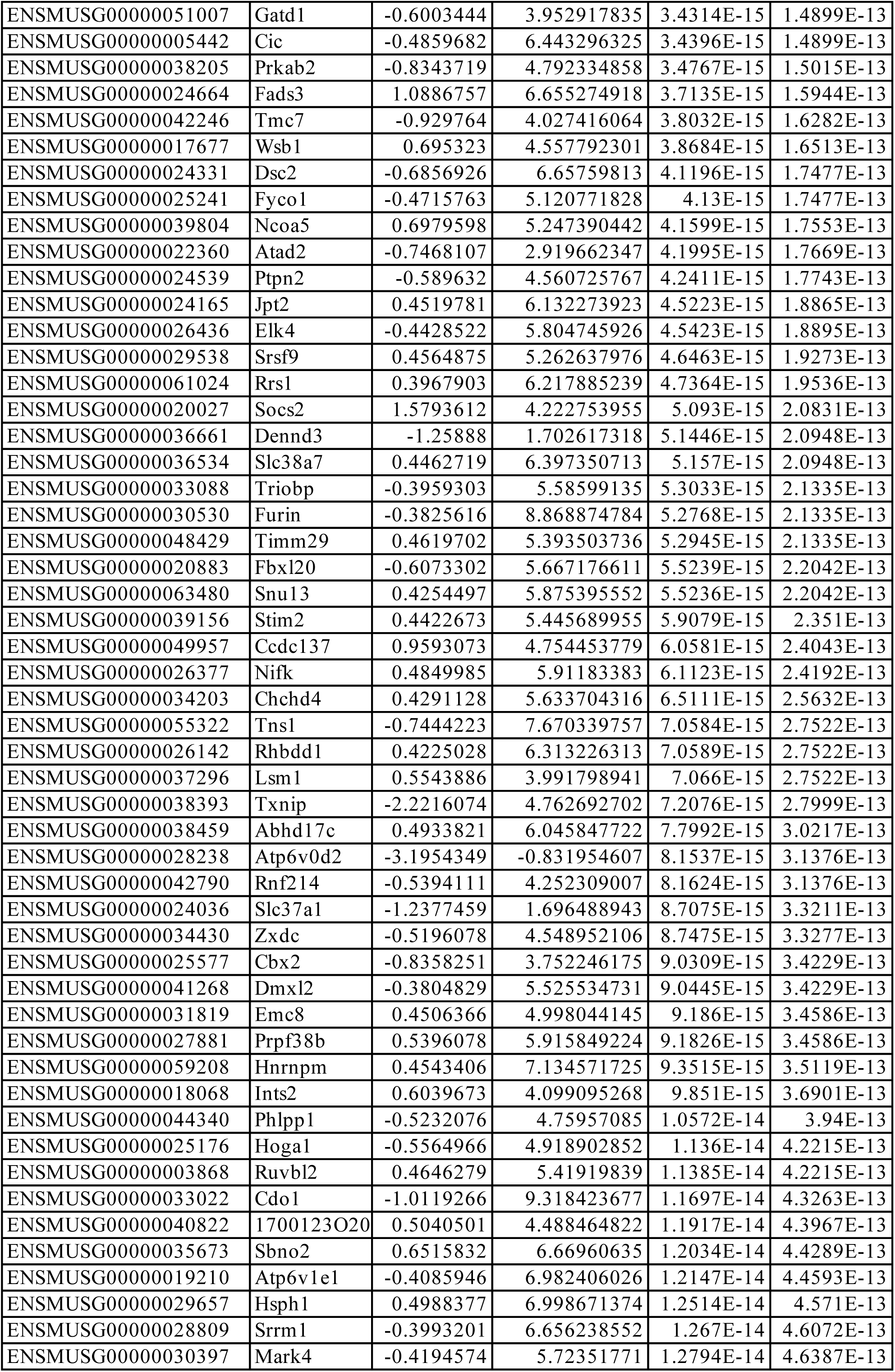
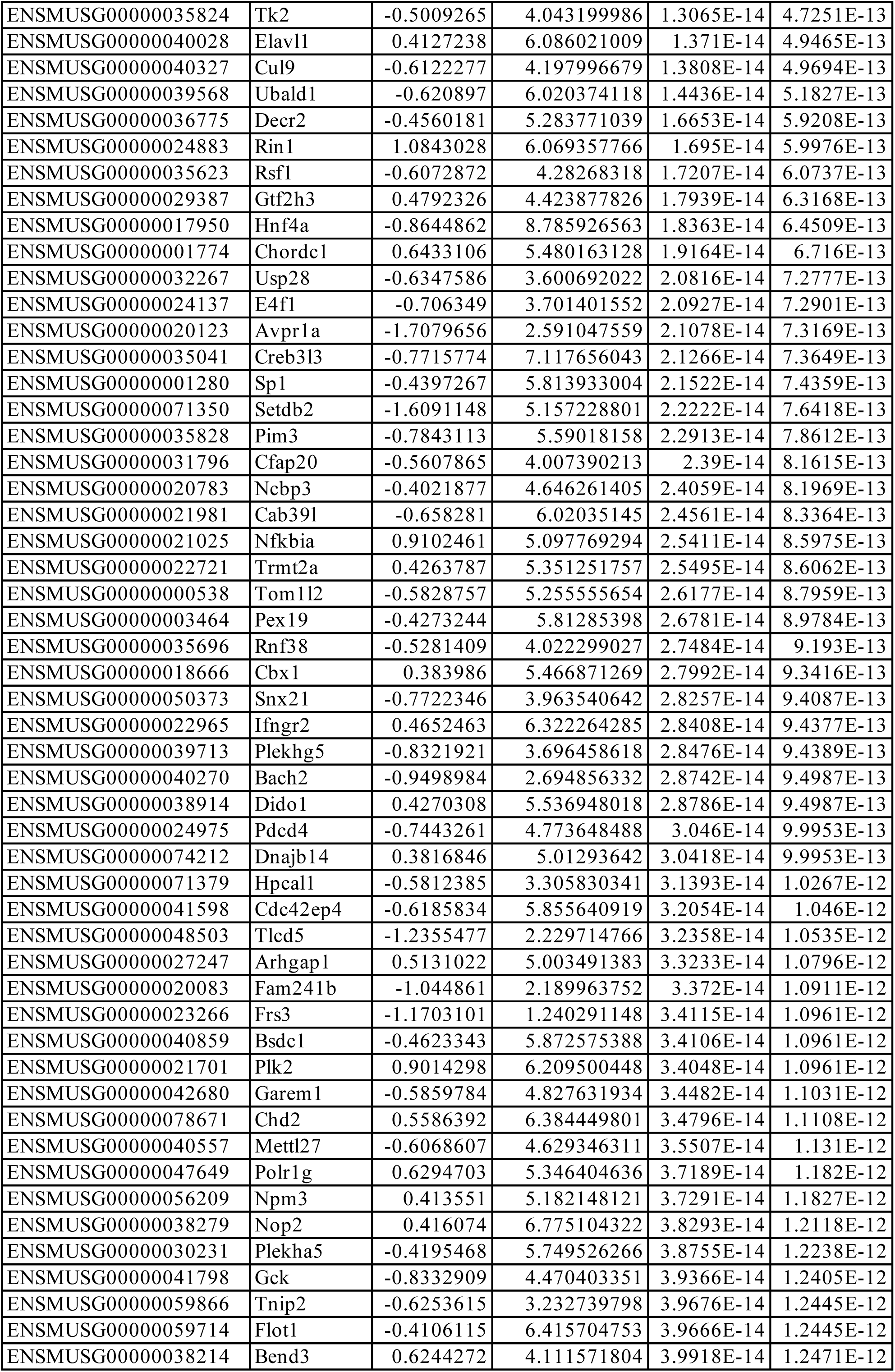
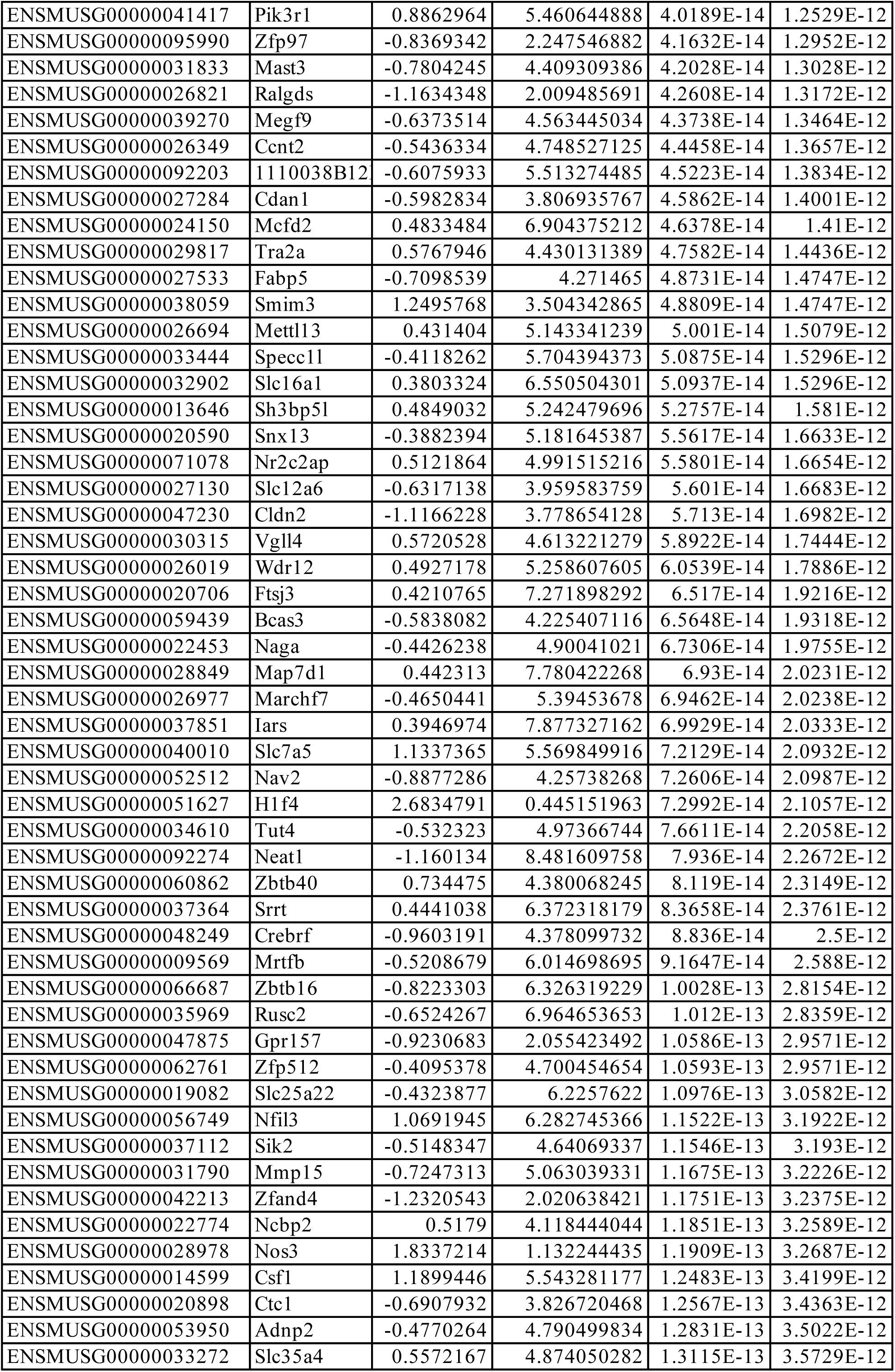
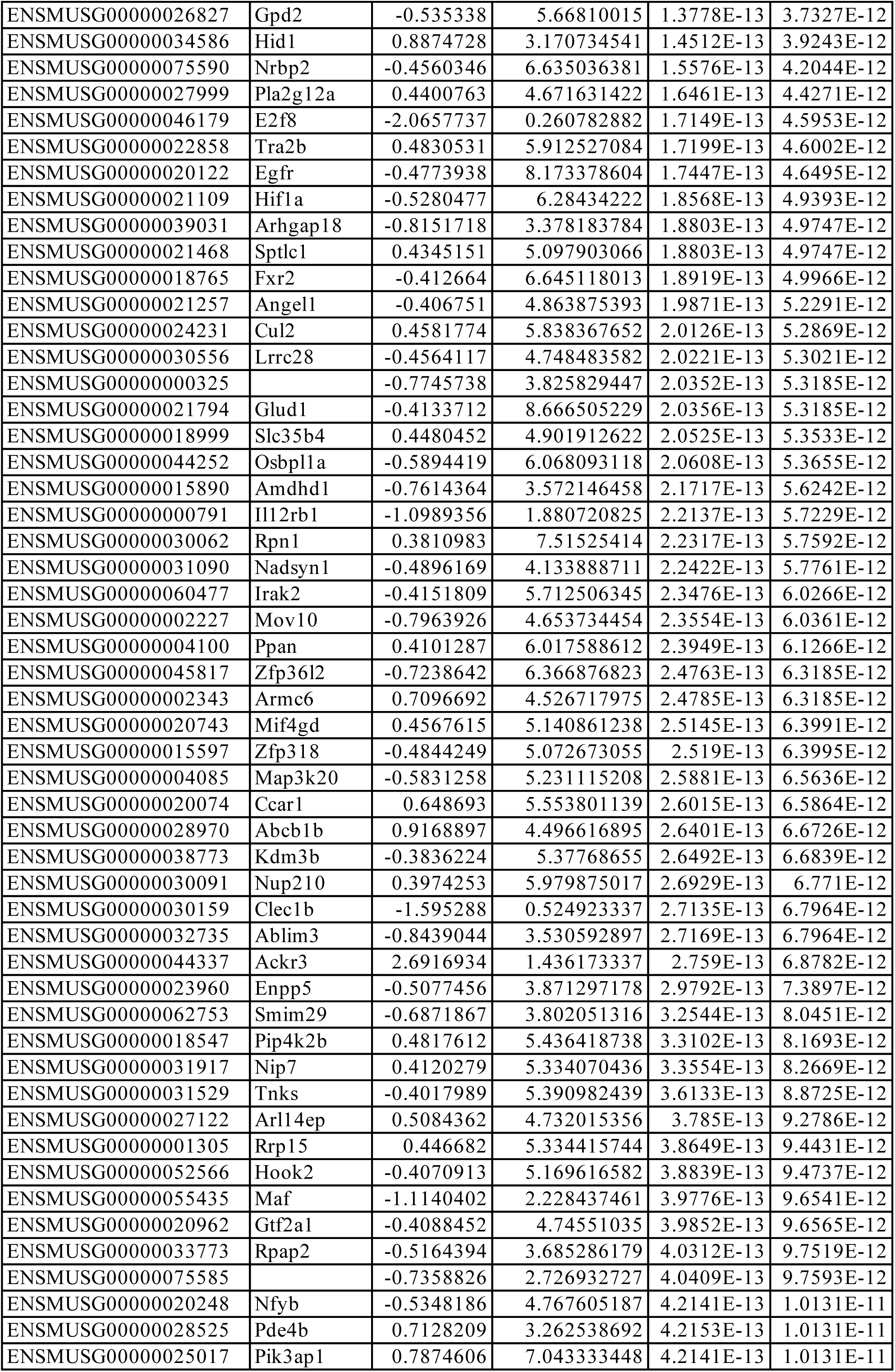
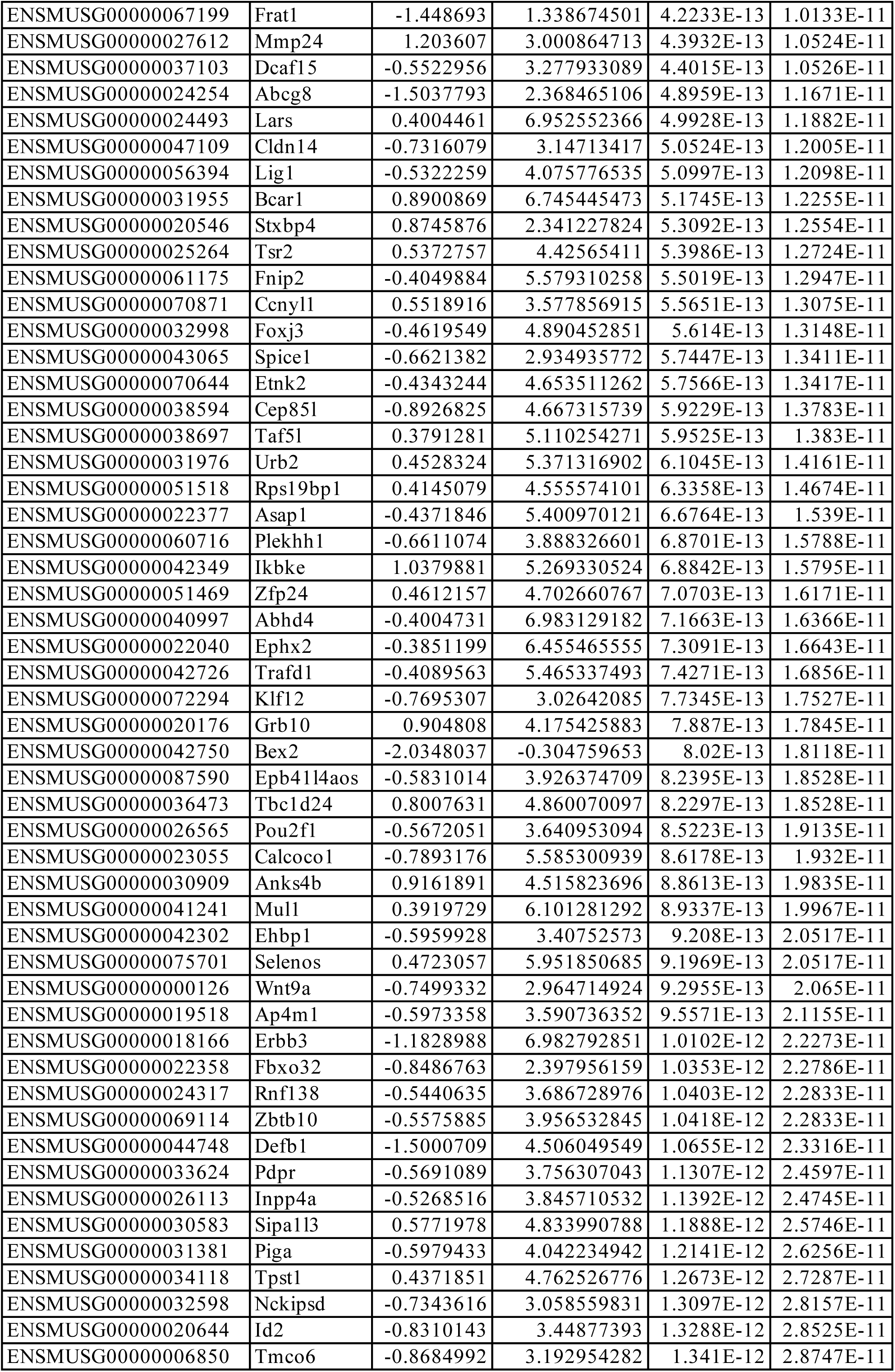
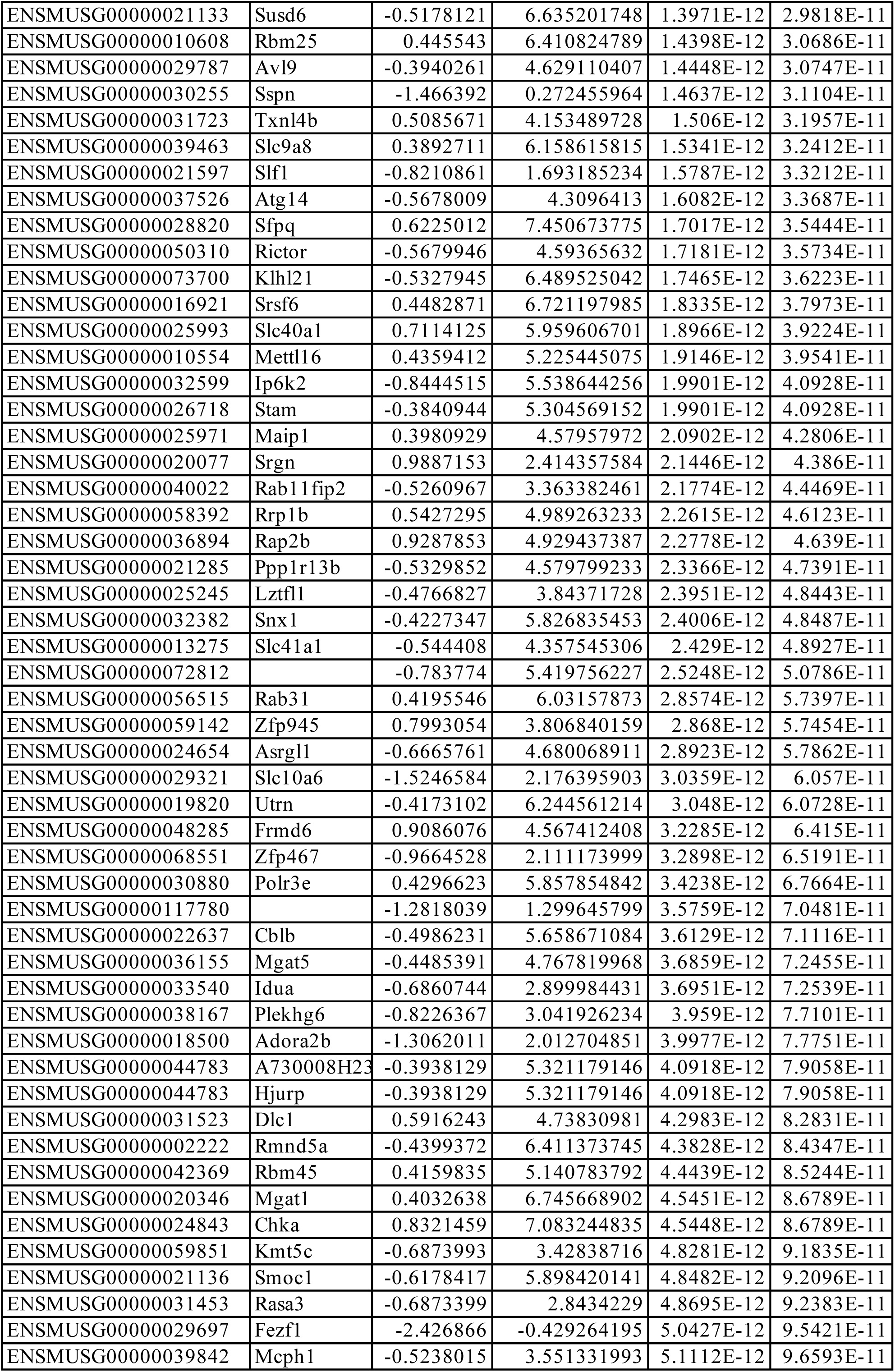
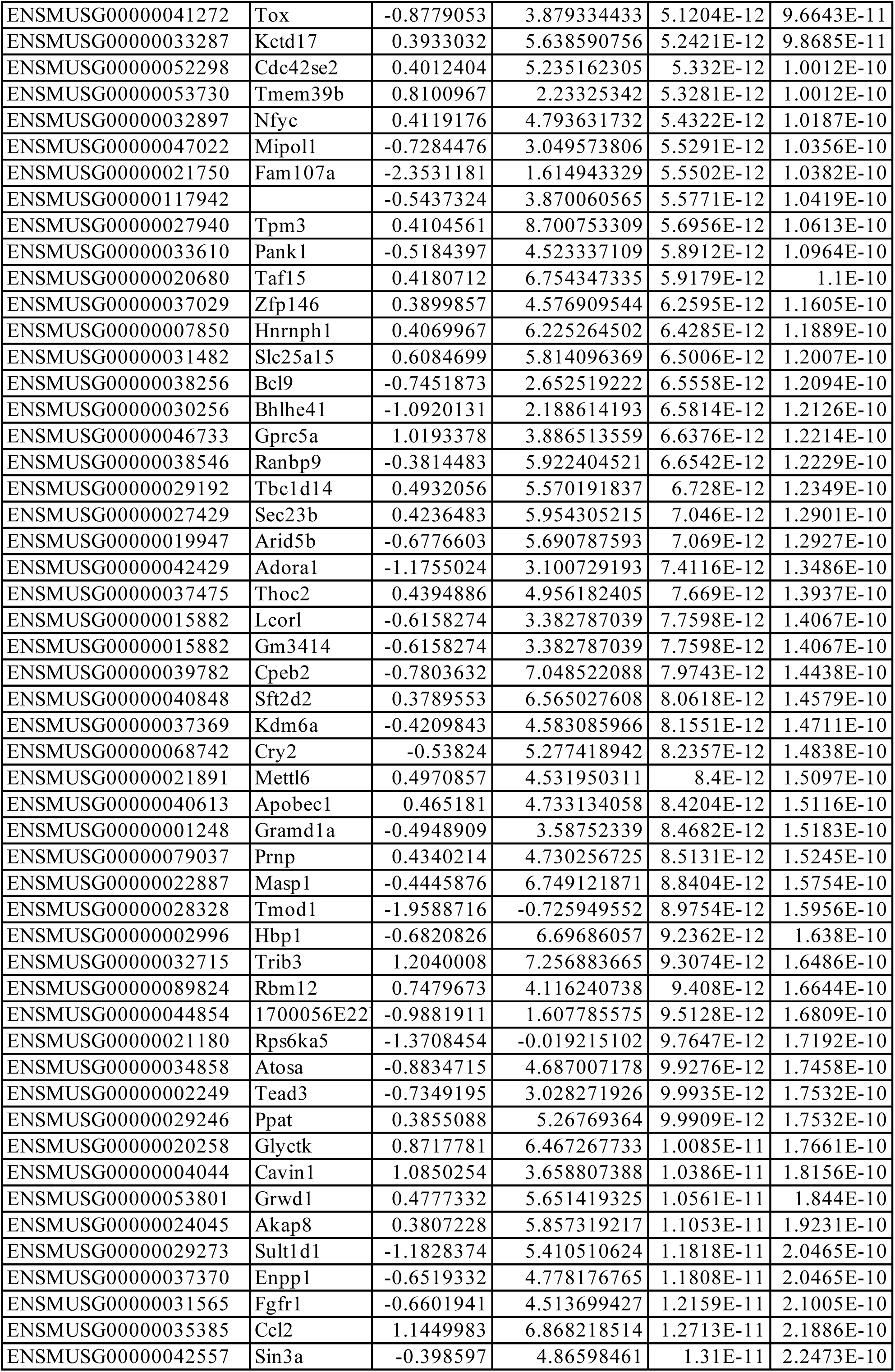
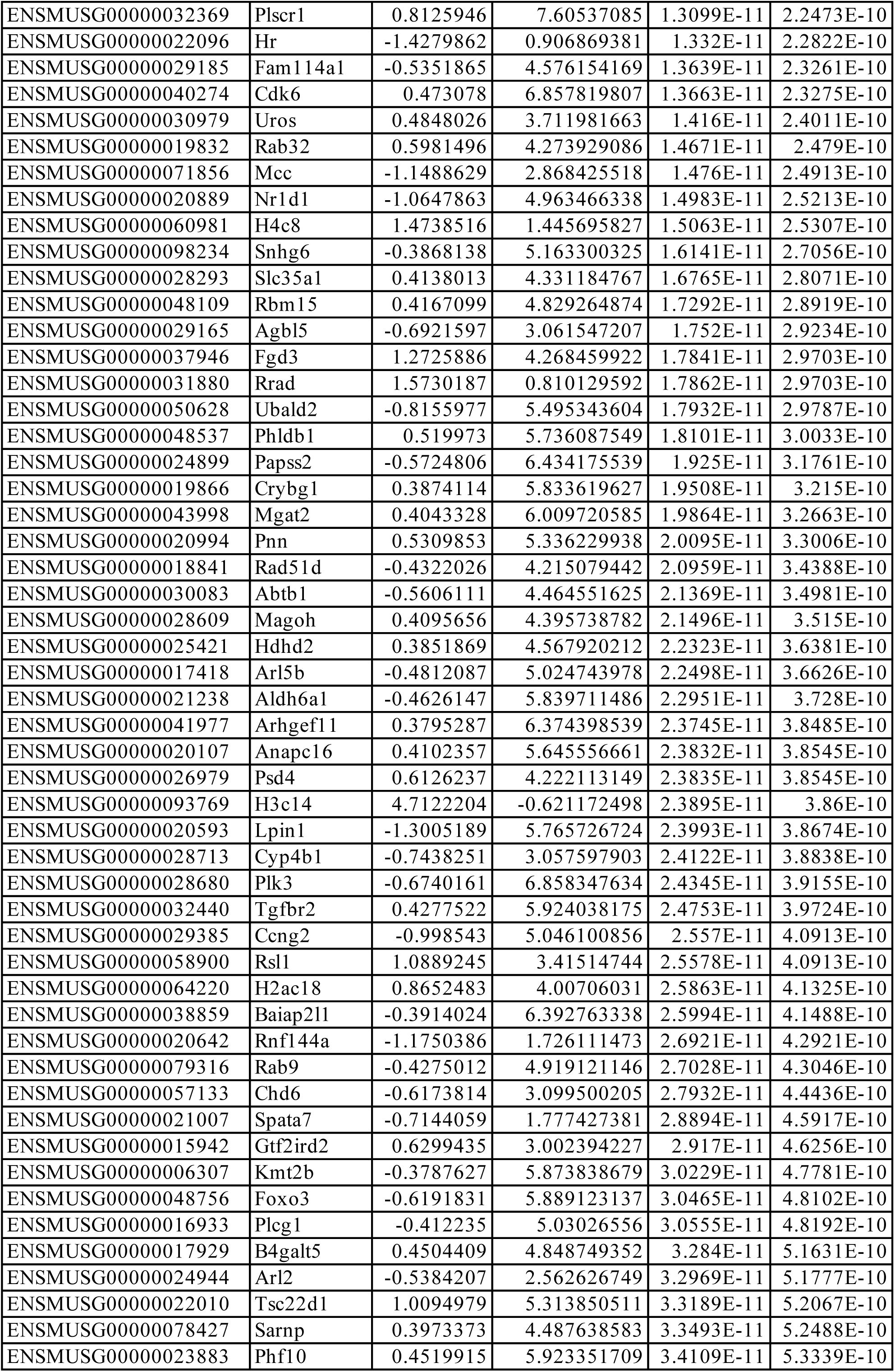
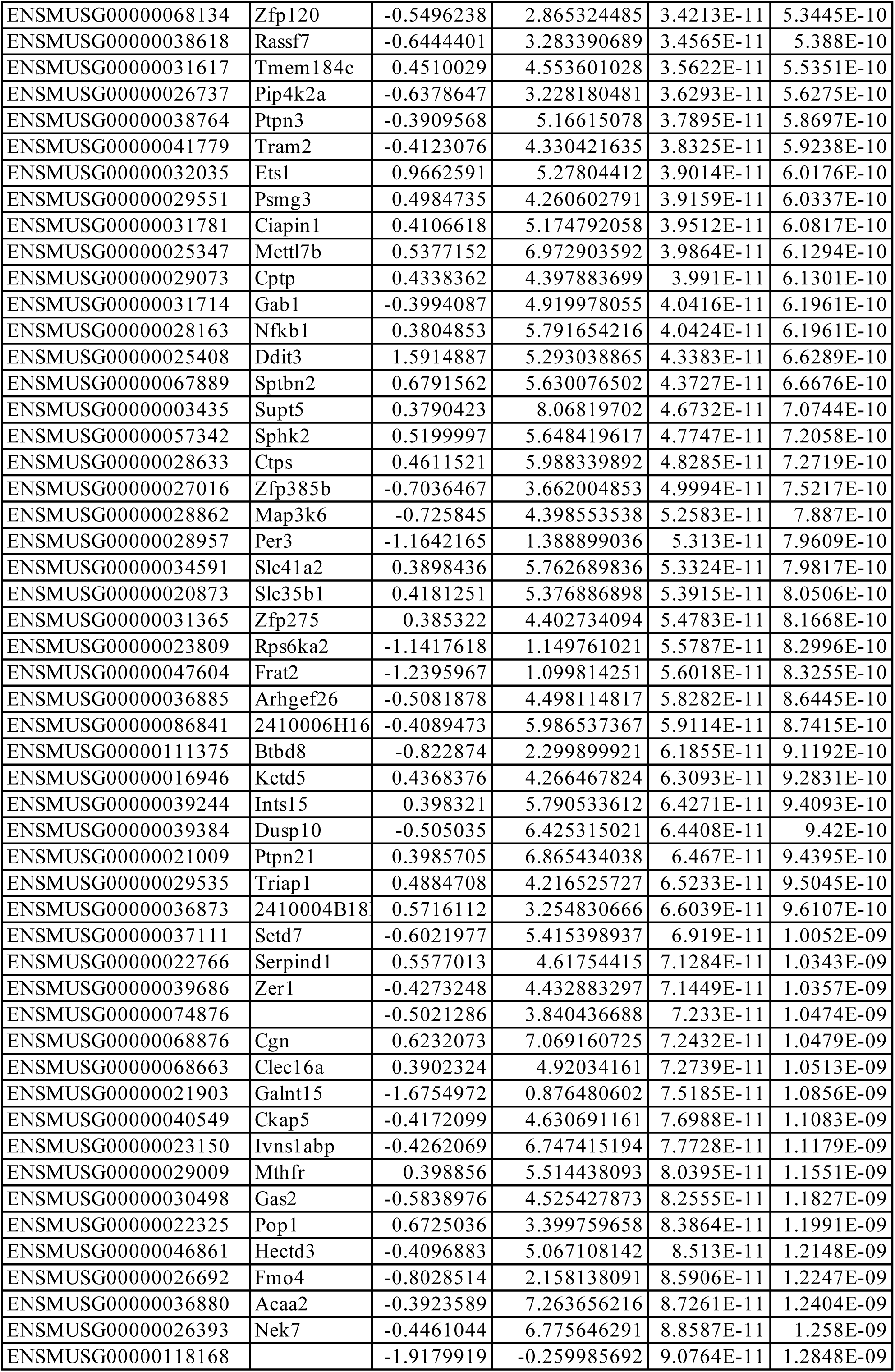
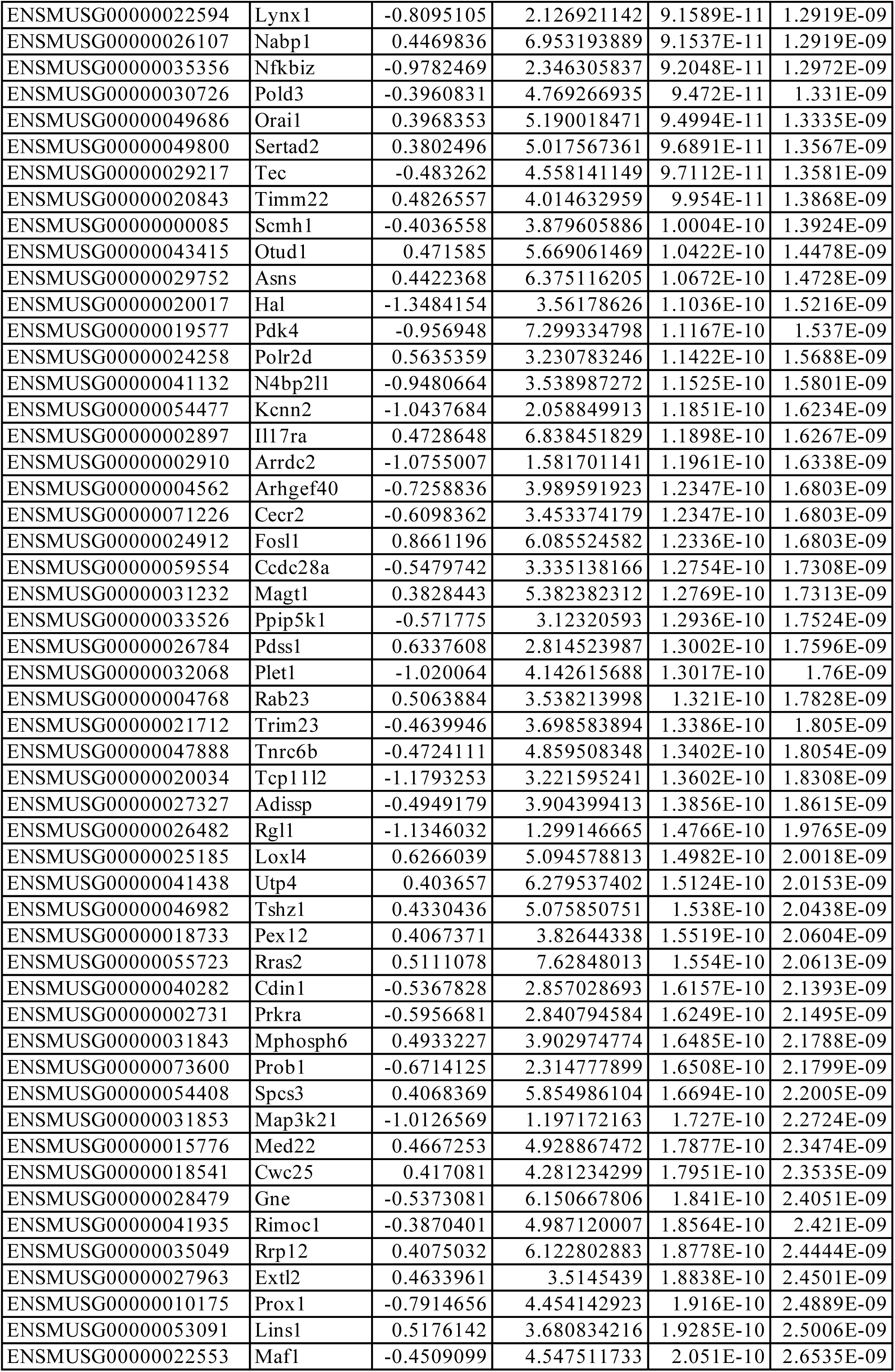
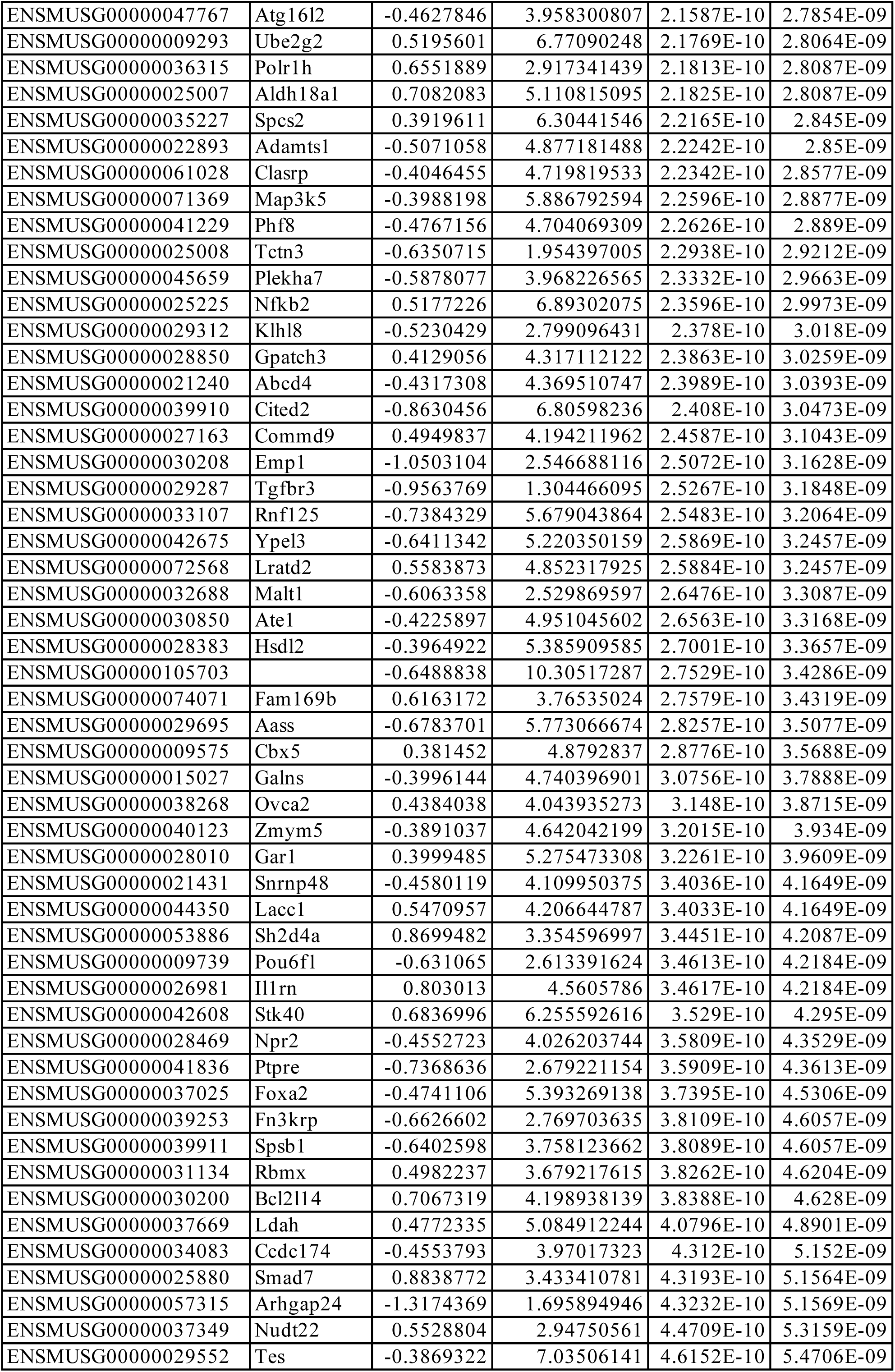
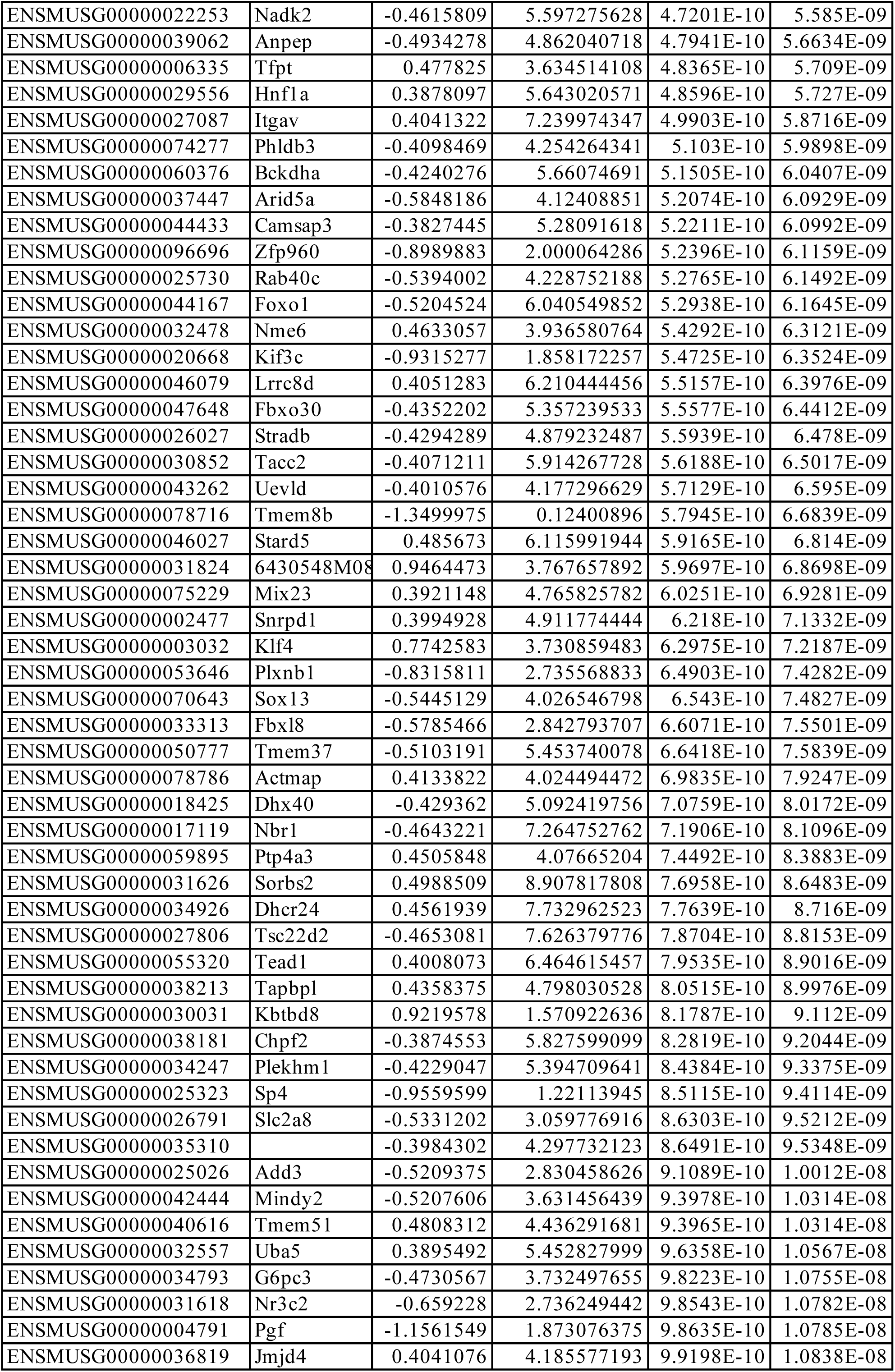
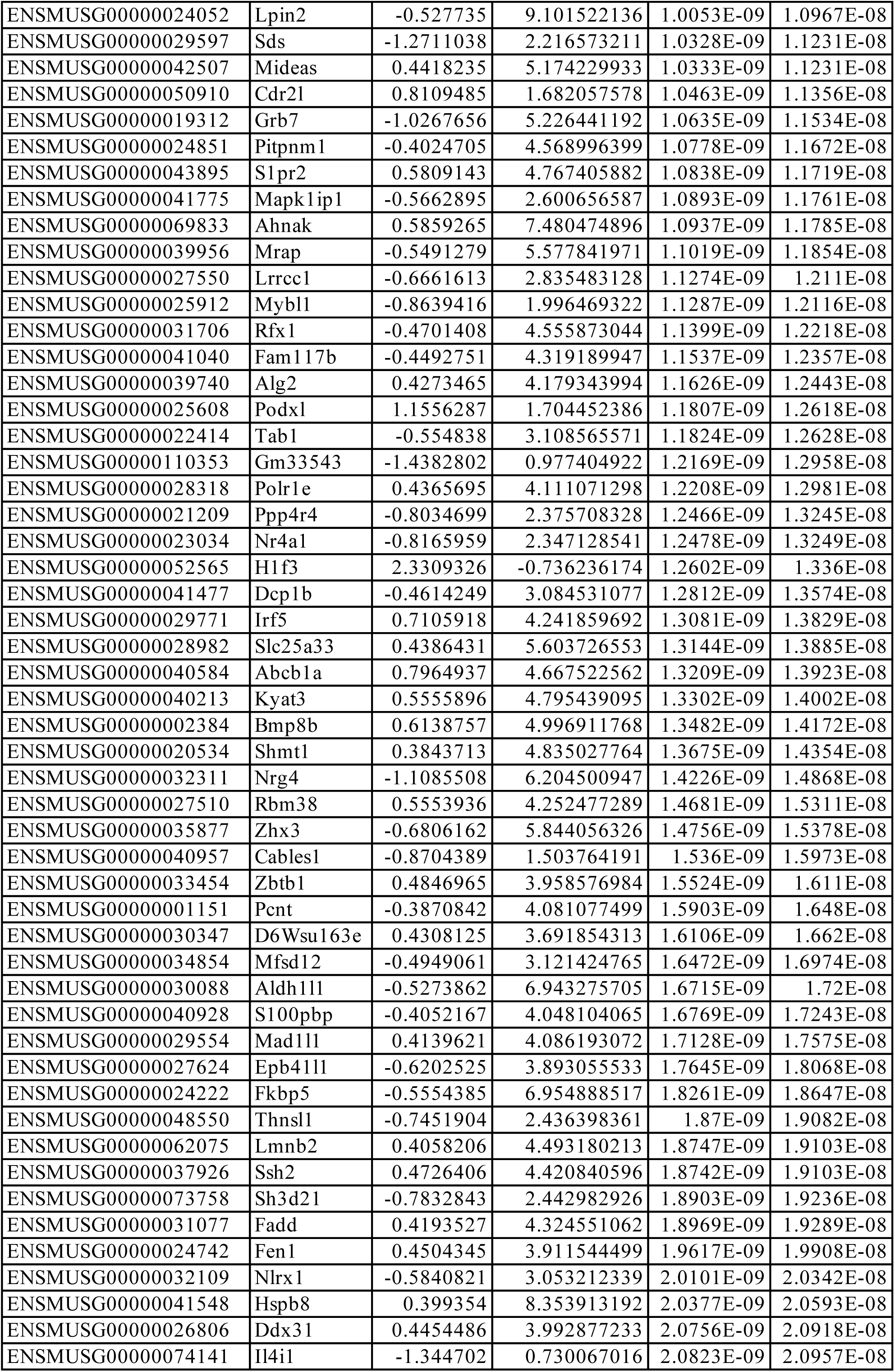
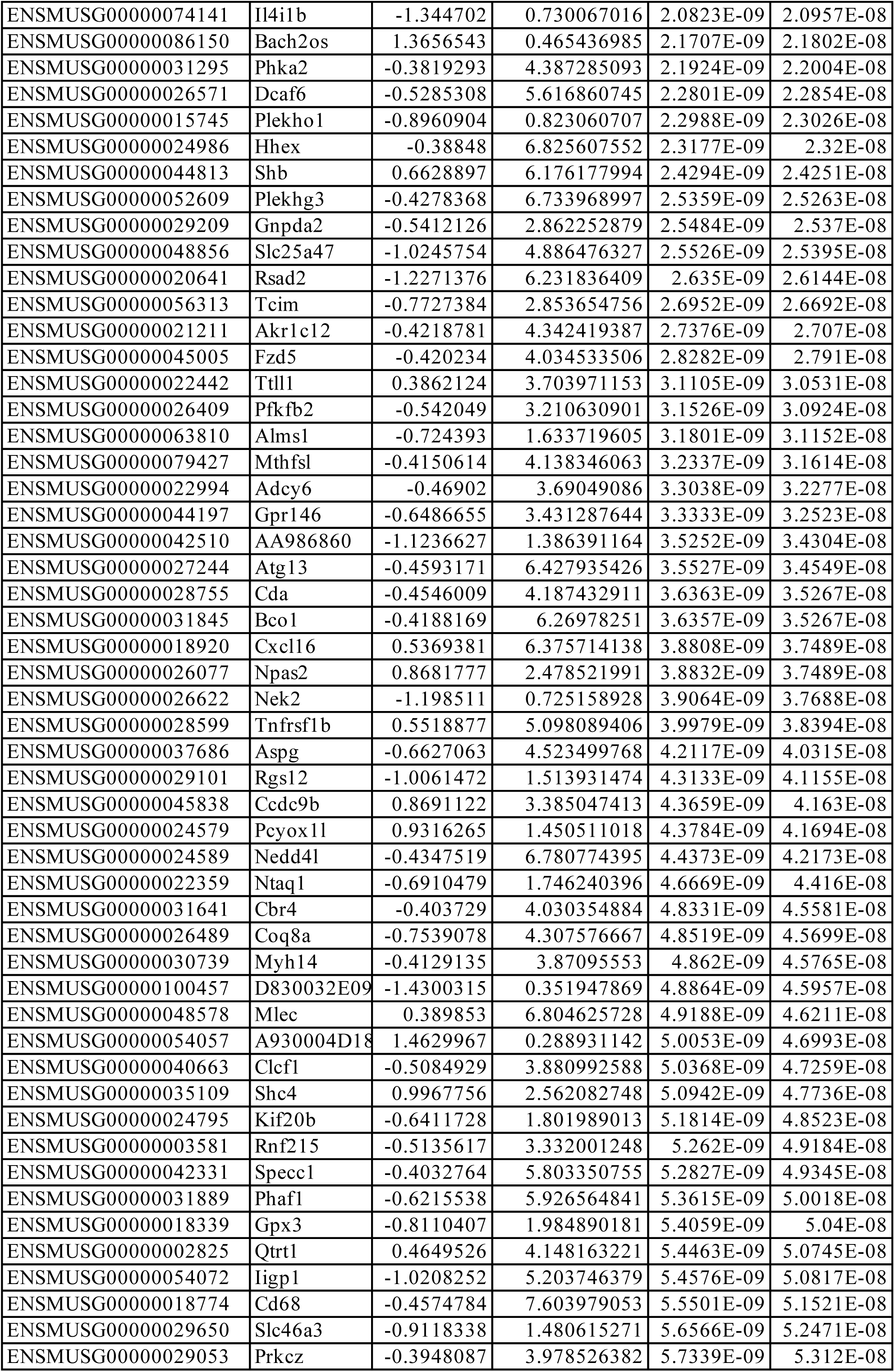
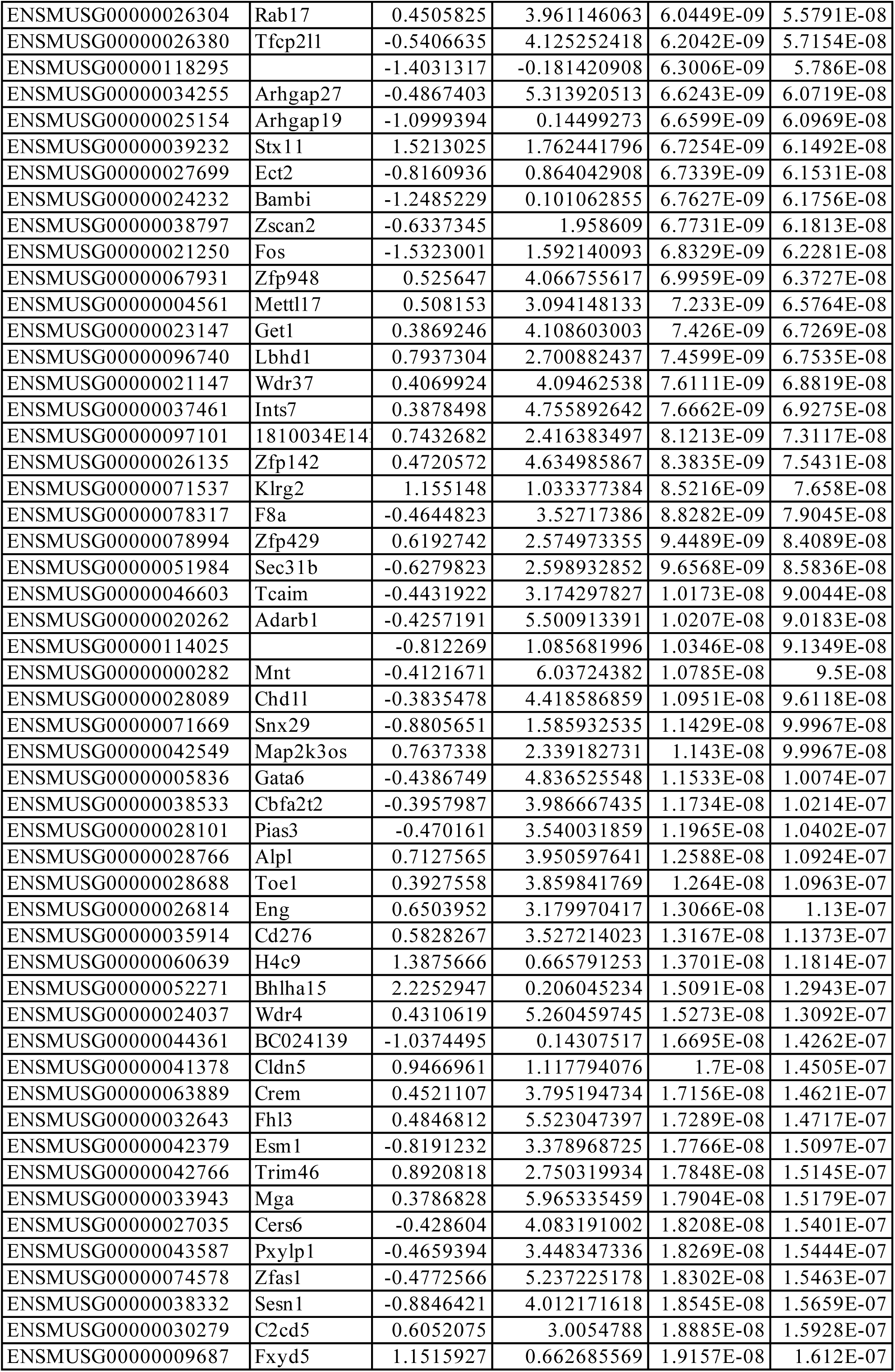
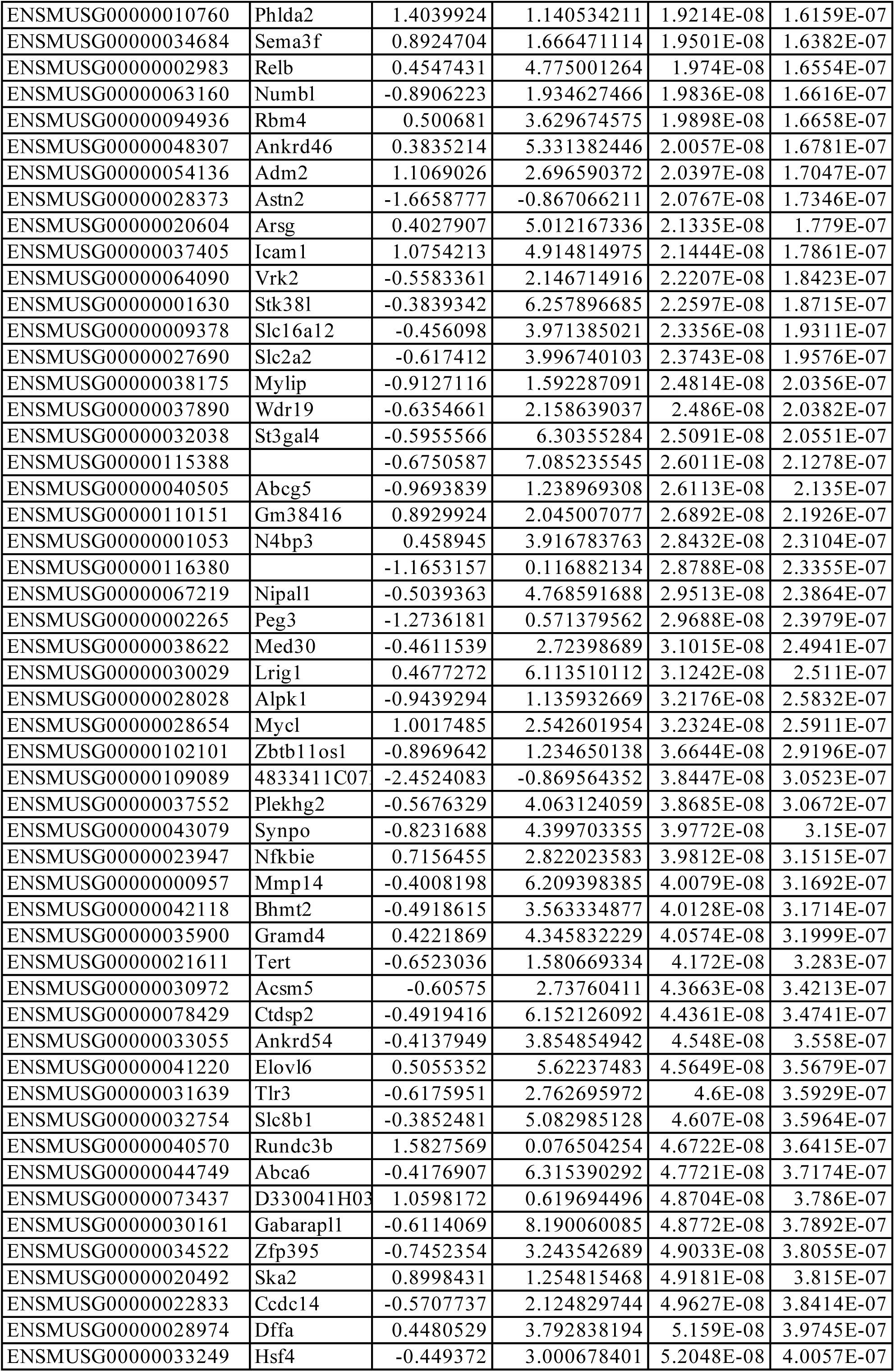
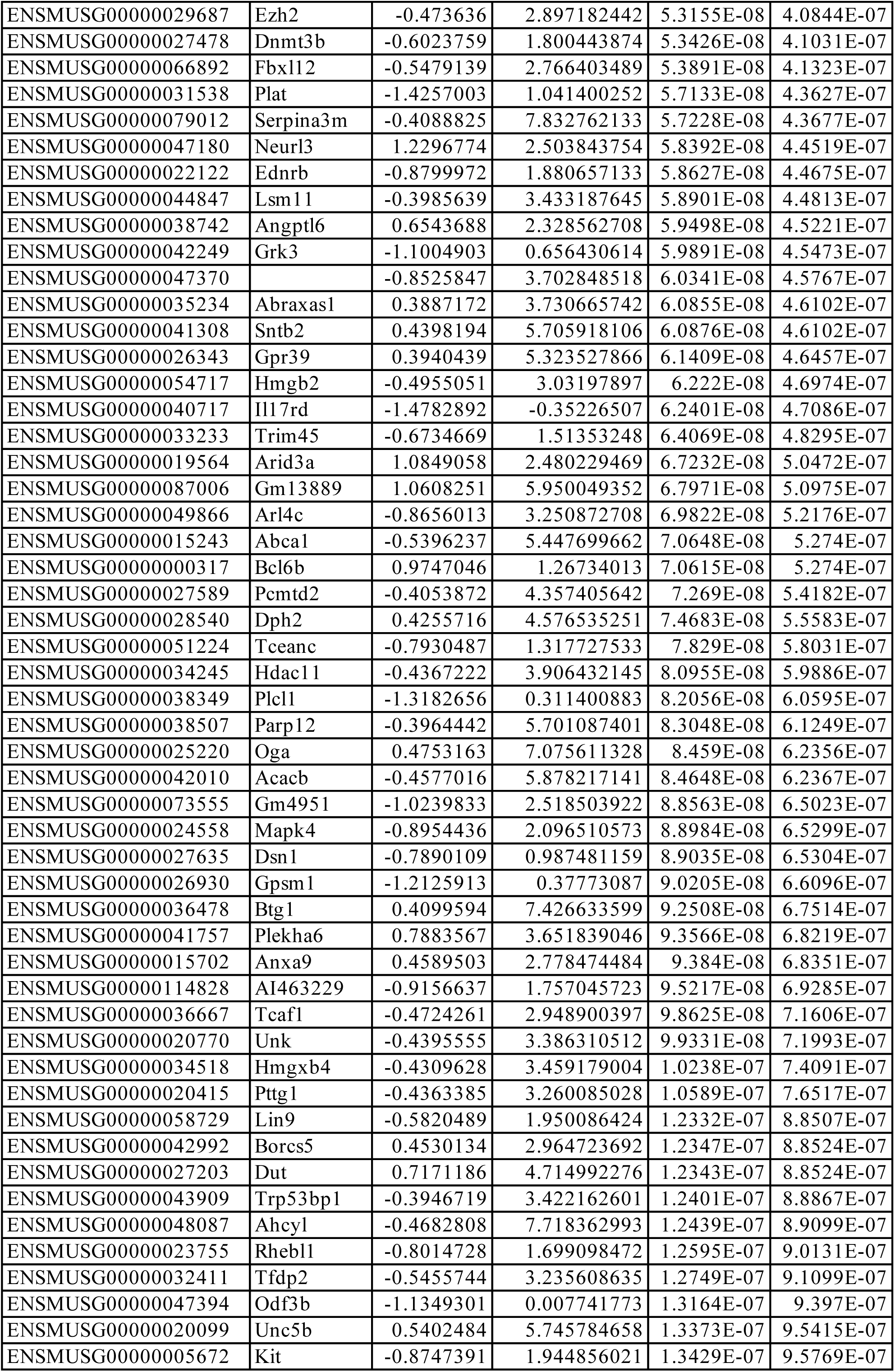
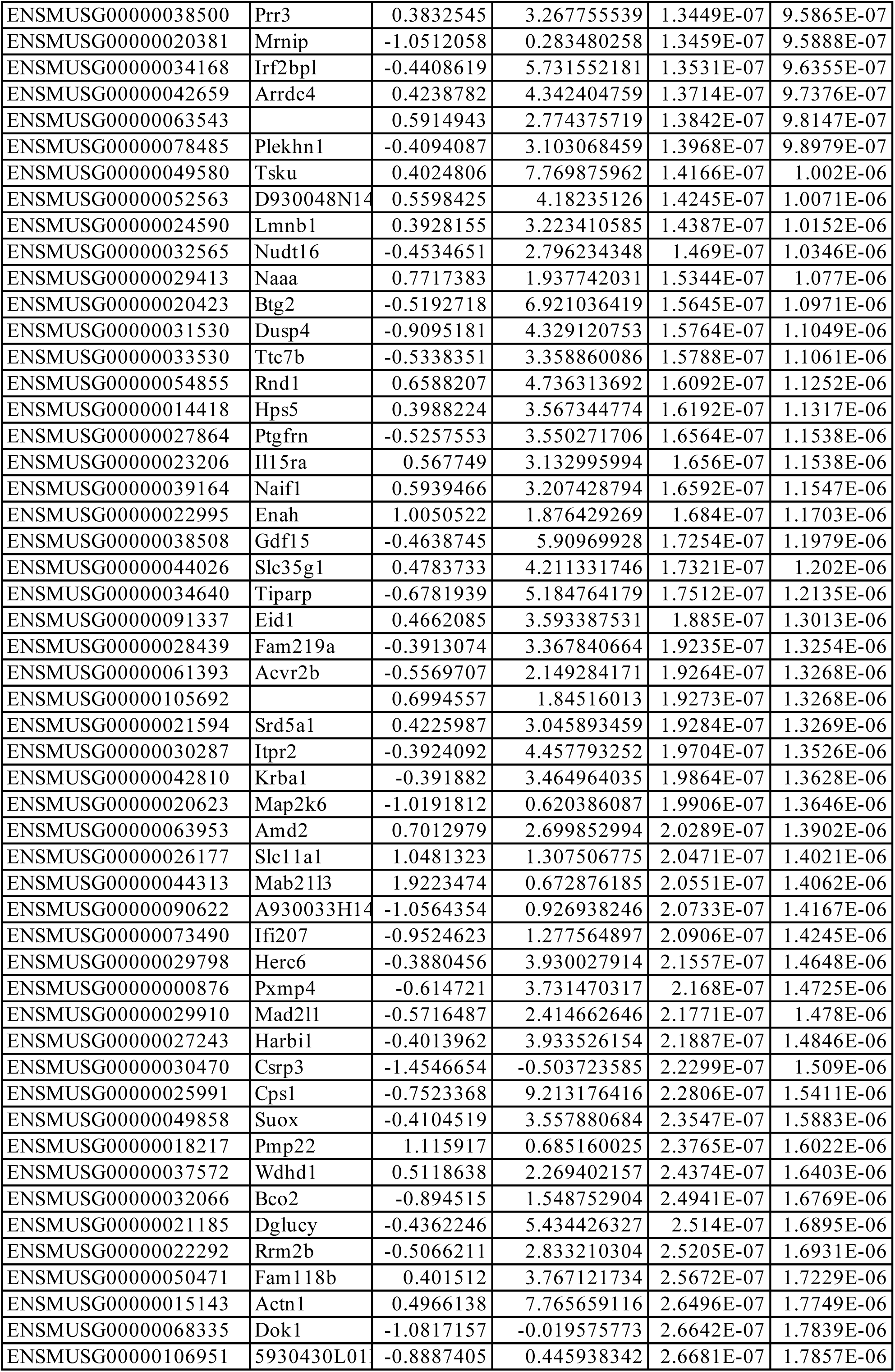
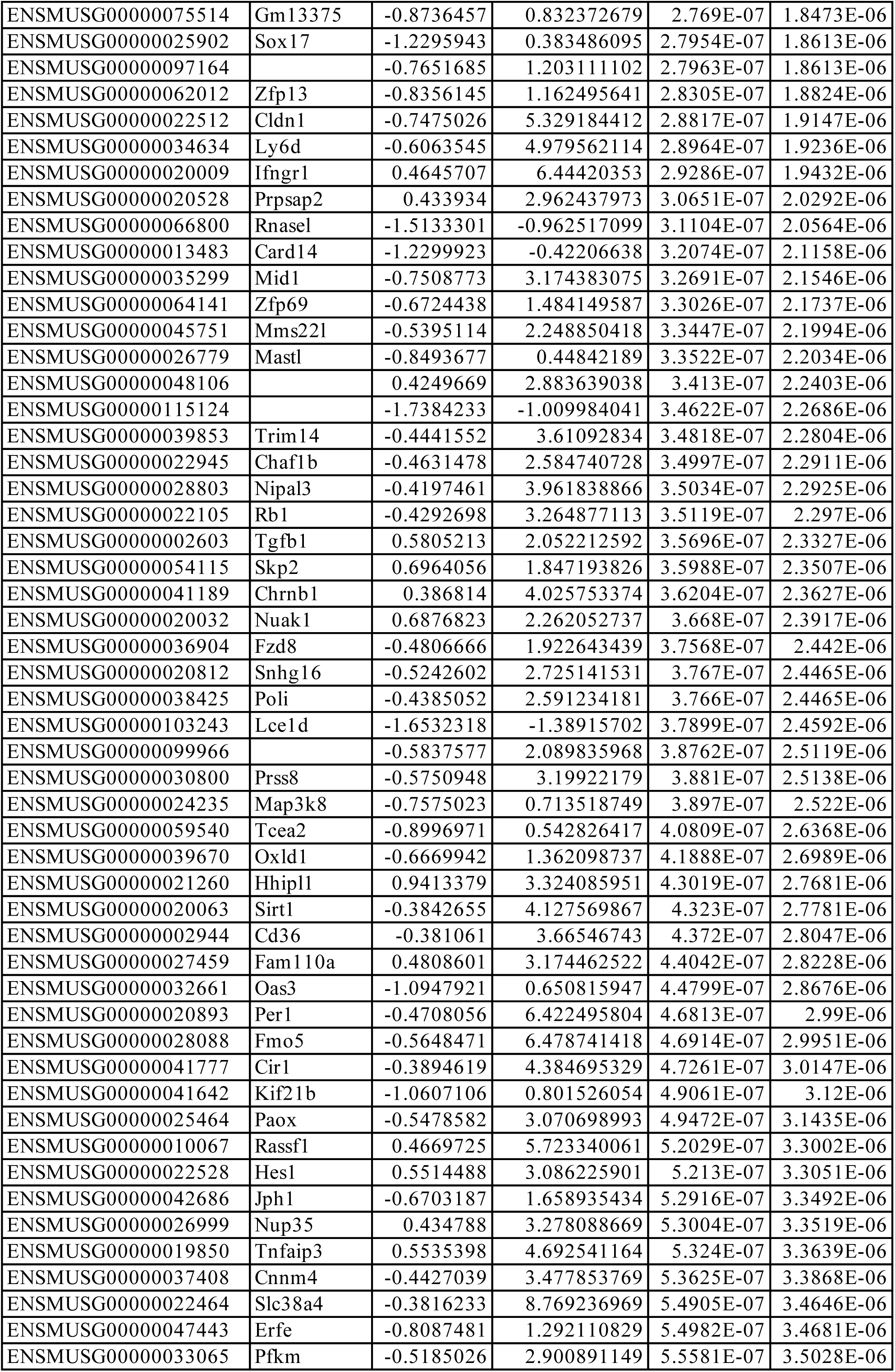
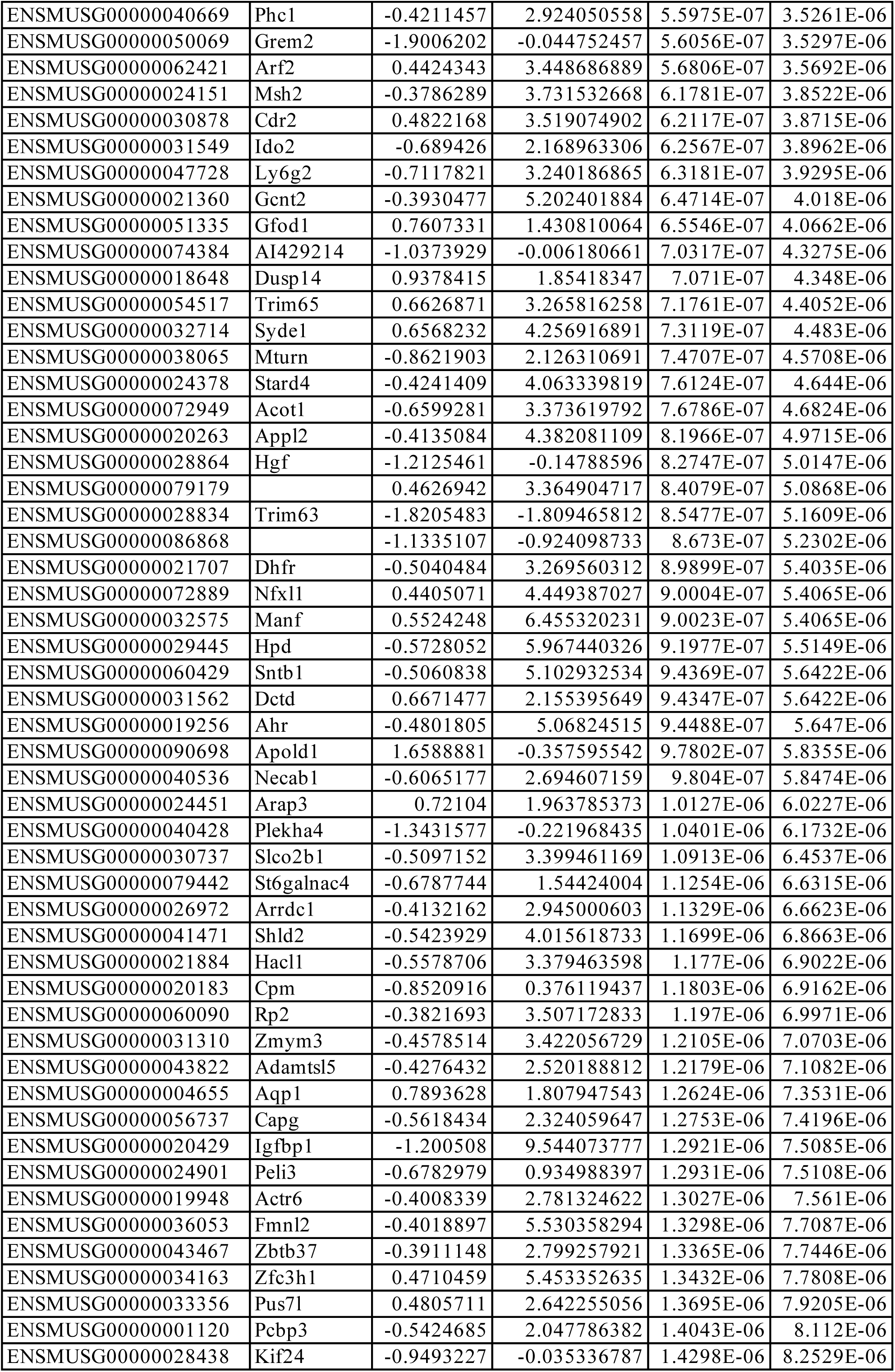
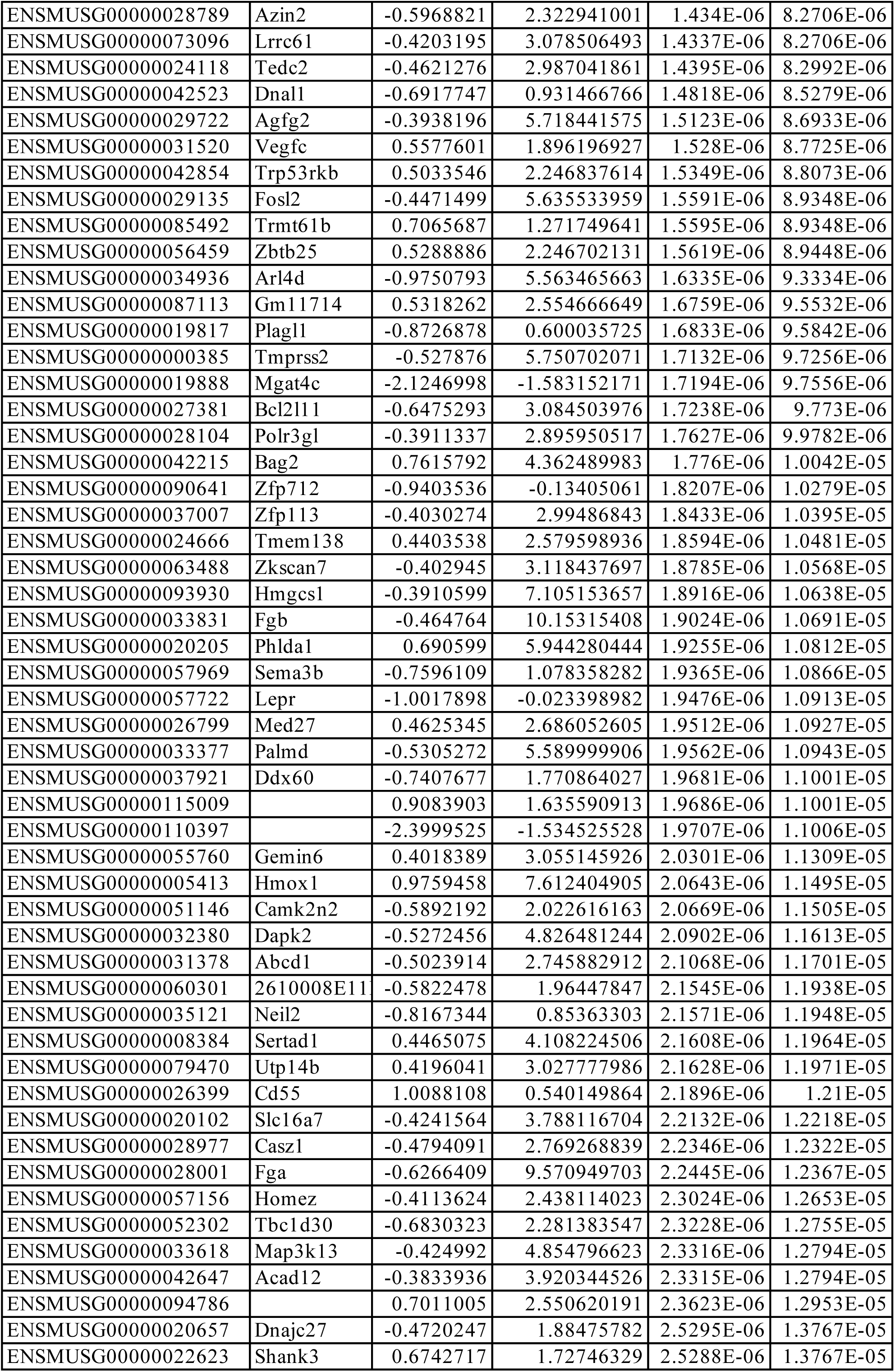
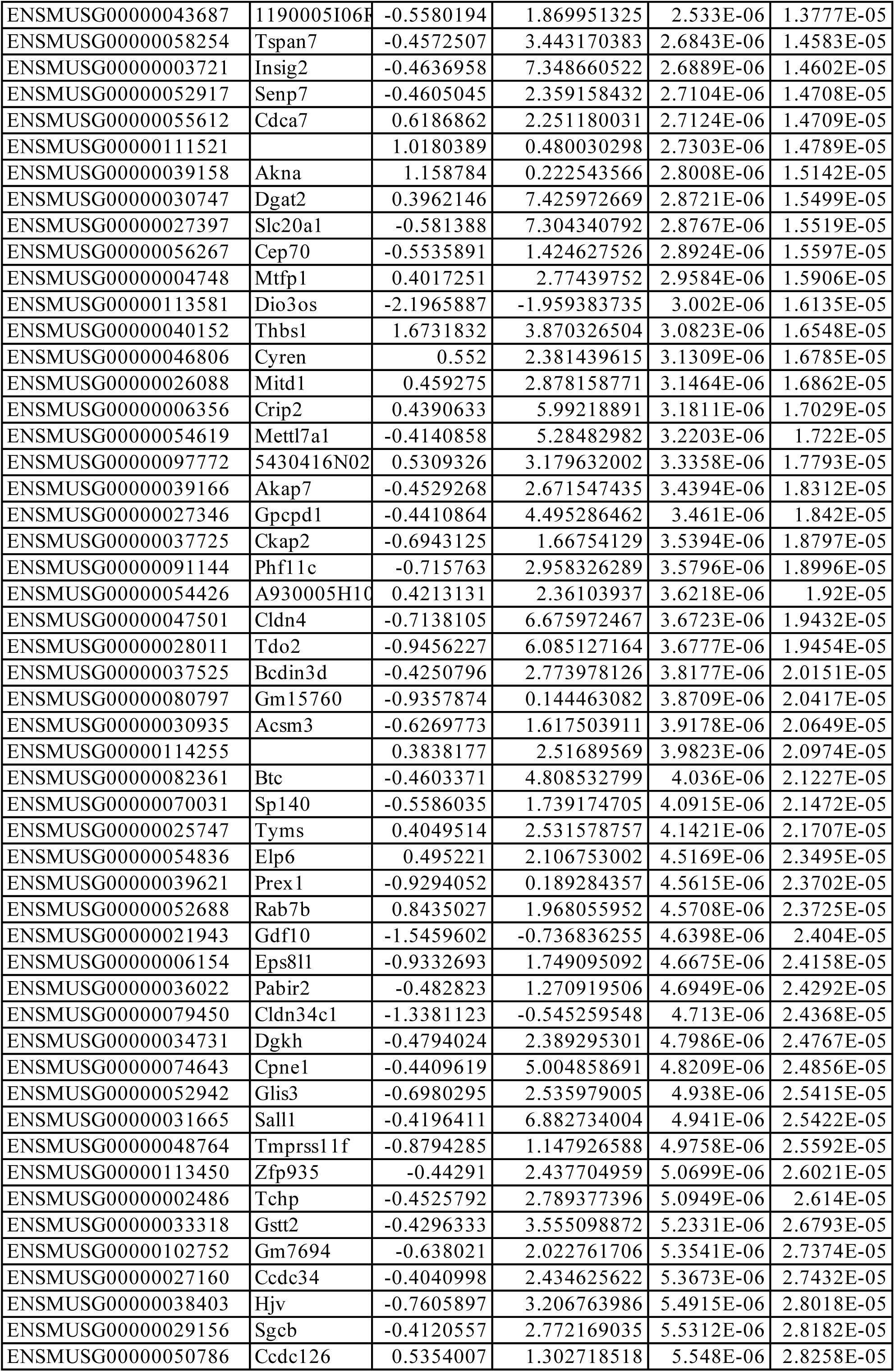
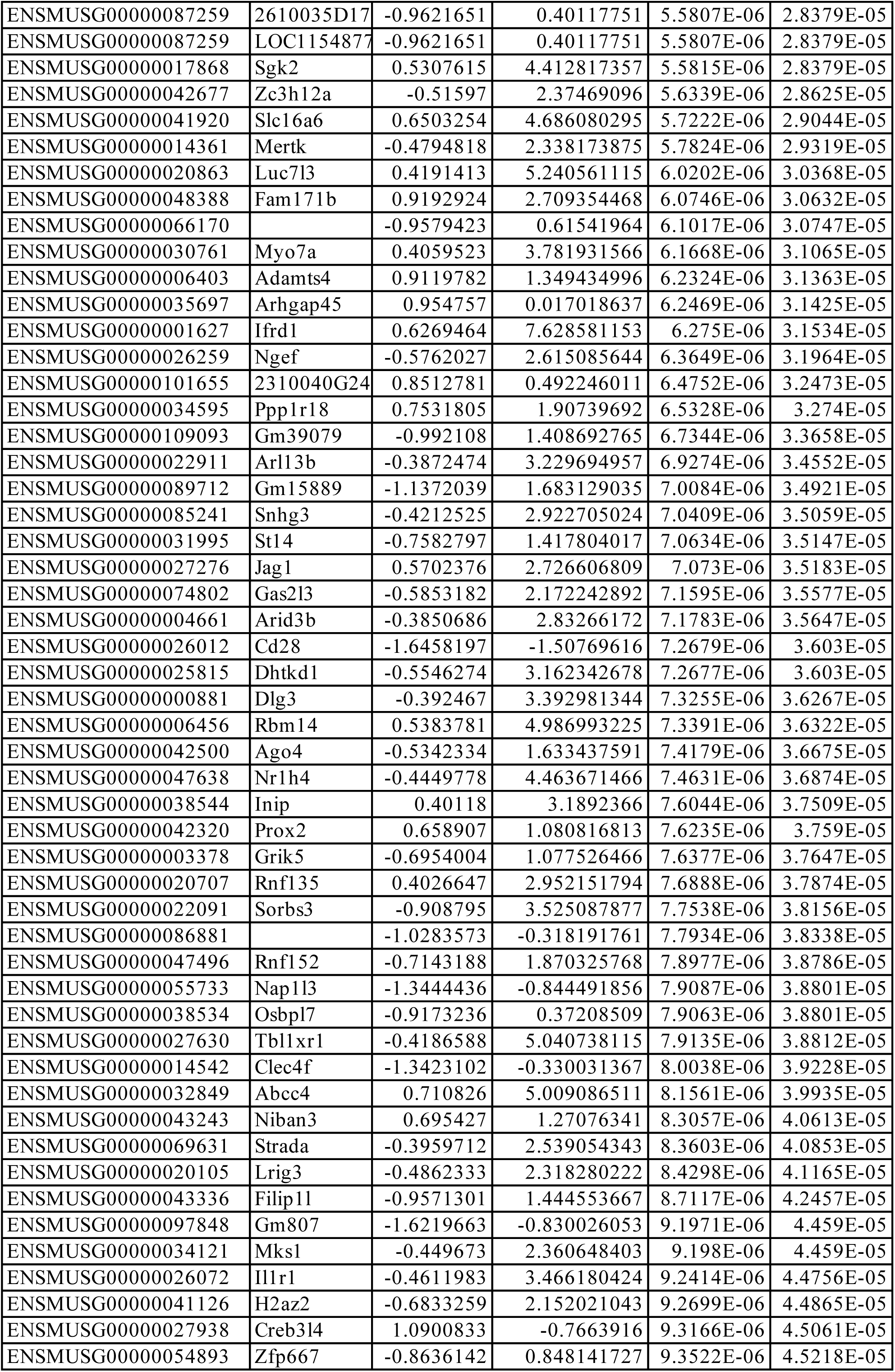
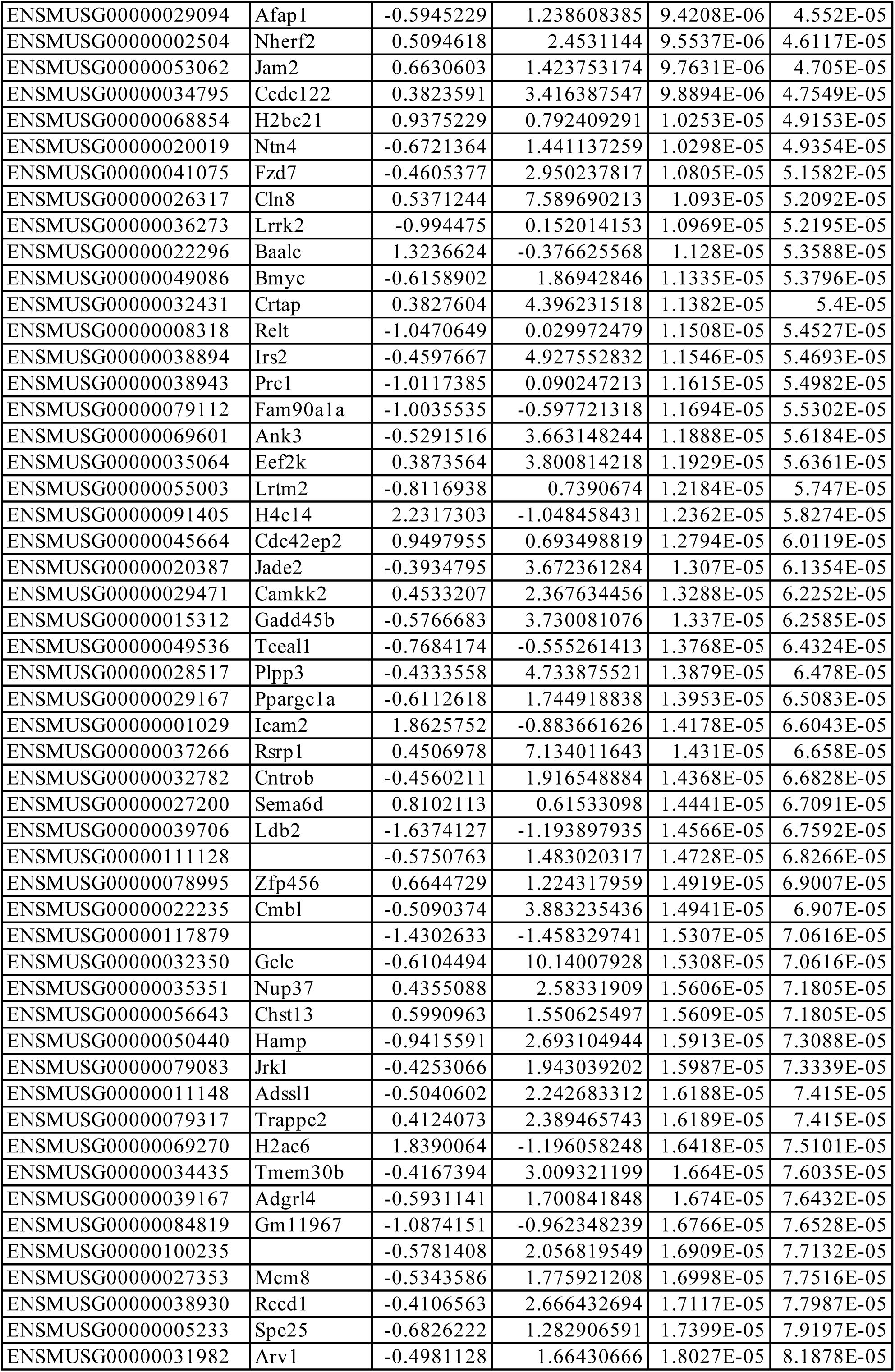
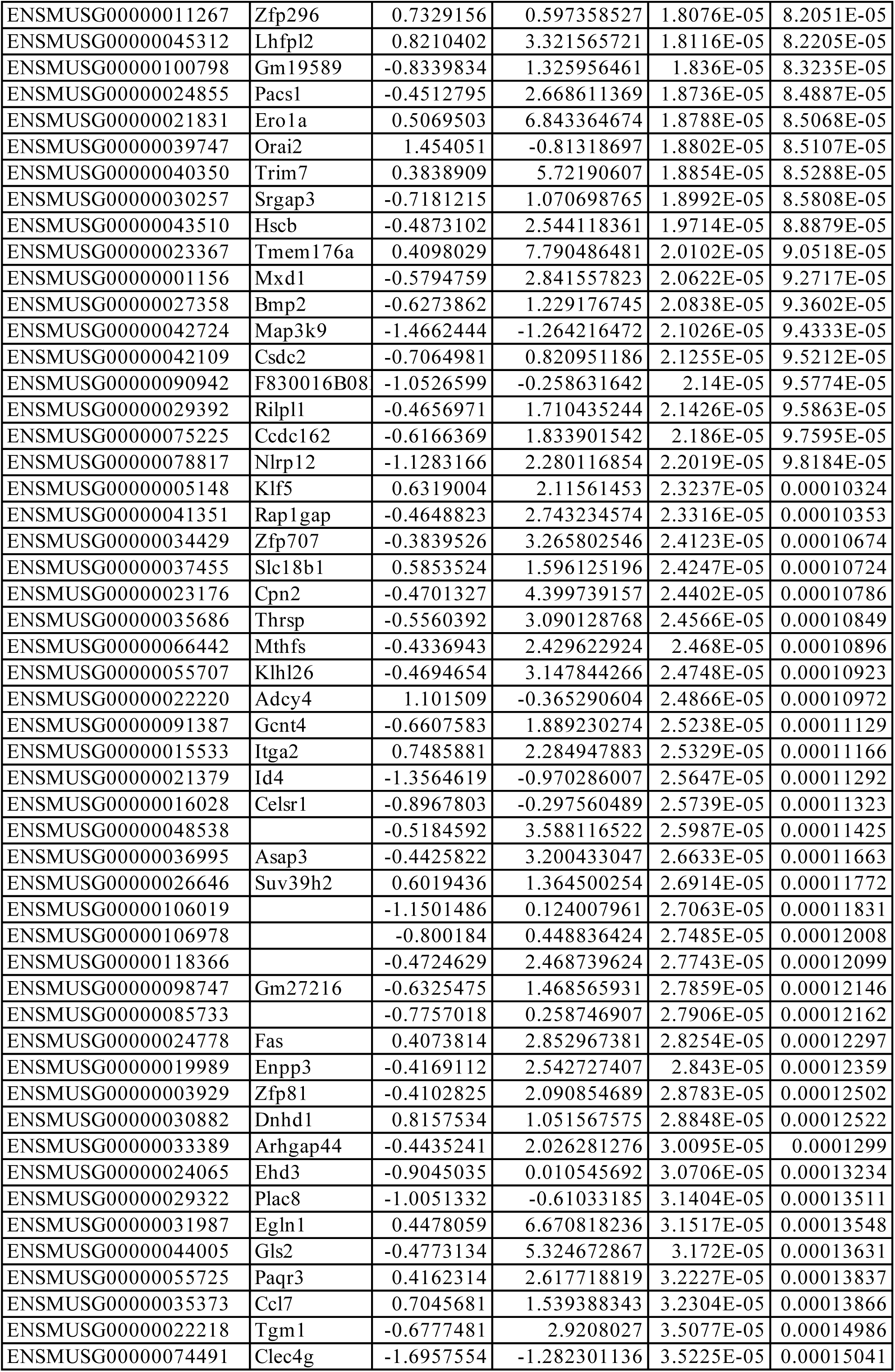
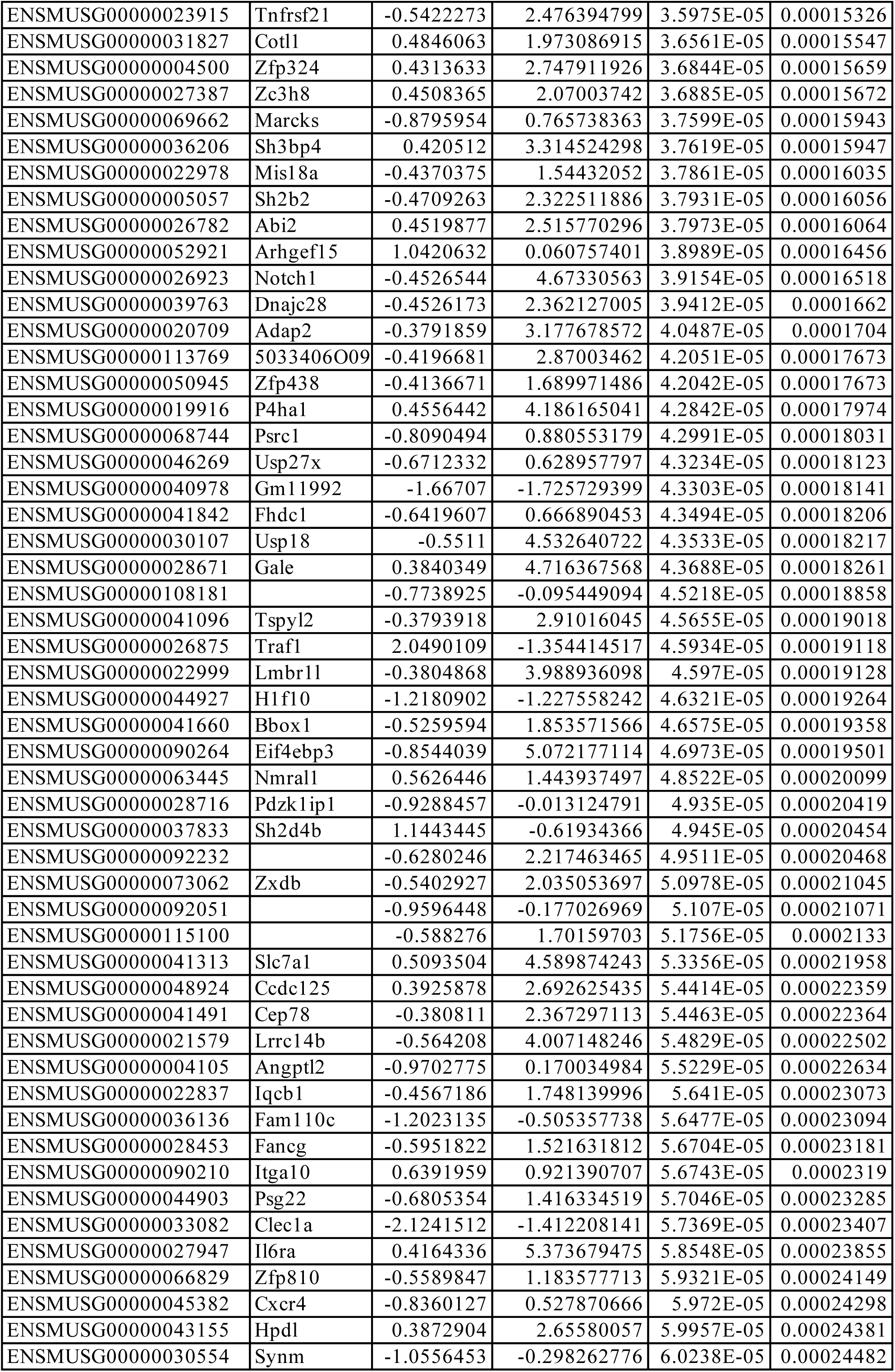
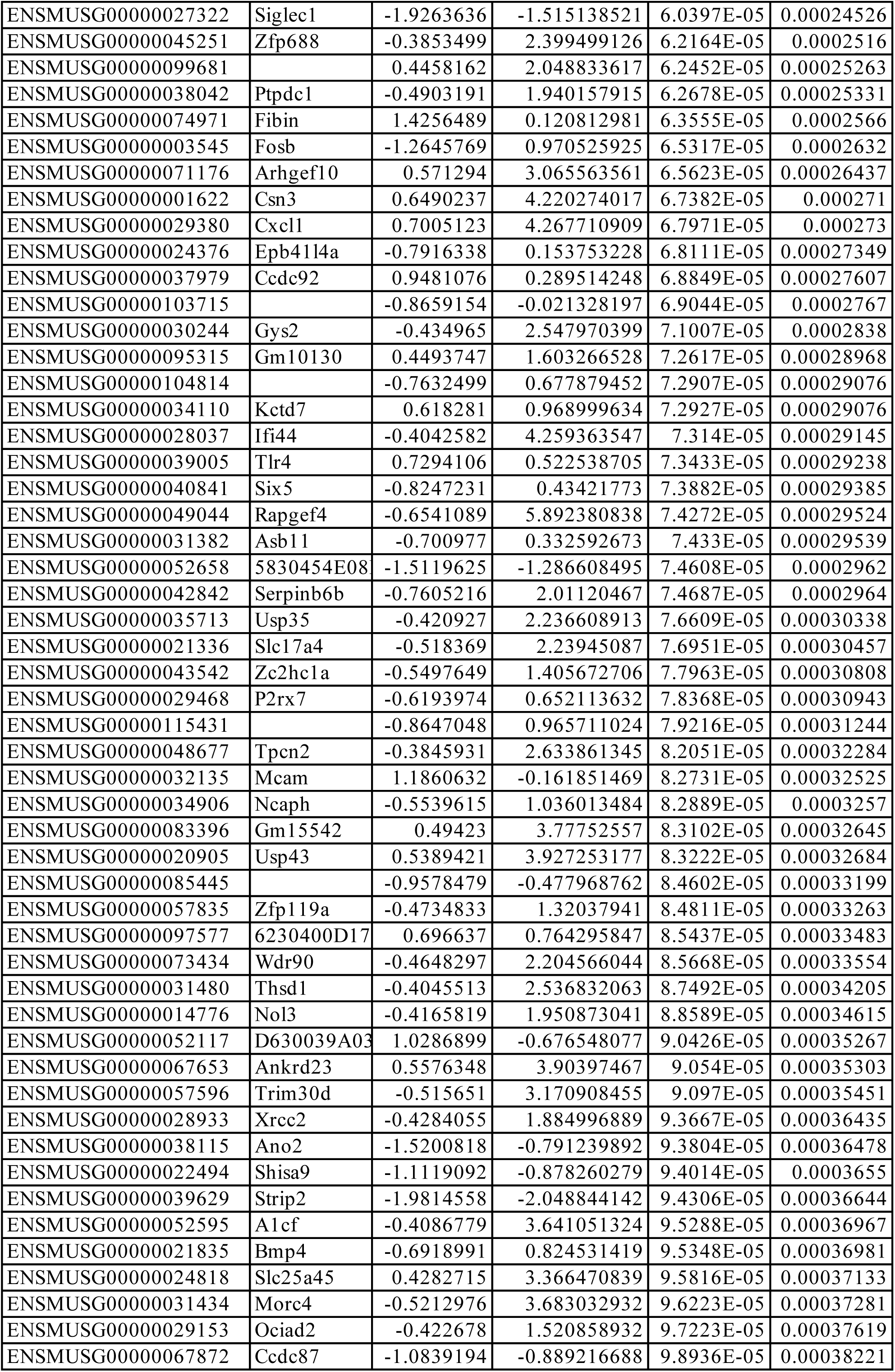
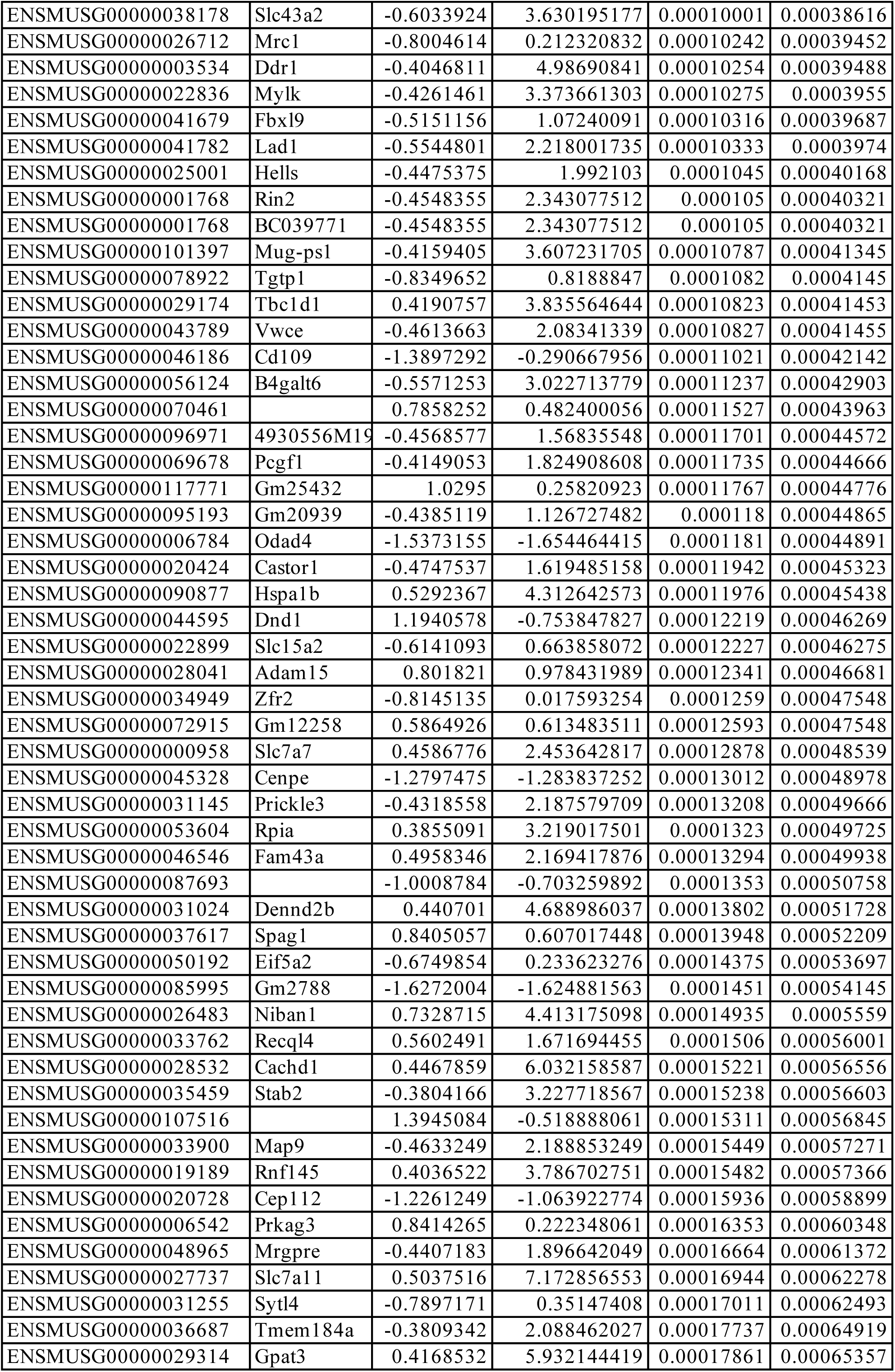
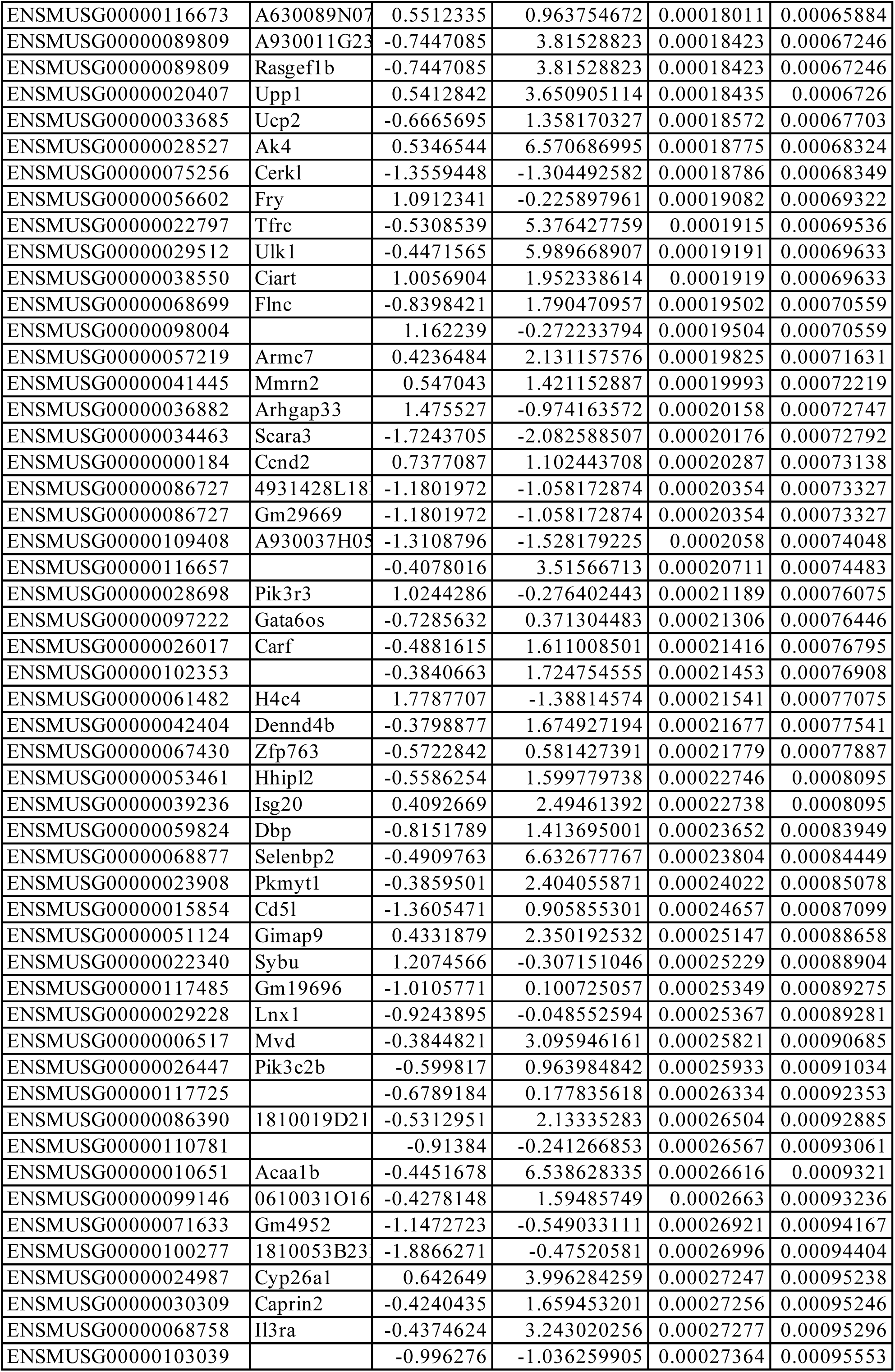
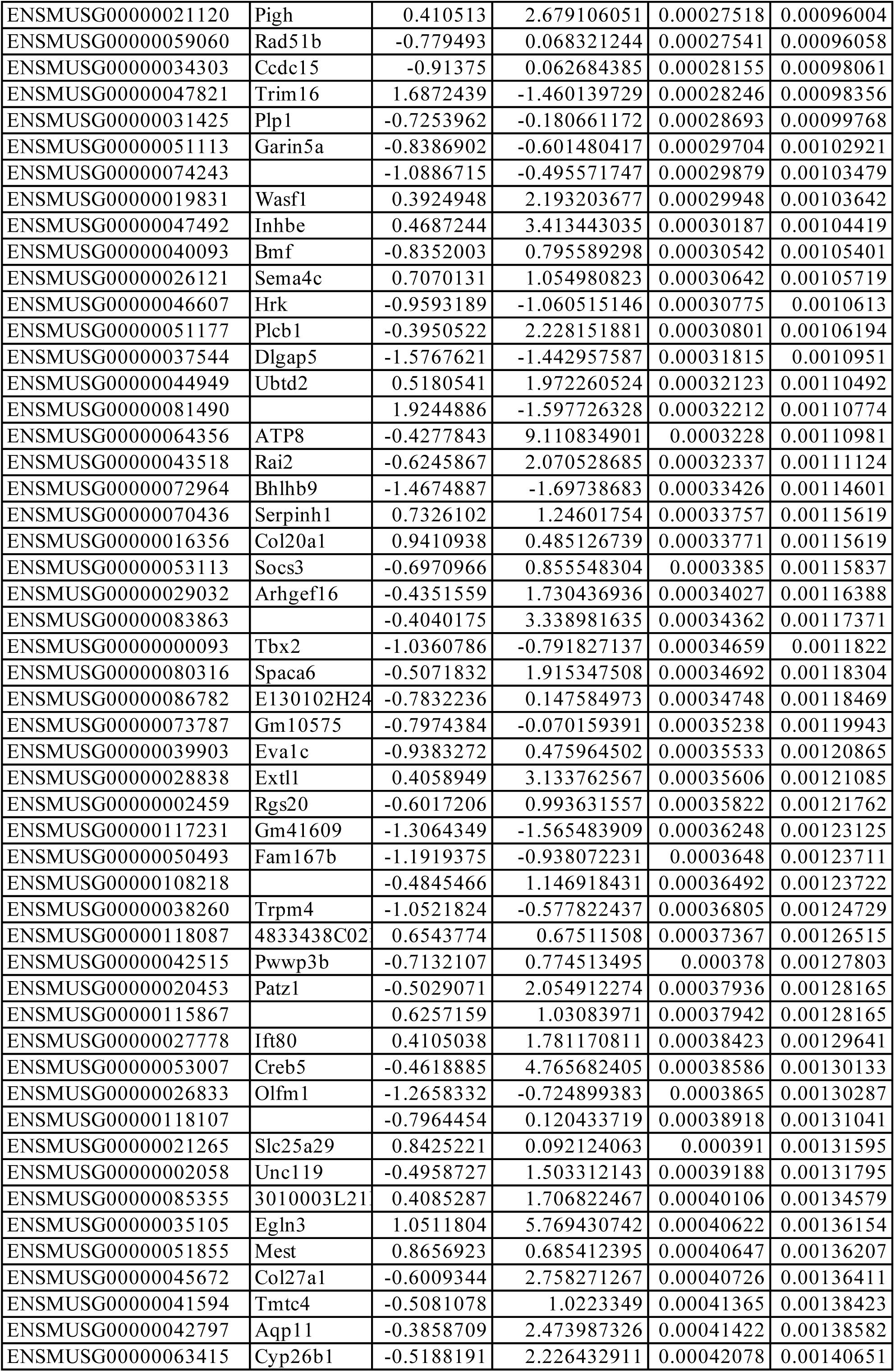
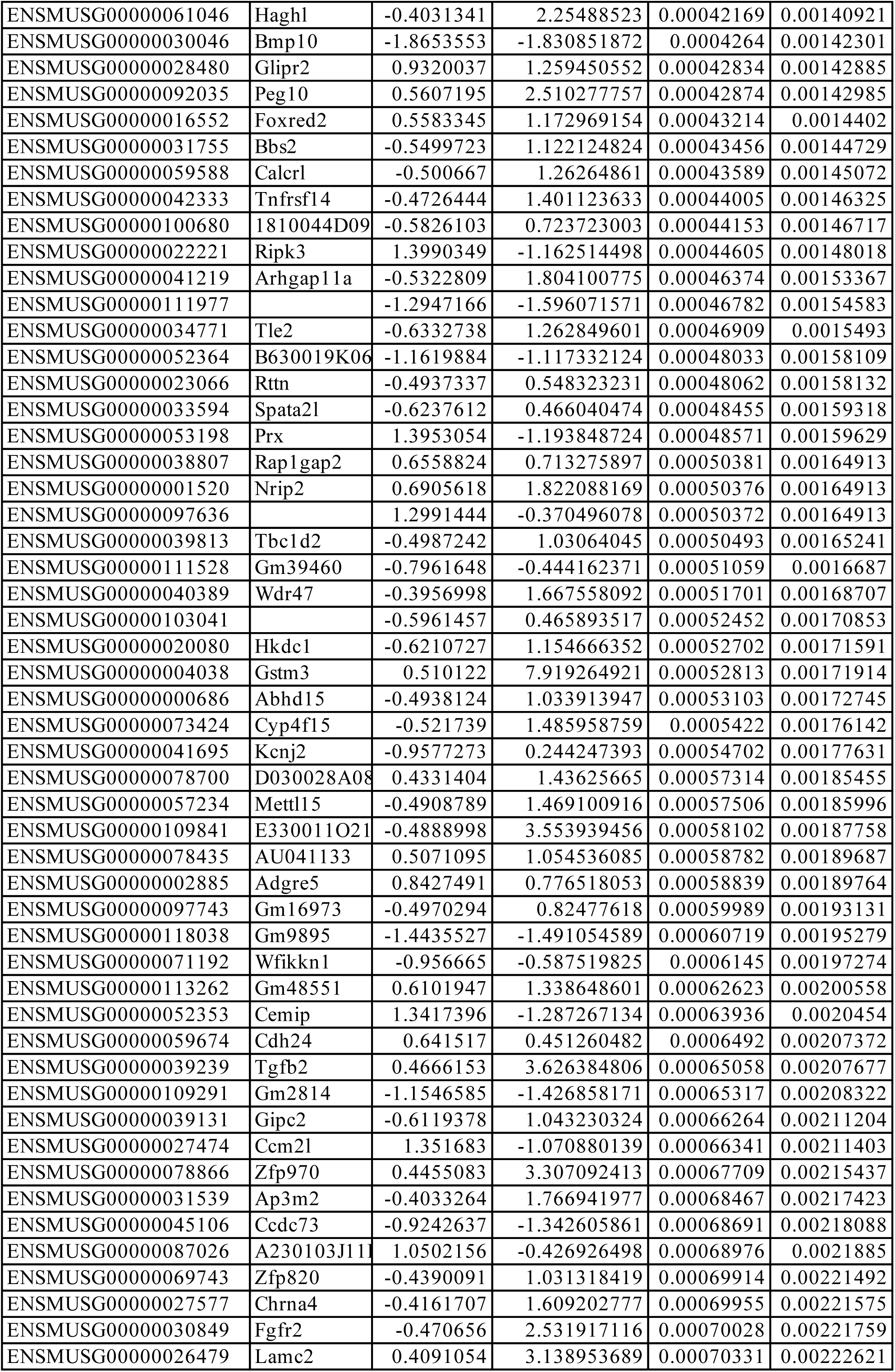
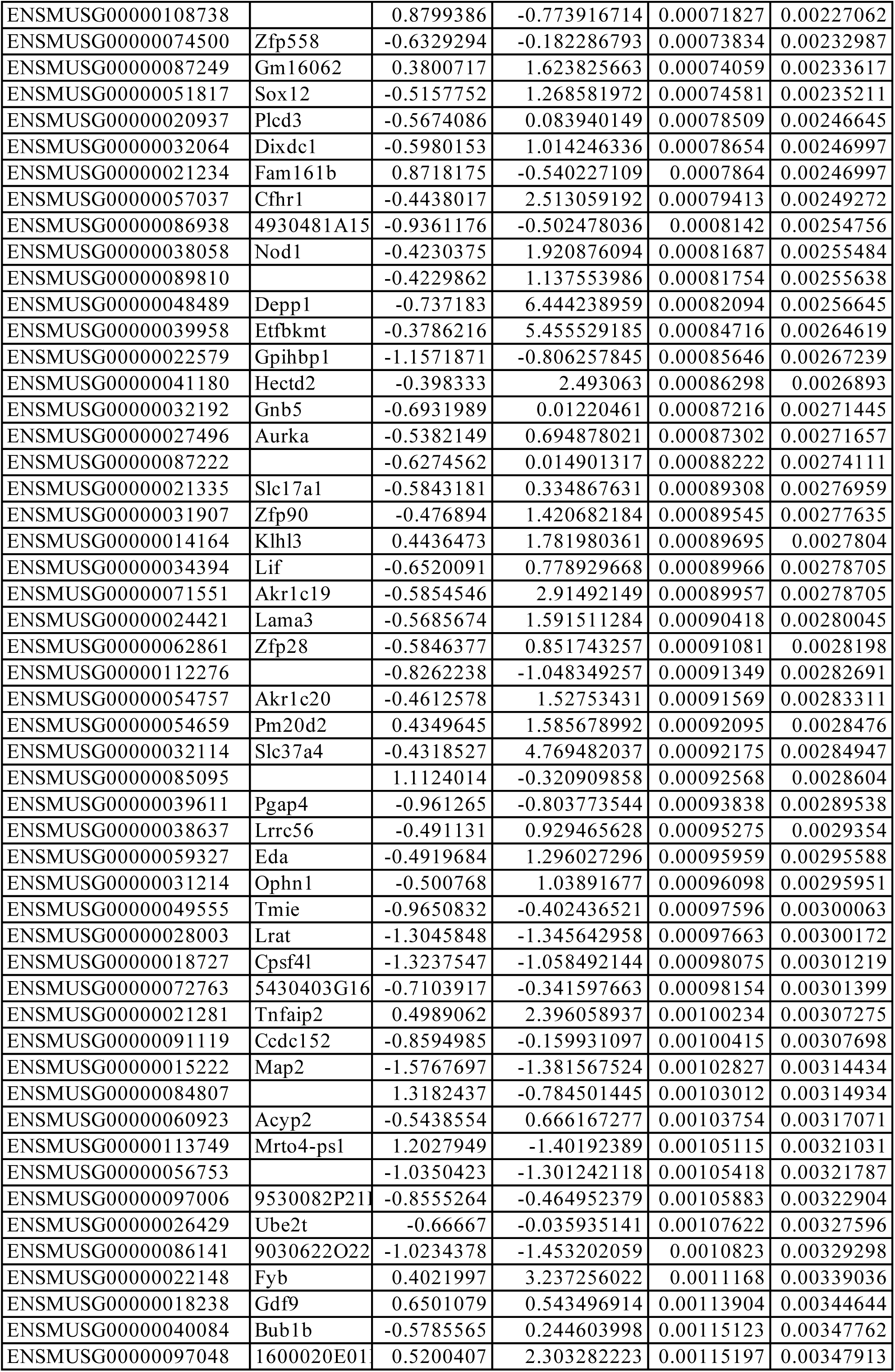
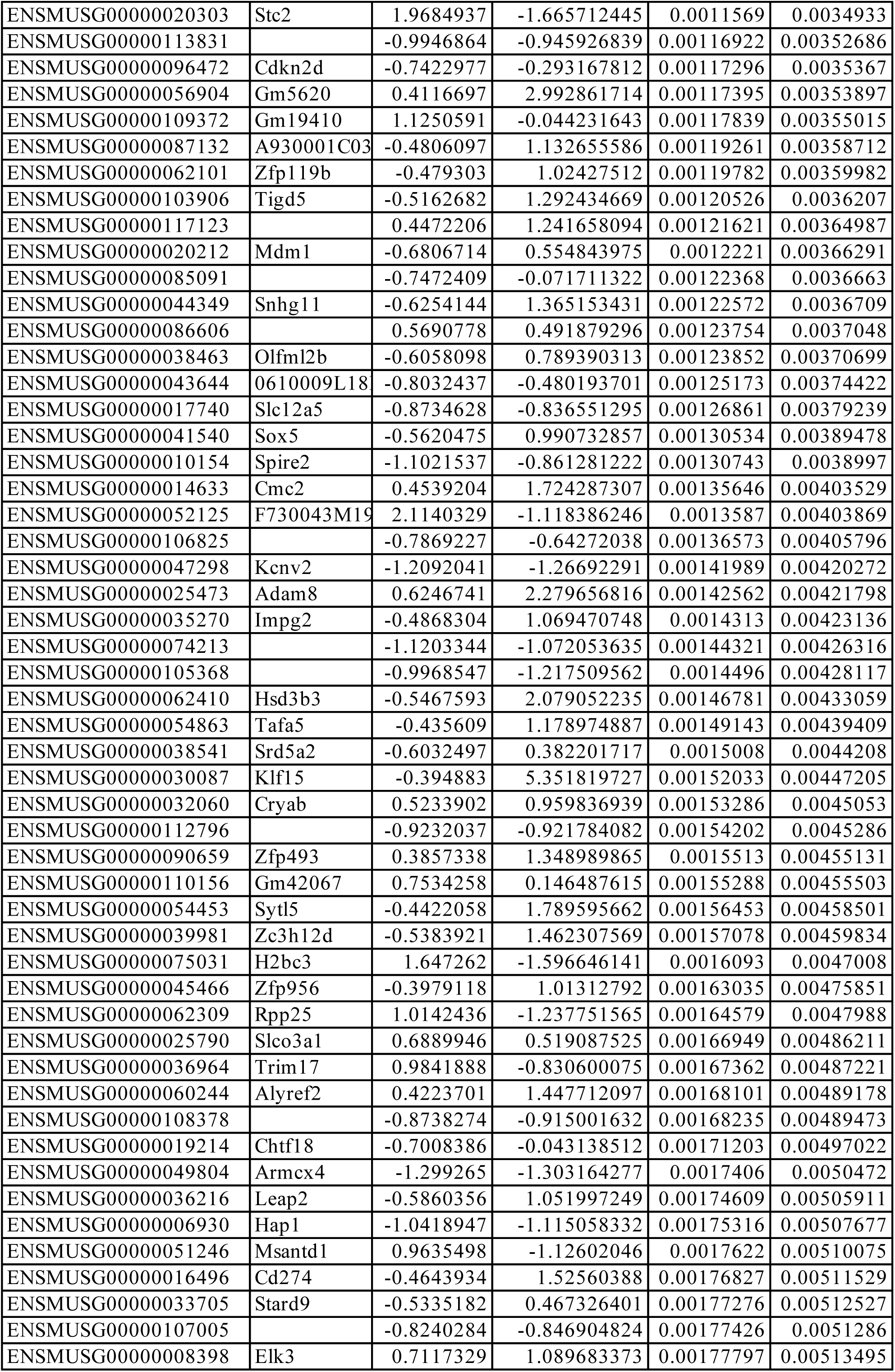
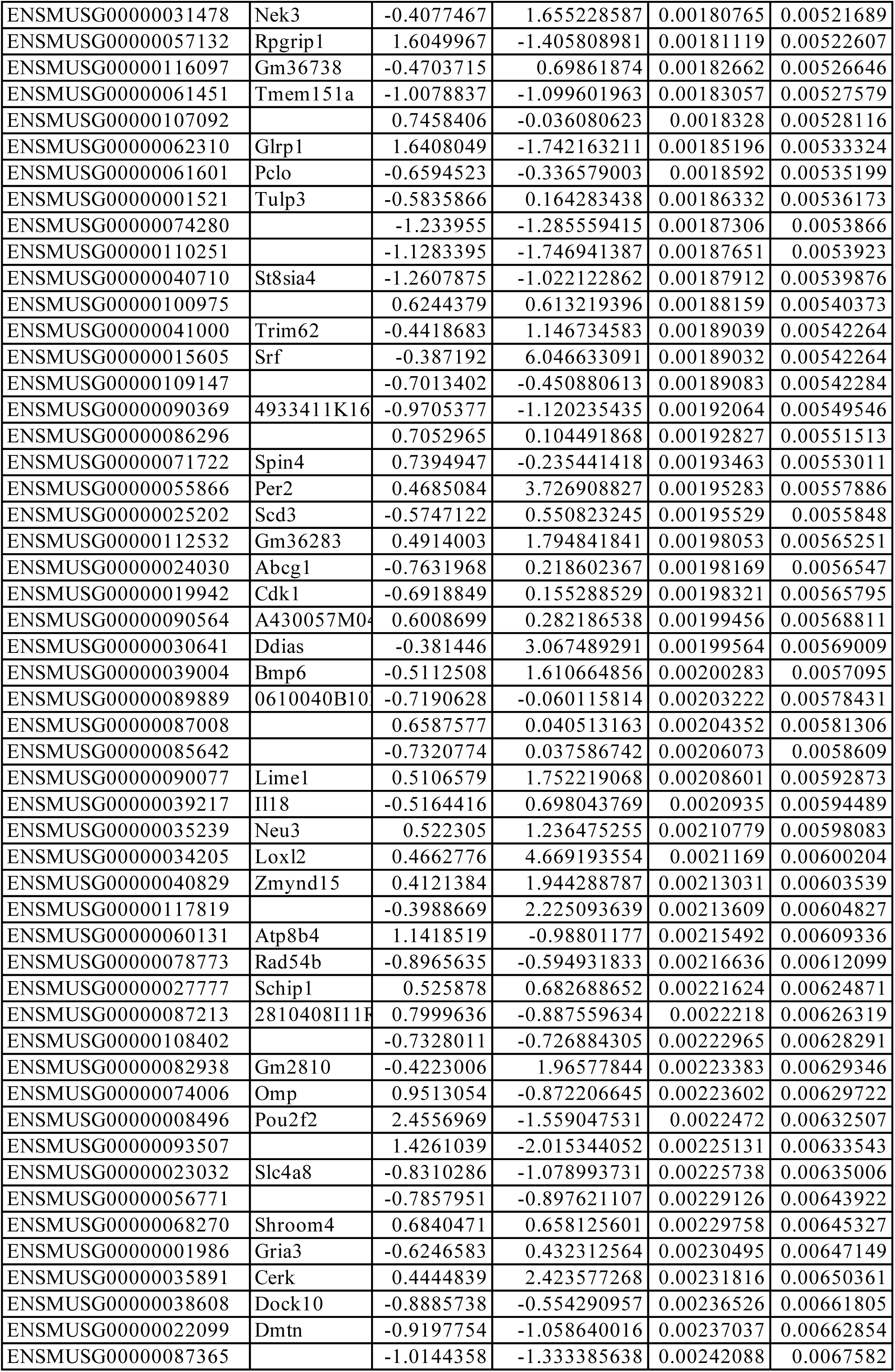
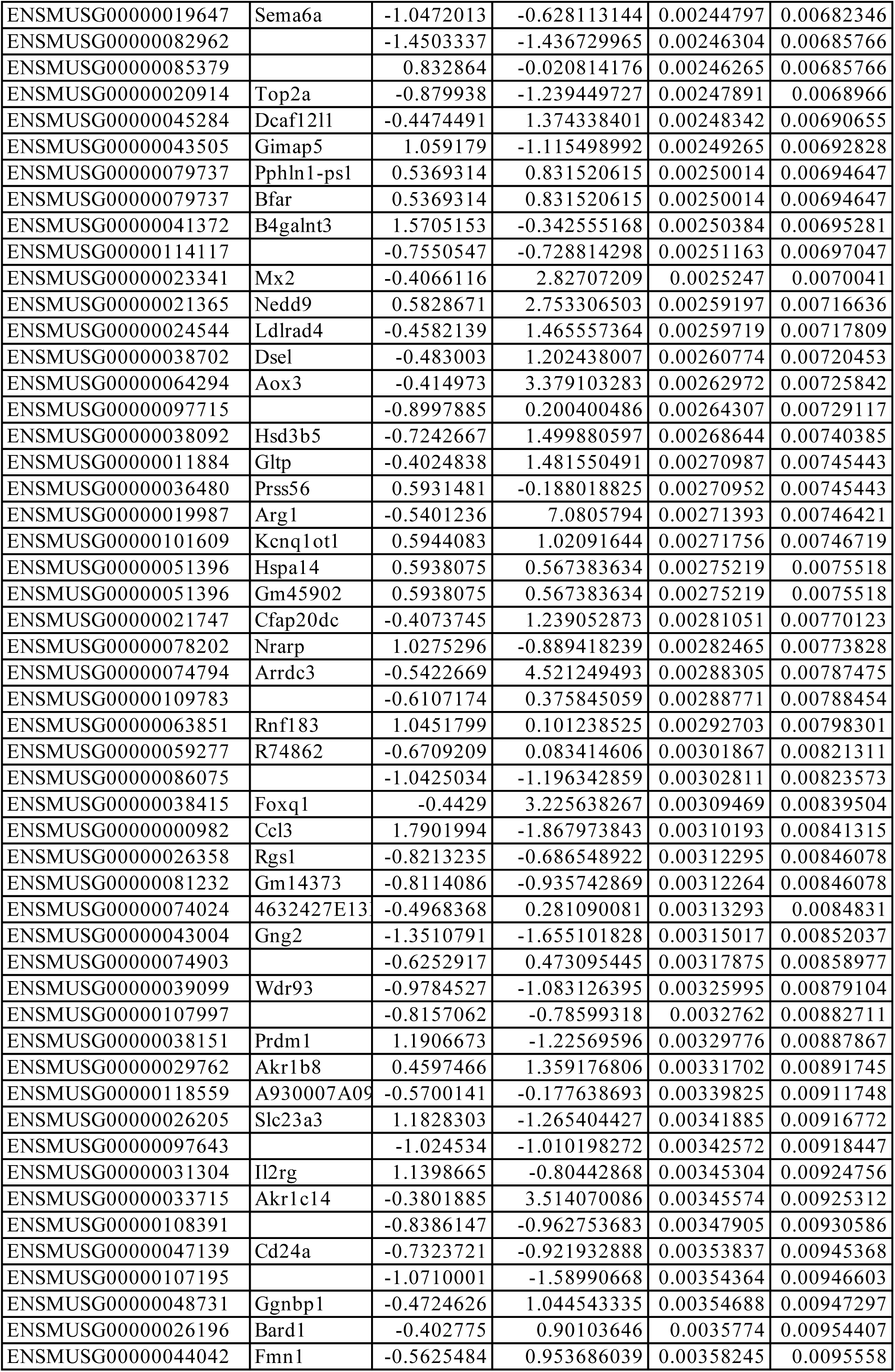
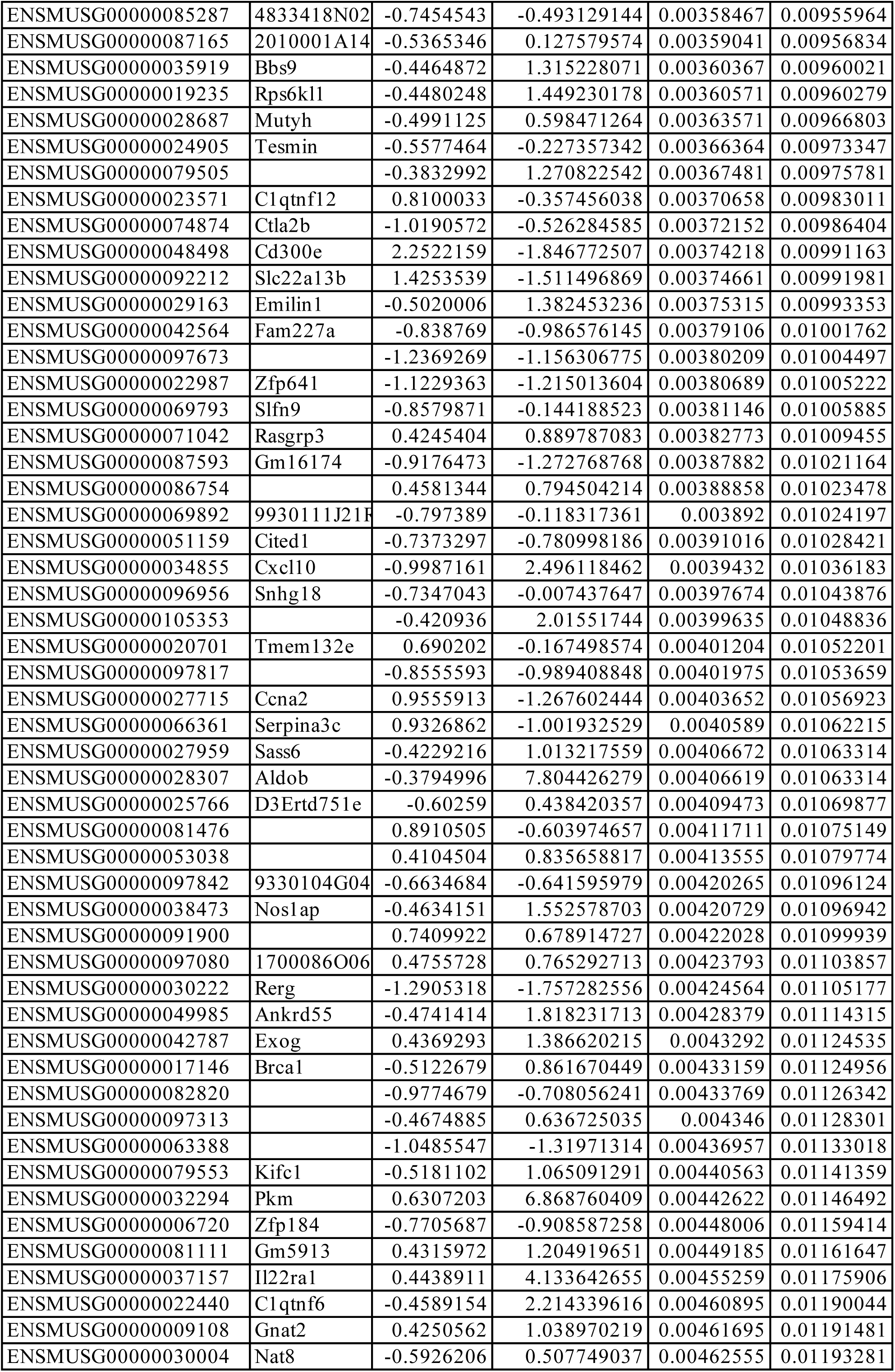
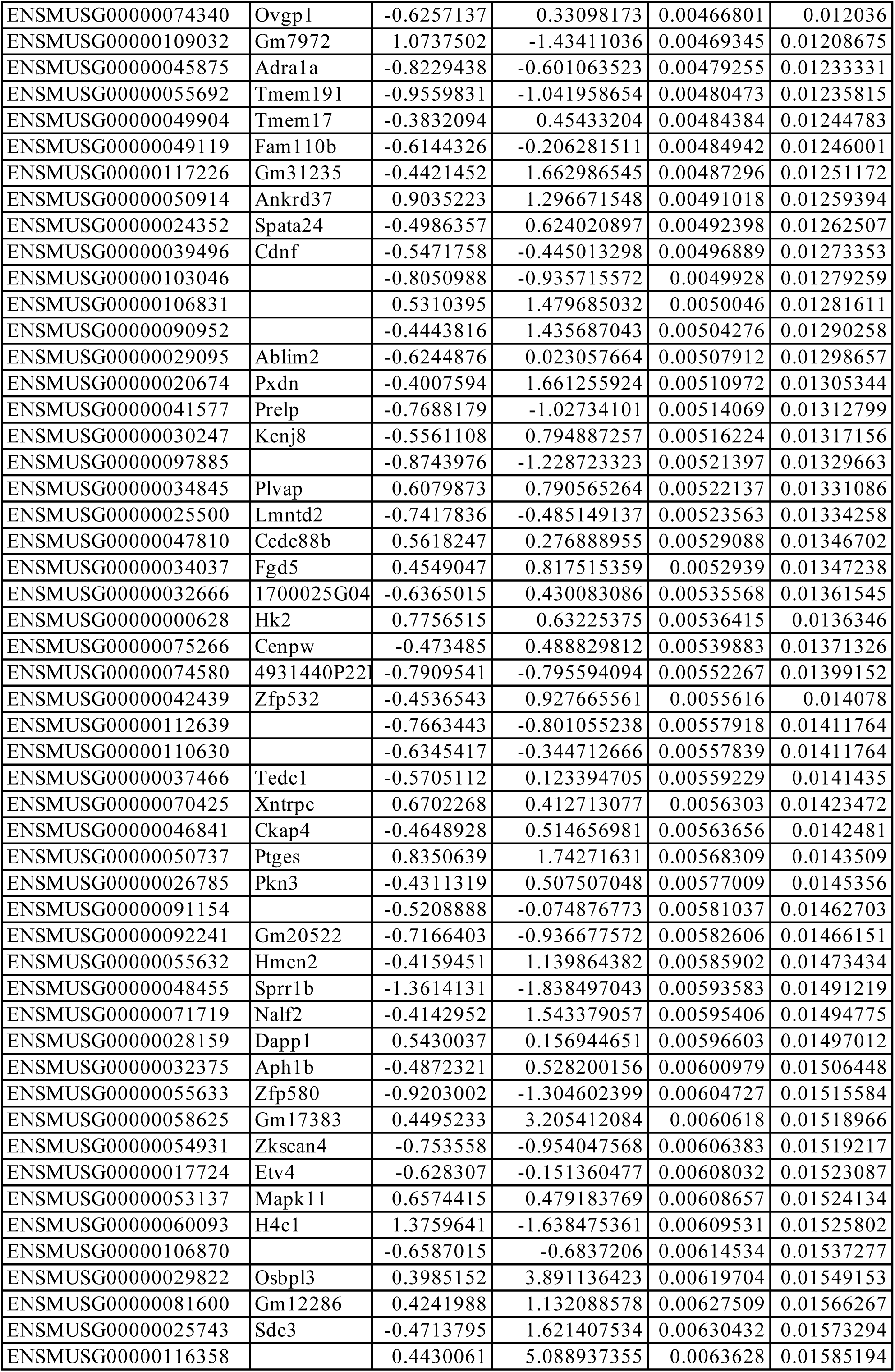
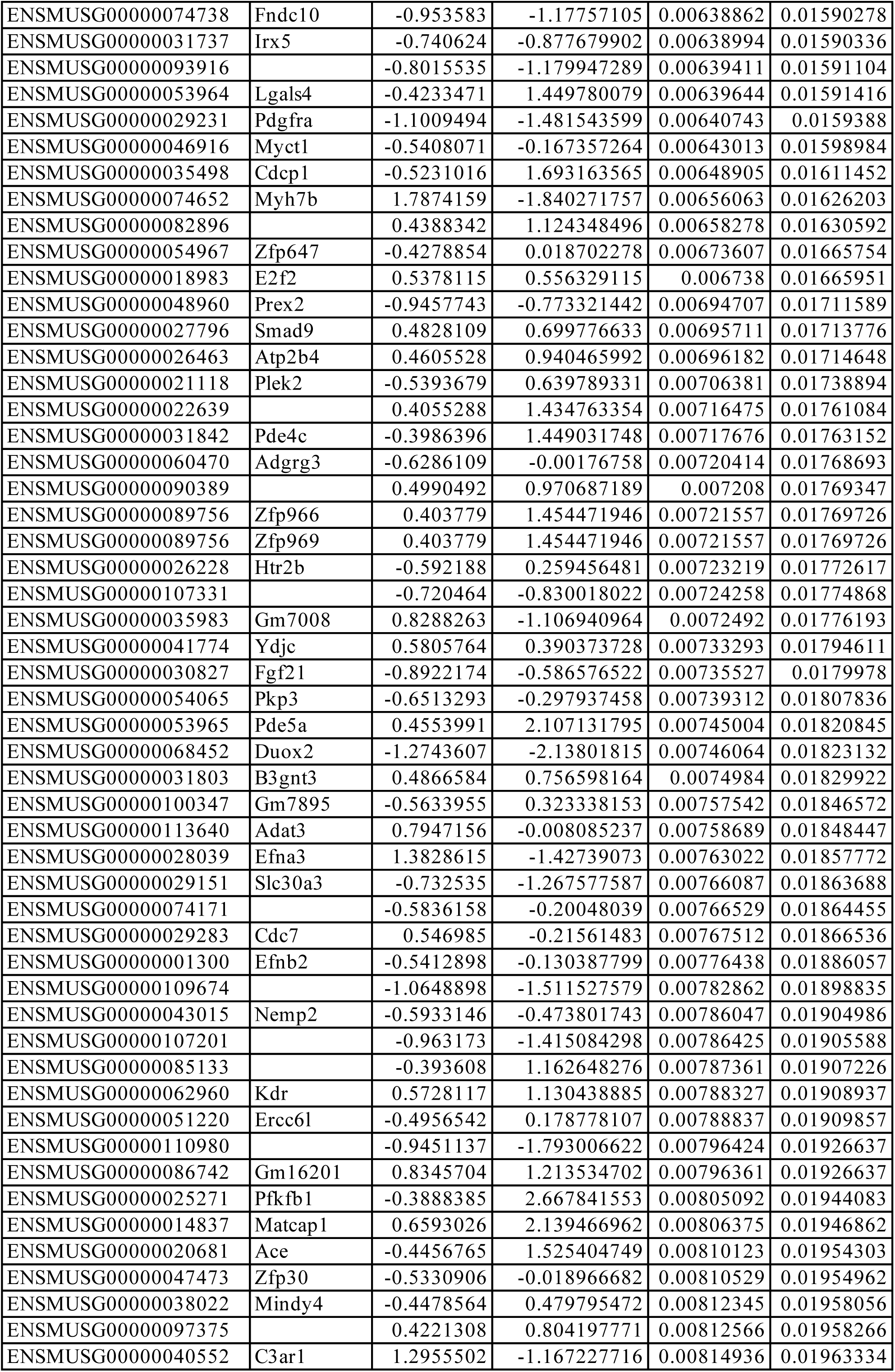
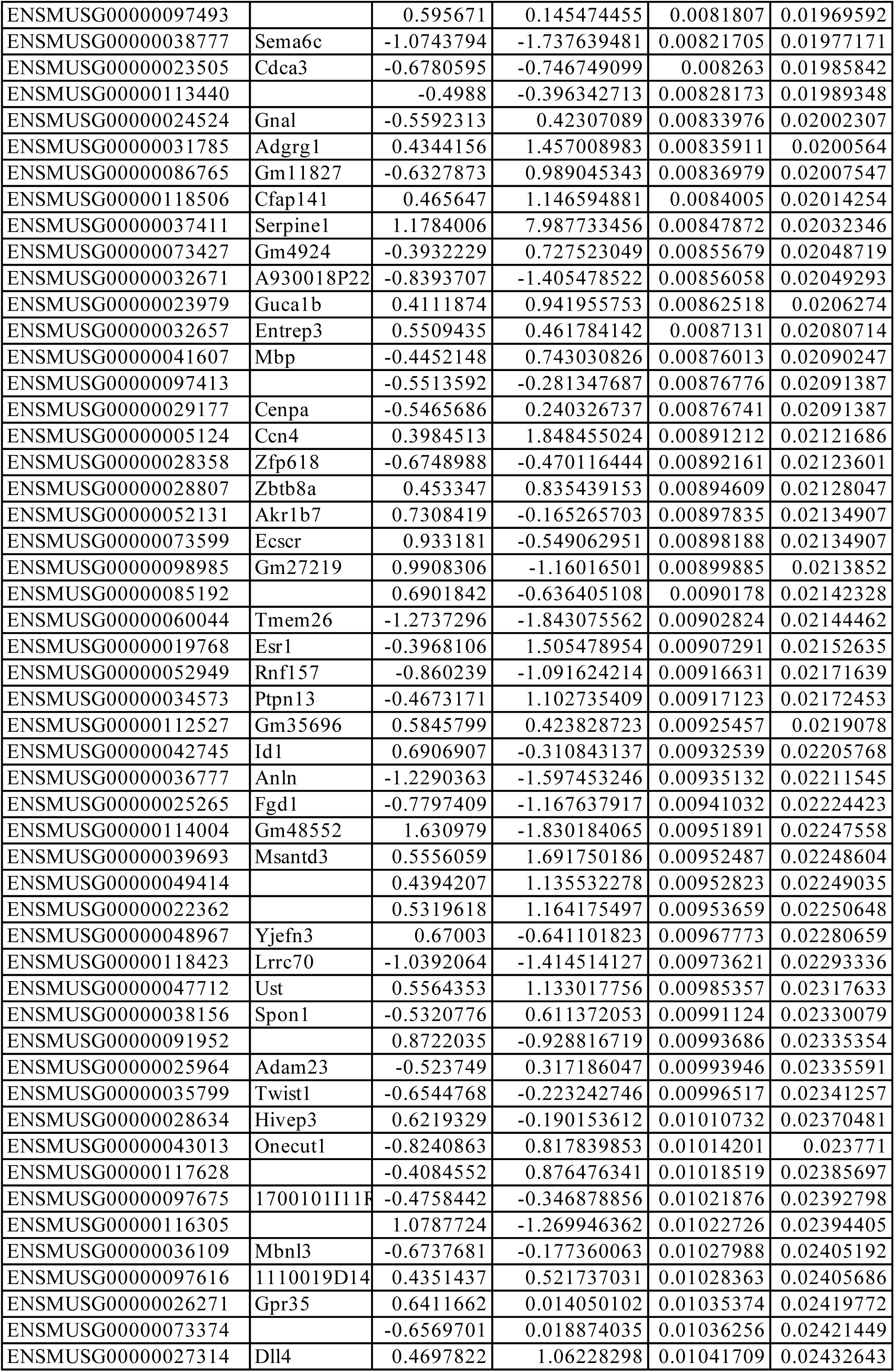
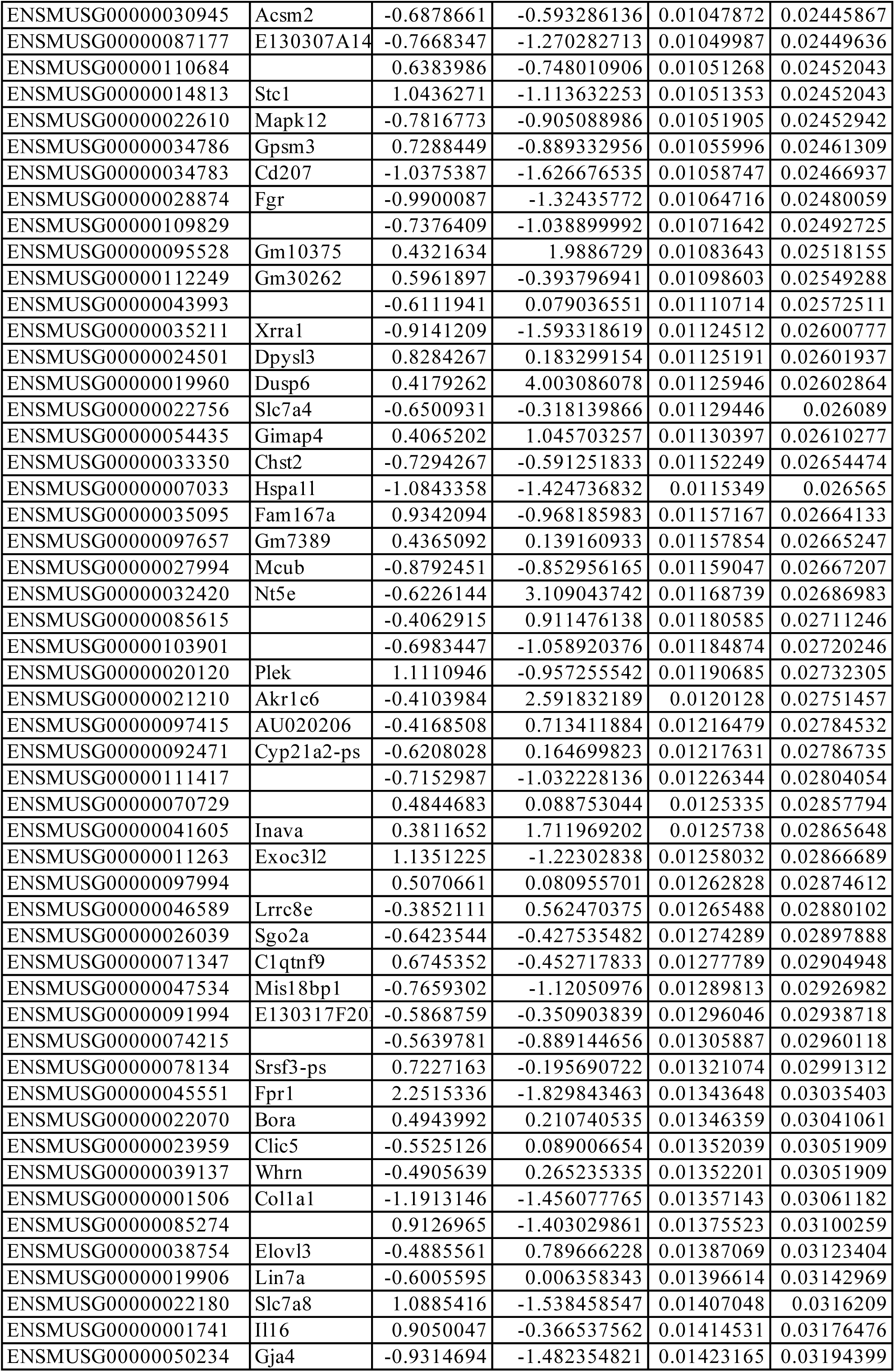
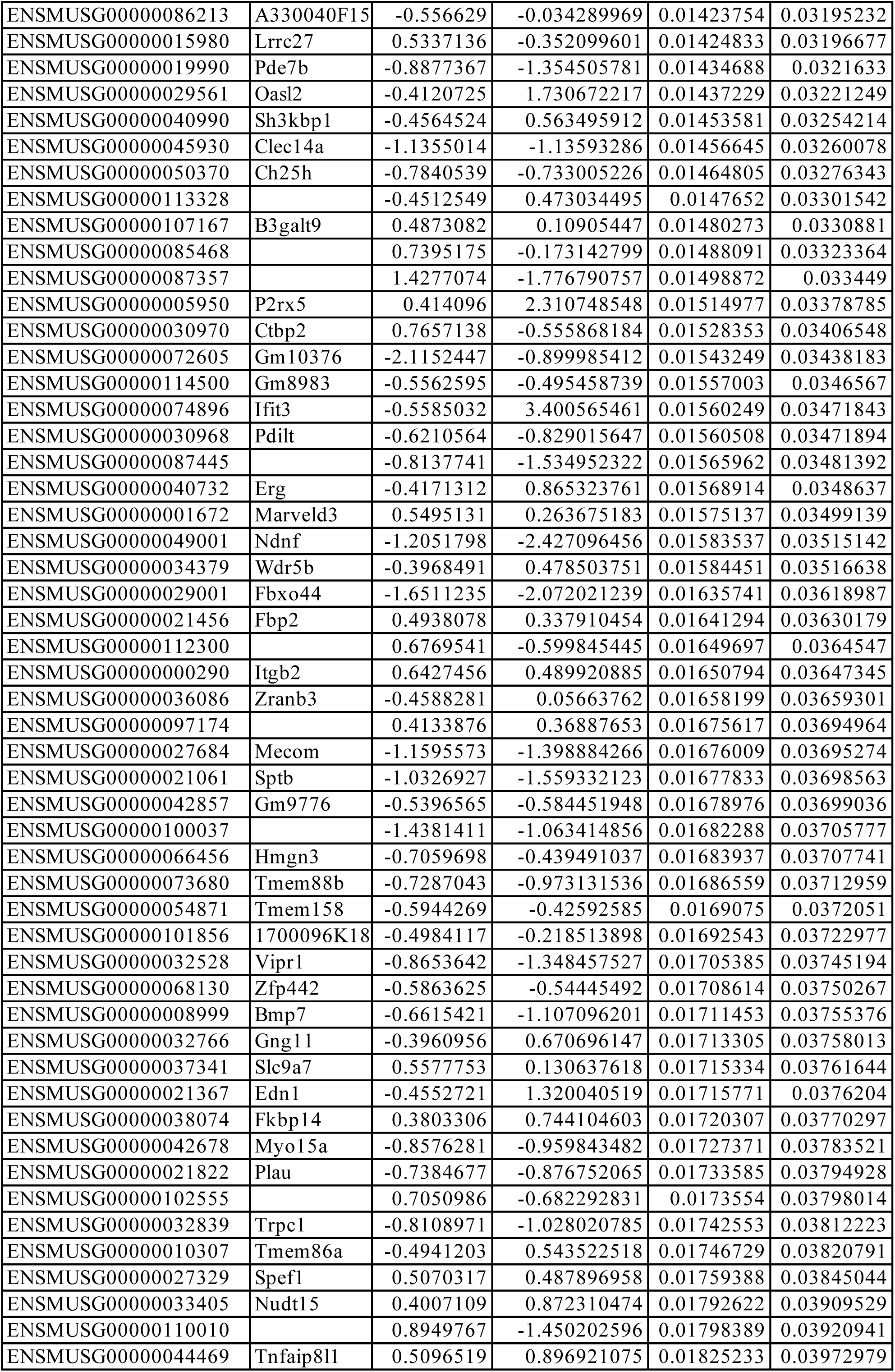
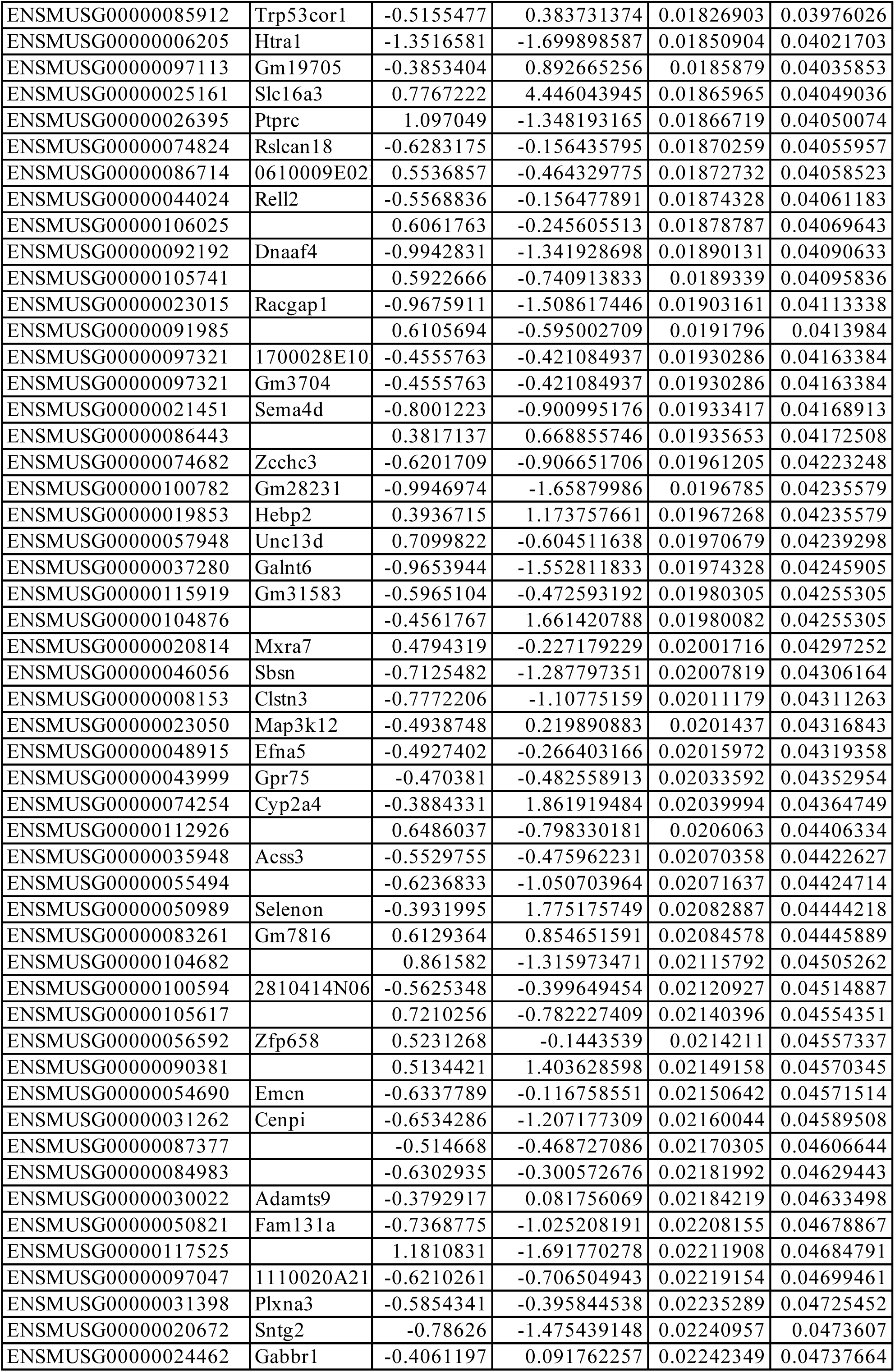
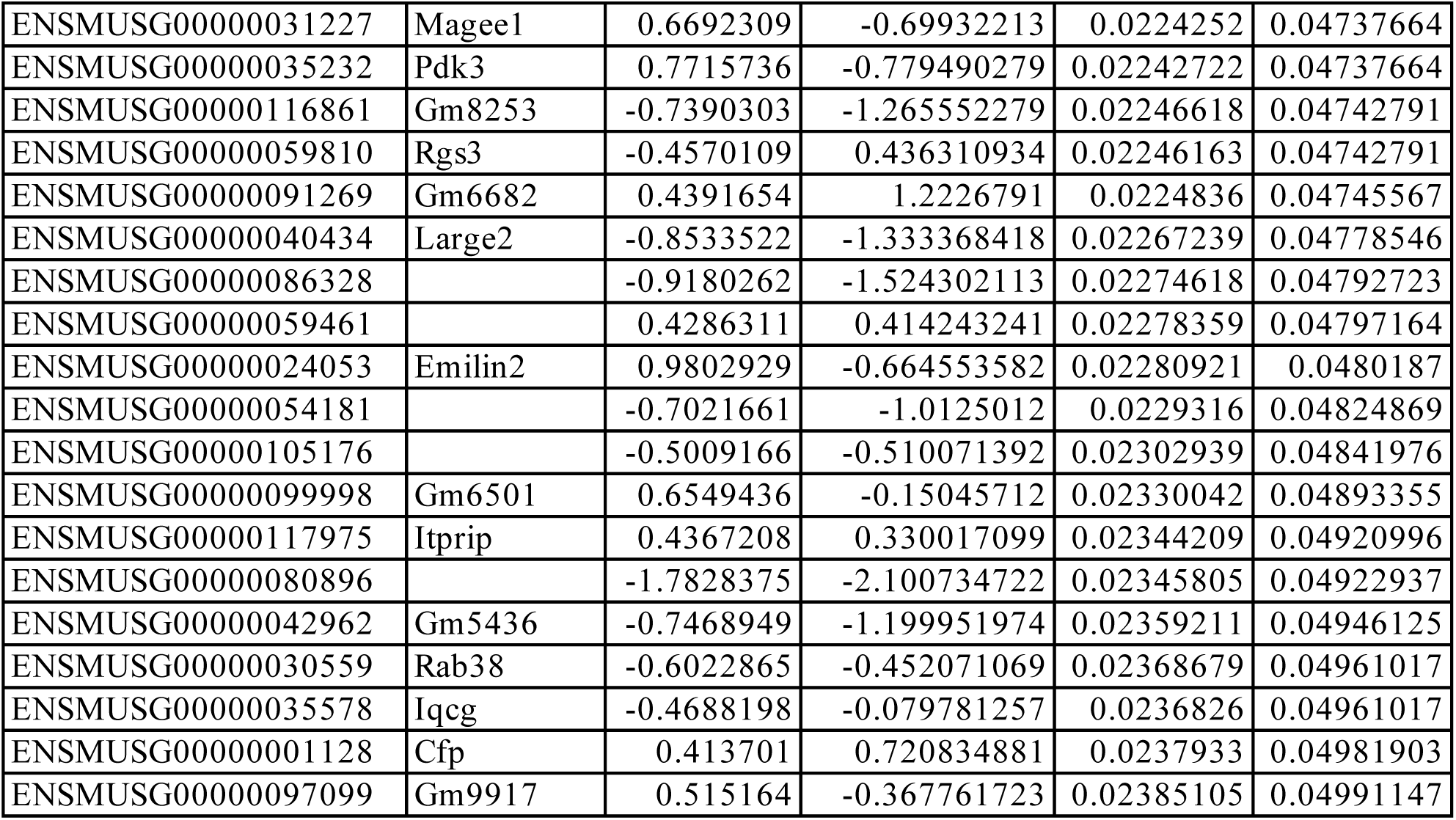
Significantly differentially expressed genes by BAY-3827 in the presence of MK-8722 in wild type primary hepatocytes. Significance set as abs (FC ≥ 1.3), FDR < 0.05.

## References

1. Hardie, D.G., Ross, F.A., and Hawley, S.A. (2012). AMPK: A nutrient and energy sensor that maintains energy homeostasis. Nature Reviews Molecular Cell Biology 13, 251–262.

2. Steinberg, G.R., and Hardie, D.G. (2023). New insights into activation and function of the AMPK. Nature Reviews Molecular Cell Biology 24, 255–272.

3. Hardie, D.G., and Sakamoto, K. (2006). AMPK: A key sensor of fuel and energy status in skeletal muscle. Physiology 21, 48–60.

4. Ross, F.A., MacKintosh, C., and Hardie, D.G. (2016). AMP-activated protein kinase: a cellular energy sensor that comes in 12 flavours. FEBS Journal 2987–3001.

5. Smiles, W.J., Ovens, A.J., Oakhil, J.S., Kofler, B. (2024). The metabolic sensor AMPK: Twelve enzymes in one. Molecular Metabolism 90.

6. Xiao, B., Sanders, M.J., Underwood, E., Heath, R., Mayer, F.V., Carmena, D., Jing, C., Walker, P.A., Eccleston, J.F., Haire, L.F., Saiu, P., Howell, S., Aasland, R., Martin, S., Carling, D., Gamblin, S.(2011). Structure of mammalian AMPK and its regulation by ADP. Nature 472, 230–233.

7. Langendorf, C.G., and Kemp, B.E. (2015). Choreography of AMPK activation. Cell Research 25, 5–6.

8. Steinberg, G.R., and Carling, D. (2019). AMP-activated protein kinase: the current landscape for drug development. Nature Reviews Drug Discovery 18, 527–551.

9. Carling, D., Zammit, V.A., and Hardie, D.G. (1987). A common bicyclic protein kinase cascade inactivates the regulatory enzymes of fatty acid and cholesterol biosynthesis. FEBS Letters 223, 217–222.

10. Hardie, D.G. (2022). AMP-activated protein kinase — a journey from 1 to 100 downstream targets. Biochemical Journal 479, 2327–2343.

11. Hardie, D.G., and Sakamoto, K. (2006). AMPK: A key sensor of fuel and energy status in skeletal muscle. Physiology 21, 48–60.

12. Sakamoto, K., and Holman, G.D. (2008). Emerging role for AS160/TBC1D4 and TBC1D1 in the regulation of GLUT4 traffic. American Journal of Physiology - Endocrinology and Metabolism 295.

13. Chen, Q., Xie, B., Zhu, S., Rong, P., Sheng, Y., Ducommun, S., Chen, L., Quan, C., Li, M., Sakamoto, K., MacKintosh, C., Chen, S., Wang, H. (2017). A Tbc1d1 Ser231Ala-knockin mutation partially impairs AICAR- but not exercise-induced muscle glucose uptake in mice. Diabetologia 60, 336–345.

14. Kjøbsted, R., Roll, J.L.W., Jørgensen, N.O., Birk, J.B., Foretz, M., Viollet, B., Chadt, A., Al-Hasani, H., and Wojtaszewski, J.F.P. (2019). AMPK and TBC1D1 regulate muscle glucose uptake after, but not during, exercise and contraction. Diabetes 68, 1427–1440.

15. Cool, B., Zinker, B., Chiou, W., Kifle, L., Cao, N., Perham, M., Dickinson, R., Adler, A., Gagne, G., Iyengar, R., Zhao, G., Marsh, K., Kym, P., Jung, P., Camp, H., Frevert, E. (2006). Identification and characterization of a small molecule AMPK activator that treats key components of type 2 diabetes and the metabolic syndrome. Cell Metabolism 3, 403–416.

16. Göransson, O., McBride, A., Hawley, S.A., Ross, F.A., Shpiro, N., Foretz, M., Viollet, B., Hardie, D.G., and Sakamoto, K. (2007). Mechanism of action of A-769662, a valuable tool for activation of AMP-activated protein kinase. Journal of Biological Chemistry 282, 32549– 32560.

17. Xiao, B., Sanders, M.J., Carmena, D., Bright, N.J., Haire, L.F., Underwood, E., Patel, B.R., Heath, R.B., Walker, P.A., Hallen, S., Giordanetto, F., Martin, S., Carling, D., Gamblin, S.(2013). Structural basis of AMPK regulation by small molecule activators. Nature Communications 4.

18. Myers, R.W., Guan, H.P., Ehrhart, J., Petrov, A., Prahalada, S., Tozzo, E., Yang, X., Kurtz, M.M., Trujillo, M., Trotter, D.G., Feng, D., Xu, S., Eiermann, G., Holahan, M. A., Rubins, D., Conarello, S., Niu, X., Souza, S. C., Miller, C., Liu, J., Lu, K., Feng, W., Li, Y., Painter, R., Milligan, J., He, H., Liu, F., Ogawa, A., Wisniewski, D., Rohm, R., Wang, L., Bunzel, M., Qian, Y., Zhu, W., Wang, H., Bennet, B., Scheuch, L., Fernandez, G., Li, C., Klimas, M., Zhou, G., Van Heek, M., Biftu, T., Weber, A., Kelley, D., Thornberry, N., Erion, M., Kemp, D., Sebhat, I. (2017). Systemic pan-AMPK activator MK-8722 improves glucose homeostasis but induces cardiac hypertrophy. Science 357, 507–511.

19. Cokorinos, E.C., Delmore, J., Reyes, A.R., Albuquerque, B., Kjøbsted, R., Jørgensen, N.O., Tran, J.L., Jatkar, A., Cialdea, K., Esquejo, R. M., Meissen, J., Calabrese, M. F., Cordes, J., Moccia, R., Tess, D., Salatto, C. T., Coskran, T. M., Opsahl, A. C., Flynn, D., Blatnik, M., Li, W., Kindt, E., Foretz, M., Viollet, B., Ward, J., Kurumbail, R., Kalgutkar, A., Wojtaszewski, J., Cameron, K., Miller, R. (2017). Activation of Skeletal Muscle AMPK Promotes Glucose Disposal and Glucose Lowering in Non-human Primates and Mice. Cell Metabolism 25, 1147–1159.e10.

20. Strang, J.E., Astridge, D.D., Nguyen, V.T., Reigan, P. (2025). Small Molecule Modulators of AMP-Activated Protein Kinase (AMPK) Activity and Their Potential in Cancer Therapy. Journal of Medicinal Chemistry 68.

21. Vara-Ciruelos, D., Russell, F.M., and Grahame Hardie, D. (2019). The strange case of AMPK and cancer: Dr Jekyll or Mr Hyde? Open Biology 9.

22. Alessi, D.R., Sakamoto, K., and Bayascas, J.R. (2006). LKB1-dependent signaling pathways. Annual Review of Biochemistry 75, 137–163.

23. Shaw, R.J., Bardeesy, N., Manning, B.D., Lopez, L., Kosmatka, M., DePinho, R.A., and Cantley, L.C. (2004). The LKB1 tumor suppressor negatively regulates mTOR signaling. Cancer Cell 6, 91–99.

24. Huang, X., Wullschleger, S., Shpiro, M., McGuire, V.A., Sakamoto, K., Woods, Y.L., McBurnie, W., Fleming, S., and Alessi, D.R. (2008). Important role of the LKB1-AMPK pathway in suppressing tumorigenesis in PTEN-deficient mice. Biochemical Journal 412, 211–221.

25. Russell, F.M., and Hardie, D.G. (2021). AMP-activated protein kinase: Do we need activators or inhibitors to treat or prevent cancer? International Journal of Molecular Sciences 22, 1–30.

26. Monteverde, T., Muthalagu, N., Port, J., and Murphy, D.J. (2015). Evidence of cancer-promoting roles for AMPK and related kinases. FEBS Journal 282, 4658–4671.

27. Zhou, G., Myers, R., Li, Y., Chen, Y., Shen, X., Fenyk-Melody, J., Wu, M., Ventre, J., Doebber, T., Fujii, N., Musi, N., Hirshman, M. F., Goodyear, L. J., and Moller, D. E. (2001). Role of AMP-activated protein kinase in mechanism of metformin action. Journal of Clinical Investigation 108, 1167–1174.

28. Yu, P.B., Hong, C.C., Sachidanandan, C., Babitt, J.L., Deng, D.Y., Hoyng, S.A., Lin, H.Y., Bloch, K.D., and Peterson, R.T. (2008). Dorsomorphin inhibits BMP signals required for embryogenesis and iron metabolism. Nature Chemical Biology 4, 33–41.

29. Handa, N., Takagi, T., Saijo, S., Kishishita, S., Takaya, D., Toyama, M., Terada, T., Shirouzu, M., Suzuki, A., Lee, S., Yamauchi, T., Okada-Iwabu, M., Iwabu, M., Kadowaki, T., Minokoshi, Y., Yokoyama, S. (2011). Structural basis for compound C inhibition of the human AMP-activated protein kinase α2 subunit kinase domain. Acta Crystallographica Section D: Biological Crystallography 67, 480–487.

30. 30. Chaikuad, A., Alfano, I., Kerr, G., Sanvitale, C.E., Boergermann, J.H., Triffitt, J.T., Von Delft, F., Knapp, S., Knaus, P., and Bullock, A.N. (2012). Structure of the bone morphogenetic protein receptor ALK2 and implications for fibrodysplasia ossificans progressiva. Journal of Biological Chemistry 287, 36990–36998.

31. Bain, J., Plater, L., Elliott, M., Shpiro, N., Hastie, C.J., Mclauchlan, H., Klevernic, I., Arthur, J.S.C., Alessi, D.R., and Cohen, P. (2007). The selectivity of protein kinase inhibitors: A further update. Biochemical Journal 408, 297–315.

32. Vogt, J., Traynor, R., and Sapkota, G.P. (2011). The specificities of small molecule inhibitors of the TGFß and BMP pathways. Cellular Signalling 23, 1831–1842.

33. Dite, T. A., Langendorf, C. G., Hoque, A., Galic, S., Rebello, R. J., Ovens, A. J., Lindqvist, L. M., Ngoei, K. R. W., Ling, N. X. Y., Furic, L., Kemp, B. E., Scott, J. W., Oakhill, J. S. (2018). AMP-activated protein kinase selectively inhibited by the type II inhibitor SBI-0206965. Journal of Biological Chemistry 293, 8874–8885.

34. 34. Dasgupta, B., and Seibel, W. (2018). Compound C/Dorsomorphin: Its use and misuse as an AMPK inhibitor. Methods in Molecular Biology, (Humana Press Inc.), pp. 195–202.

35. Egan, D.F., Chun, M.G.H., Vamos, M., Zou, H., Rong, J., Miller, C.J., Lou, H.J., Raveendra-Panickar, D., Yang, C.C., Sheffler, D.J., Teriete, P., Asara, J., Turk, B., Cosford, N., Shaw, R.(2015). Small Molecule Inhibition of the Autophagy Kinase ULK1 and Identification of ULK1 Substrates. Molecular Cell 59, 285–297.

36. Ahwazi, D., Neopane, K., Markby, G.R., Kopietz, F., Ovens, A.J., Dall, M., Hassing, A.S., Gräsle, P., Alshuweishi, Y., Treebak, J.T., Salt, I., Göransson, O., Zeqiraj, E., Scott, J., Sakamoto, K. (2021). Investigation of the specificity and mechanism of action of the ULK1/AMPK inhibitor SBI-0206965. Biochemical Journal 478, 2977–2997.

37. Lemos, C., Schulze, V.K., Baumgart, S.J., Nevedomskaya, E., Heinrich, T., Lefranc, J., Bader, B., Christ, C.D., Briem, H., Kuhnke, L.P., Holton, S., Bömer, U., Lienau, P., von Nussbaum, F., Nising, C., Bauser, M., Hägebarth, A., Mumberg, D., Haendler, B.(2021). The potent AMPK inhibitor BAY-3827 shows strong efficacy in androgen-dependent prostate cancer models. Cellular Oncology 44, 581–594.

38. Manning, G., Whyte, D.B., Martinez, R., Hunter, T., and Sudarsanam, S. (2002). The protein kinase complement of the human genome. Science 298, 1912–1934.

39. Ashkenazy, H., Abadi, S., Martz, E., Chay, O., Mayrose, I., Pupko, T., and Ben-Tal, N. (2016). ConSurf 2016: an improved methodology to estimate and visualize evolutionary conservation in macromolecules. Nucleic Acids Research, 44.

40. Yariv, B., Yariv, E., Kessel, A., Masrati, G., Chorin, A.B., Martz, E., Mayrose, I., Pupko, T., and Ben-Tal, N. (2023). Using evolutionary data to make sense of macromolecules with a “face-lifted” ConSurf. Protein Science 32.

41. Langendorf, C.G., Ngoei, K.R.W., Scott, J.W., Ling, N.X.Y., Issa, S.M.A., Gorman, M.A., Parker, M.W., Sakamoto, K., Oakhill, J.S., and Kemp, B.E. (2016). Structural basis of allosteric and synergistic activation of AMPK by furan-2-phosphonic derivative C2 binding. Nature Communications 7.

42. Modi, V., and Dunbrack, R.L. (2019). Defining a new nomenclature for the structures of active and inactive kinases. Proceedings of the National Academy of Sciences of the United States of America 116, 6818–6827.

43. Möbitz, H. (2015). The ABC of protein kinase conformations. Biochimica et Biophysica Acta - Proteins and Proteomics 1854, 1555–1566.

44. Taylor, S.S., Keshwani, M.M., Steichen, J.M., and Kornev, A.P. (2012). Evolution of the eukaryotic protein kinases as dynamic molecular switches. Philosophical Transactions of the Royal Society B: Biological Sciences 367, 2517–2528.

45. Sanders, M.J., Ratinaud, Y., Neopane, K., Bonhoure, N., Day, E.A., Ciclet, O., Lassueur, S., Pinta, M.N., Deak, M., Brinon, B., Christen, S., Steinberg, G., Barron, D., Sakamoto, K. (2022). Natural (dihydro)phenanthrene plant compounds are direct activators of AMPK through its allosteric drug and metabolite–binding site. Journal of Biological Chemistry 298.

46. Hawley, S.A., Russell, F.M., Ross, F.A., and Hardie, D.G. (2024). BAY-3827 and SBI- 0206965: Potent AMPK Inhibitors That Paradoxically Increase Thr172 Phosphorylation. International Journal of Molecular Sciences 25.

47. Boudaba, N., Marion, A., Huet, C., Pierre, R., Viollet, B., and Foretz, M. (2018). AMPK Re-Activation Suppresses Hepatic Steatosis but its Downregulation Does Not Promote Fatty Liver Development. EBioMedicine 28, 194–209.

48. Hunter, R.W., Foretz, M., Bultot, L., Fullerton, M.D., Deak, M., Ross, F.A., Hawley, S.A., Shpiro, N., Viollet, B., Barron, D., Kemp, B., Steinberg, G., Hardie, D., Sakamoto, K.(2014). Mechanism of action of compound-13: An α1-selective small molecule activator of AMPK. Chemistry and Biology 21, 866–879.

49. Hashimoto, T., Urushihara, Y., Murata, Y., Fujishima, Y., and Hosoi, Y. (2022). AMPK increases expression of ATM through transcriptional factor Sp1 and induces radioresistance under severe hypoxia in glioblastoma cell lines. Biochemical and Biophysical Research Communications 590, 82–88.

50. Rao, X.S., Cong, X.X., Gao, X.K., Shi, Y.P., Shi, L.J., Wang, J.F., Ni, C.Y., He, M.J., Xu, Y., Yi, C., Meng, Z., Liu, J., Lin, P., Zheng, L., Zhou Y.(2021). AMPK-mediated phosphorylation enhances the auto-inhibition of TBC1D17 to promote Rab5-dependent glucose uptake. Cell Death and Differentiation 28, 3214–3234.

51. 51. Yamano, K., Fogel, A.I., Wang, C., van der Bliek, A.M., and Youle, R.J. (2014). Mitochondrial Rab GAPs govern autophagosome biogenesis during mitophagy. ELife 3.

52. Yang, W.S., Caliva, M.J., Khadka, V.S., Tiirikainen, M., Matter, M.L., Deng, Y., and Ramos, J.W. (2023). RSK1 and RSK2 serine/threonine kinases regulate different transcription programs in cancer. Frontiers in Cell and Developmental Biology 10.

53. Sapkota, G.P., Cummings, L., Newell, F.S., Armstrong, C., Bain, J., Frodin, M., Grauert, M., Hoffmann, M., Schnapp, G., Steegmaier, M., Cohen, P., Alessi, D.(2007). BI-D1870 is a specific inhibitor of the p90 RSK (ribosomal S6 kinase) isoforms in vitro and in vivo. Biochemical Journal 401, 29–38.

54. Utepbergenov, D., Derewenda, U., Olekhnovich, N., Szukalska, G., Banerjee, B., Hilinski, M.K., Lannigan, D.A., Stukenberg, P.T., and Derewenda, Z.S. (2012). Insights into the inhibition of the p90 ribosomal S6 kinase (RSK) by the flavonol glycoside SL0101 from the 1.5 Å crystal structure of the N-terminal domain of RSK2 with bound inhibitor. Biochemistry 51, 6499–6510.

55. Jain, R., Mathur, M., Lan, J., Costales, A., Atallah, G., Ramurthy, S., Subramanian, S., Setti, L., Feucht, P., Warne, B., Doyle, L., Basham, S., Jefferson, A., Lindvall, M., Appleton, B., Shafer C. (2015). Discovery of Potent and Selective RSK Inhibitors as Biological Probes. Journal of Medicinal Chemistry 58, 6766–6783.

56. Murray, B.W., Guo, C., Piraino, J., Westwick, J.K., Zhang, C., Lamerdin, J., Dagostino, E., Knighton, D., Loi, C.M., Zager, M., Kraynov, E., Popoff, I., Christensen, J., Martinez, R., Kephart, S., Marakovits, J., Karlicek, S., Bergqvist, S., Smeal, T. (2010). Small-molecule p21-activated kinase inhibitor PF-3758309 is a potent inhibitor of oncogenic signaling and tumor growth. Proceedings of the National Academy of Sciences of the United States of America 107, 9446–9451.

57. Schneider, C., Hilbert, J., Genevaux, F., Höfer, S., Krauß, L., Schicktanz, F., Contreras, C.T., Jansari, S., Papargyriou, A., Richter, T., Alfayomy, A.M., Falcomatà, C., Schneeweis, C, Orben, F., Öllinger, R., Wegwitz, F., Boshnakovska, A., Rehling, P., Müller, D., Ströbel, P., Ellenrieder, V., Conradi, L., Hessmann, E., Ghadimi, M., Grade, M., Wirth, M., Steiger, K., Rad, R., Kuster, B., Sippl, W., Reichert, M., Saur, D., Schneider, G. (2024). A Novel AMPK Inhibitor Sensitizes Pancreatic Cancer Cells to Ferroptosis Induction. Advanced Science 11.

58. Anjum, R., and Blenis, J. (2008). The RSK family of kinases: Emerging roles in cellular signalling. Nature Reviews Molecular Cell Biology 9, 747–758.

59. Koutsougianni, F., Alexopoulou, D., Uvez, A., Lamprianidou, A., Sereti, E., Tsimplouli, C., Ilkay Armutak, E., and Dimas, K. (2023). P90 ribosomal S6 kinases: A bona fide target for novel targeted anticancer therapies? Biochemical Pharmacology 210.

60. Viollet, B., and Foretz, M. (2016). Animal Models to Study AMPK. EXS 107, 441–469.

61. Ashraf N, Van Nostrand JL. Fine-tuning AMPK in physiology and disease using point-mutant mouse models. Disease Models & Mechanisms 17.

62. Sakamoto, K., McCarthy, A., Smith, D., Green, K.A., Hardie, D.G., Ashworth, A., and Alessi, D.R. (2005). Deficiency of LKB1 in skeletal muscle prevents AMPK activation and glucose uptake during contraction. EMBO Journal 24, 1810–1820.

63. Hastie, C.J., McLauchlan, H.J., and Cohen, P. (2006). Assay of protein kinases using radiolabeled ATP: A protocol. Nature Protocols 1, 968–971.

64. Dite, T.A., Ling, N.X.Y., Scott, J.W., Hoque, A., Galic, S., Parker, B.L., Ngoei, K.R.W., Langendorf, C.G., O’Brien, M.T., Kundu, M., et al. (2017). The autophagy initiator ULK1 sensitizes AMPK to allosteric drugs. Nature Communications 8.

65. Scott, J.W., Park, E., Rodriguiz, R.M., Oakhill, J.S., Issa, S.M.A., Obrien, M.T., Dite, T.A., Langendorf, C.G., Wetsel, W.C., Means, A.R., et al. (2015). Autophosphorylation of CaMKK2 generates autonomous activity that is disrupted by a T85S mutation linked to anxiety and bipolar disorder. Scientific Reports 5.

66. 66. Ewels, P. A., Peltzer, A., Fillinger, S., Patel, H., Alneberg, J., Wilm, A., Garcia, M. U., Di Tommaso, P., & Nahnsen, S. (2020). The nf-core framework for community-curated bioinformatics pipelines. Nature Biotechnology, 38, 276–278.

67. Robinson, M.D., McCarthy, D.J., and Smyth, G.K. (2009). edgeR: A Bioconductor package for differential expression analysis of digital gene expression data. Bioinformatics 26, 139–140.

68. Ritchie, M.E., Phipson, B., Wu, D., Hu, Y., Law, C.W., Shi, W., and Smyth, G.K. (2015). Limma powers differential expression analyses for RNA-sequencing and microarray studies. Nucleic Acids Research 43, e47.

69. 69. Yan L (2025). ggvenn: Draw Venn Diagram by ’ggplot2’. R package version 0.1.10, https://github.com/yanlinlin82/ggvenn

70. 70. Raivo Kolde (2019). pheatmap: Pretty Heatmaps. R package version 1.0.12. https://CRAN.R-project.org/package=pheatmap

71. Xu, S., Hu, E., Cai, Y. Xie, Z., Luo, X., Zhan, L., Tang, W., Wang, Q., Liu, B., Wang, R., Xie, W., Wu, T., Xie, L., Yu, G. (2024). Using clusterProfiler to characterize multiomics data. Nature Protocols 19, 3292–3320.

72. 72. R Core Team (2021). R: A language and environment for statistical computing. R Foundation for Statistical Computing, Vienna, Austria. https://www.R-project.org/.

73. Vonrhein, C., Flensburg, C., Keller, P., Sharff, A., Smart, O., Paciorek, W., Womack, T., and Bricogne, G. (2011). Data processing and analysis with the autoPROC toolbox. Acta Crystallographica Section D: Biological Crystallography 67, 293–302.

74. McCoy, A.J., Grosse-Kunstleve, R.W., Adams, P.D., Winn, M.D., Storoni, L.C., and Read, R.J. (2007). Phaser crystallographic software. Journal of Applied Crystallography 40, 658–674.

75. Emsley, P., Lohkamp, B., Scott, W.G., and Cowtan, K. (2010). Features and development of Coot. Acta Crystallographica Section D: Biological Crystallography 66, 486–501.

76. Afonine, P.V., Grosse-Kunstleve, R.W., Echols, N., Headd, J.J., Moriarty, N.W., Mustyakimov, M., Terwilliger, T.C., Urzhumtsev, A., Zwart, P.H., and Adams, P.D. (2012). Towards automated crystallographic structure refinement with phenix.refine. Acta Crystallographica Section D: Biological Crystallography 68, 352–367.

77. The PyMOL Molecular Graphics System, Version 3.0 Schrödinger, LLC.

78. Pettersen, E.F., Goddard, T.D., Huang, C.C., Couch, G.S., Greenblatt, D.M., Meng, E.C., and Ferrin, T.E. (2004). UCSF Chimera - A visualization system for exploratory research and analysis. Journal of Computational Chemistry 25, 1605–1612.

